# Ancestree: unified likelihood inference of ancestral alleles under supplied or inferred genealogies

**DOI:** 10.64898/2026.09.11.750934

**Authors:** Janek Sendrowski, Thomas Bataillon

**Affiliations:** Bioinformatics Research Center, Aarhus University, Aarhus 8000 DK

**Keywords:** ancestral-state inference, Felsenstein pruning, phylogenetic likelihood, coalescent inference, ancestral recombination graph, site-frequency spectrum, population genetics

## Abstract

Inferring ancestral states—determining, at each polymorphic site, which allele is ancestral and which derived—underpins many downstream population-genetic analyses, from selection scans and the unfolded site-frequency spectrum to demographic inference. However, no existing tool uniformly supports the full range of relevant inputs: plain variant data or ancestral recombination graphs (ARGs), with or without outgroups, while accommodating poly-allelic and recurrently-mutated sites. Here we present `Ancestree`, a likelihood-based engine that unifies these inputs within a single framework and returns full posteriors over the four nucleotide states at every site. It runs in three modes: a *fixed-tree mode* that assumes a single topology across sites and co-infers the per-branch substitution rates by maximum likelihood; an *ARG mode* that reads a different local tree at each site directly from a supplied ancestral recombination graph; and a *local-tree mode* that instead samples those local trees from the genotype data via a pairwise-coalescent HMM, needing no pre-existing ARG. On simulated data, the genealogy-based modes (ARG and local-tree) are more accurate and scale better, and remain robust under outgroup configurations that violate the fixed-tree assumption. Outgroups themselves remain difficult to replace: per-site inference accuracy on ingroup-polymorphic sites is markedly limited without them, and improves substantially with a single outgroup. The hardest sites are those fixed for the derived allele within the ingroup, which carry no within-ingroup signal and so need several sufficiently deep outgroups to recover, yet these are also highly informative downstream, carrying the high-frequency divergence signal on which selection and adaptation analyses often depend. `Ancestree` is available at github.com/Sendrowski/Ancestree.

## 1 Introduction

Knowing which allele is ancestral at a polymorphic site underpins a broad range of population-genetic analyses: selection tests based on the unfolded site-frequency spectrum (SFS) [Fay and Wu, 2000, Zeng et al., 2006], haplotype scans for recent positive selection [Voight et al., 2006], distribution-of-fitness-effects inference [Tataru et al., 2017], estimators of allele age, and predictions of variant deleteriousness all require per-site ancestral-state assignments. Furthermore, ARG-inference methods take them as input [Kelleher et al., 2019, Speidel et al., 2019]. Ancestral calls are what make the unfolded SFS informative about positive selection: derived alleles sit at low frequency under neutrality, so an excess of high-frequency derived variants is a hallmark of selection driving a beneficial allele up faster than drift—a signal visible only once the ancestral state is known. Errors are correspondingly costly: low-frequency variants greatly outnumber high-frequency-derived ones, so misidentifying even 1–2% of them inflates the high-frequency-derived class and biases tests of positive selection. Ancestral states can be inferred from one or more *outgroups*—lineages that diverged before the ingroup and are assumed to carry the ancestral state. Such outgroup evidence can be used to obtain an ancestral-state assignment either by parsimony or, more robustly, by computing the probability that each allele is ancestral under an explicit model of sequence evolution.

A useful reference case is the infinite-sites model (ISM) [Kimura, 1969], which postulates that each site mutates at most once across the genealogy—a simplifying assumption underlying much of population-genetic theory. Under ISM, at a biallelic ingroup site the lone mutation must lie on an ingroup-internal branch, so any outgroup—provided it is free of introgression and distant enough that incomplete lineage sorting (ILS) is unlikely—is fully informative about the ancestral state. Real data violates all three premises: at moderate outgroup depths, ILS nests the outgroup within the ingroup genealogy at a fraction of loci, hybridisation transfers alleles across the ingroup–outgroup split, and recurrent or back-mutations break ISM. Sequencing and alignment errors compound the problem. Even a correctly orthologous outgroup need not carry the ancestral state: a deep shared polymorphism retained across the ingroup–outgroup split, or a mutation on the outgroup’s own branch since that split, can leave its allele derived rather than ancestral (Figure 1). Model-based inference that combines evidence from several outgroups handles these failure modes by emitting a posterior over candidate ancestral states, degrading gracefully on the subset of sites where the trivial outgroup-as-ancestral rule fails, whereas a mis-assignment there propagates as bias into downstream summaries [Baudry and Depaulis, 2003].

**Figure 1:**
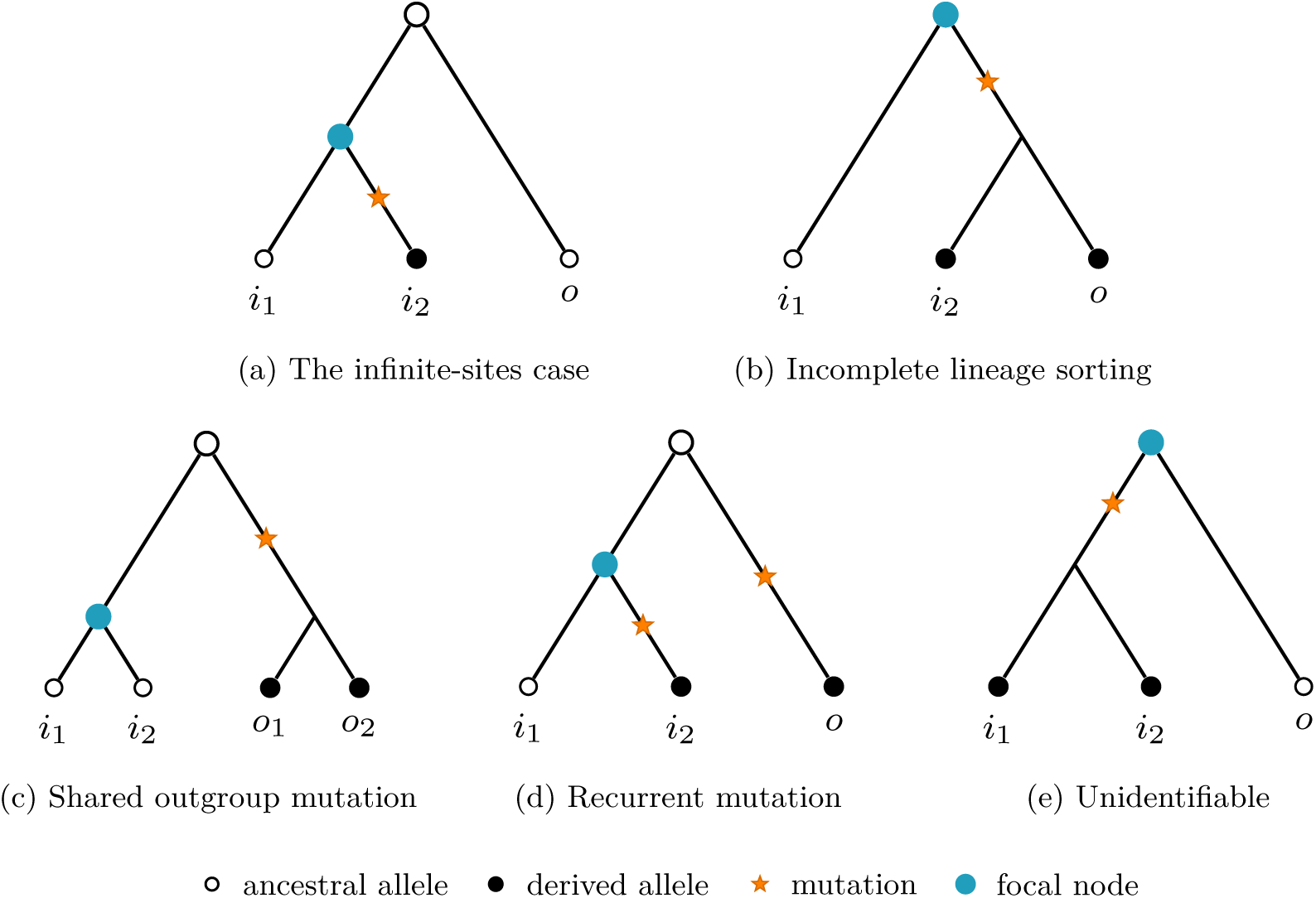
The infinite-sites case, and four ways an outgroup can mislead ancestral-allele inference. The focal node is the one whose ancestral state is inferred. **(a)** Under the infinite-sites model the single mutation falls on an ingroup-internal branch, so the outgroup *o* carries the ancestral state by construction. **(b)** Incomplete lineage sorting nests the outgroup within the ingroup, leaving it less informative about the ancestral state. **(c)** The two outgroups coalesce with each other before joining the ingroup, so the second adds little to the first. **(d)** The same derived allele arises independently on two branches (homoplasy), violating the infinite-sites assumption. **(e)** A mutation subtending all ingroup samples leaves the ancestral state unidentifiable from a single outgroup. A tree-based likelihood approach can quantify this uncertainty exactly and weight the outgroup evidence by its true information content.

### Outgroup-based inference

A line of population-genetic work has progressively refined the outgroup-based polarisation problem from early parsimony-corrected schemes [Eyre-Walker, 1998, Baudry and Depaulis, 2003, Hernandez et al., 2007] into maximum-likelihood frameworks that combine outgroup-tree evidence with ingroup site-frequency information, of which `EST-SFS` [Keightley and Jackson, 2018] is the current standard. `EST-SFS` assumes one tree topology, shared by every site, linking the focal *ingroup* to up to three distantly related outgroups, whose homologous alleles inform on the ingroup’s ancestral state. That topology is fixed in advance, and its per-branch substitution rates are estimated by maximum likelihood across all sites. A second optimisation step then incorporates ingroup allele-frequency information to refine the per-site posteriors. Several practical limitations follow from this design. The assumed topology fails whenever the species tree is misspecified or ILS nests a too-closely-related outgroup inside the ingroup, which biases the assignments and can leave the output overconfident. In addition, usable outgroups are not always available: relict or non-model species may have none, and genomic alignments involving deeply diverged ingroup and outgroup genomes are often unreliable [Bergman and Schierup, 2021]. Furthermore, `EST-SFS`’s kernel is limited to three outgroups and biallelic sites, so larger panels and poly-allelic data are excluded. `EST-SFS` also does not support *divergence* sites (ingroup-fixed for the derived allele): its SFS prior constrains any monomorphic site’s posterior to the ingroup-fixed allele regardless of outgroup evidence. At the sample level, a fixed ingroup sample size is required, so missing genotypes are handled by random subsampling of haplotypes which injects sampling noise into the per-site posterior.

### Genealogy-based inference

A different route to ancestral-allele inference is taken by `PolarBEAR` [Liang and Dutheil, 2026]: it reads the local genealogy directly from an ARG inferred over the ingroup. An ARG encodes the full coalescent history of the sample, with a different local genealogy associated with each non-recombining segment of the genome [Kelleher et al., 2018]; for each polymorphic site, the local (marginal) tree provides the topology and branch lengths that `EST-SFS` would otherwise have to assume from a fixed species tree. This is attractive on two counts. The ingroup’s coalescent structure is informative about the ancestral state at a site—segregating alleles whose carriers form a tight clade are more likely to be derived than those scattered across the tree—and reading the local tree directly from the ARG avoids the tree misspecification described above. The method also dispenses with outgroups entirely, extending ancestral-state inference to species for which outgroup alignments are unreliable or unavailable. The ingroup genealogy does not, however, carry ancestral-state signal at every site: sites where the ingroup is monomorphic for the derived allele cannot be polarised from the ingroup alone, and high-frequency derived alleles more generally provide weaker signal than singletons. `PolarBEAR` thus focuses on biallelic sites whose ancestral state can be resolved with confidence given the local tree. More fundamentally, the genealogy route presupposes an inferred ARG, yet most ARG-inference methods themselves require ancestral states as input [Kelleher et al., 2019, Speidel et al., 2019]—a circularity that constrains where genealogy-based ancestral-state inference can be applied. For this reason Liang and Dutheil [2026] focus on ARGs inferred without ancestral alleles, particularly those from HMM-based methods that leverage heterozygosity. This sidesteps the circularity, but makes each analysis conditional on a separately inferred ARG.

### Ancestral-sequence reconstruction

A separate literature on phylogenetic ancestral-sequence reconstruction (ASR) reconstructs the sequence at internal nodes of a known phylogeny—`PAML` [Yang, 2007], `FastML` [Ashkenazy et al., 2012], and `IQ-TREE` [Minh et al., 2020], building on Yang et al. [1995], Pupko et al. [2000]—for a few well-separated taxa at deep divergence. These methods assume one sequence per taxon, a single tree for the whole alignment, and dense phylogenetic coverage. That coverage condition is often unmet in practice, and with one sequence per taxon ASR disregards within-species polymorphism, the signal a population sample carries about the ancestral state. Population-genetic ancestral-allele inference—`EST-SFS`, `PolarBEAR`, and the present work—instead capitalises on large within-species samples at shallow coalescent depth. The two lines of work share the same sequence-evolution likelihood kernel and DNA substitution models, and the rest of this paper focuses primarily on the population-genetic regime, where the genealogy changes along the genome.

### This work

Here, we present `Ancestree`, a likelihood-based engine that unifies both approaches—outgroup-based polarisation (`EST-SFS`) and genealogy-based polarisation (`PolarBEAR`)—behind a single kernel (Section 2.1). It is exposed through three inference modes—*fixed-tree mode*, *ARG mode*, and *local-tree mode*—that share this kernel and differ only in where the tree it scores comes from. Fixed-tree mode generalises the `EST-SFS`-style outgroup-ladder approach to an arbitrary number of outgroups and alleles per site, while also supporting sites that are monomorphic as well as polymorphic within the ingroup. ARG mode mirrors `PolarBEAR` in its input (a `tskit` ARG), exposing a per-state posterior at every polymorphic site. The two modes each carry a structural cost: fixed-tree mode needs outgroups and assumes one fixed species topology for every site; ARG mode treats outgroups as optional and assumes no fixed topology, reading a fresh local tree at each site, but presupposes an inferred ARG. For this reason, we also introduce local-tree mode (Section 2.4), which samples a dated local genealogy per window directly from genotypes via a pairwise-coalescent HMM, then applies the ARG-mode kernel to it, so the per-site, ILS-aware robustness of reading the genealogy becomes available from plain variant data, requiring no pre-existing ARG. Although inferred local trees are only an approximation of the true genealogy, the mode proves robust in practice, tracking ARG mode on the true genealogy closely across benchmark scenarios (Section 3.2). Even though ARG and local-tree mode do not require outgroups, outgroups remain highly informative where available: they attach as extra tips to the local tree, sharpening the posteriors at ingroup-polymorphic sites and supplying the evidence at divergence sites and where the ingroup genealogy is uninformative.

We rely on simulated and real data to validate `Ancestree` against both `EST-SFS` and `PolarBEAR` in their respective regimes, and compare the three modes across a range of scenarios (Section 3). `Ancestree` exposes a native Python API and a command-line interface, and supports VCF [Pedersen and Quinlan, 2017], Zarr [Czech et al., 2025] and `tskit` [Kelleher et al., 2018] ARG data on both input and output. It is extensively documented, unit-tested, and installable from PyPI and conda-forge.

## 2 Methods

Every inference mode in `Ancestree` shares one underlying computation: the Felsenstein pruning likelihood of a site’s observed-allele configuration on a tree under a prior (Section 2.1). The modes differ only in where that tree comes from—a local genealogy read from a supplied ARG (Section 2.2), a fixed outgroup-ladder tree whose branch lengths are fitted by maximum likelihood (Section 2.3), or a local tree inferred from genotypes alone (Section 2.4)—and they draw on a common set of components: **Priors** and **Substitution models**.

### 2.1 The Felsenstein Pruning Kernel

Given a rooted tree with branch lengths and a per-site assignment of alleles to tip samples, Felsenstein’s pruning algorithm [Felsenstein, 1981] returns the likelihood of the observed tip pattern conditional on each possible state at the root.

#### Setup

Let *T* be a rooted tree (Figure 2) with node set *V*, root *r ∈ V*, and branch lengths *t_v_* on the edge connecting each non-root node *v* to its parent. Polytomies (nodes with more than two children) are permitted and require no special treatment. Let *S* = {*A, C, G, T}* index the four nucleotide states; *S* = |S| = 4. Substitutions along branches are governed by a continuous-time Markov chain with instantaneous rate matrix **Q ∊** ℝ*^S^*^×^*^S^*. The probability of transitioning from state *i* to state *j* along a branch of length *t* is the (*i, j*) entry of the transition matrix

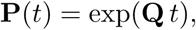

**Figure 2:**
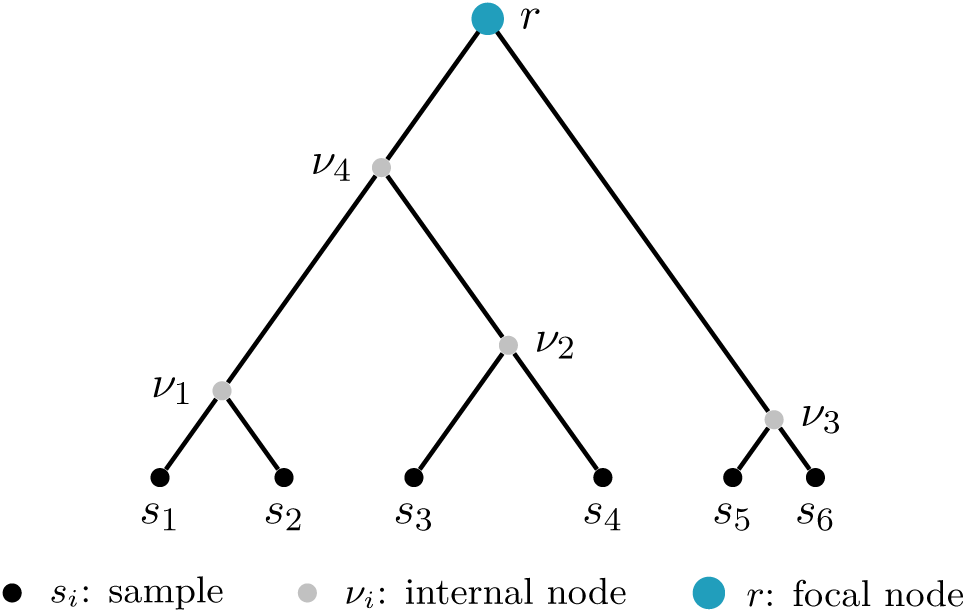
**A tree** with six samples *s*_1_*,..., s*_6_ and root *r*. The likelihood kernel evaluates each polymorphic site against the local tree covering its position, summing over candidate root states under a prior *π*. All tips enter symmetrically; the distinction between ingroup and outgroup enters only through the choice of focal node, the node whose ancestral state is reported, marked here in colour. Here it corresponds to the tree’s own root; with outgroups in the panel it is typically the ingroup MRCA, an interior node of the same tree.

with **Q** supplied by **Substitution models**.

A site is an observation **x** = (*x_u_*)*_u_*_∈*V*_ over the tip set *V*_tip_ ⊆ *V*, where each entry *x_u_* ∊ {∅} = {*A, C, G, T,* ∅} is the allele observed at tip *u*, with ∅ marking a missing genotype.

#### Recursion

For each node *v ∊ V*, we recursively compute a vector ***ℓ****_v_ ∊* ℝ*^S^* of *partial likelihoods*, with ***ℓ****_v_*(*s*) the probability of the observed tip pattern in the subtree rooted at *v*, conditional on *v* being in state *s*. At each tip *u ∊ V*_tip_, the partial likelihood is a one-hot indicator of the observed allele,

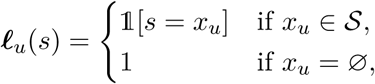

where 1[·] is the indicator function, equal to 1 when its argument holds and 0 otherwise, so the tip places all mass on its observed allele *x_u_* while missing data (*x_u_* = ∅) contributes a 1, thereby leaving the likelihood function unchanged. For an internal node *v* with children *C*(*v*), the partials are obtained by traversing the tree in post-order (every node visited only after all of its descendants, so that ***ℓ****_c_* is already available for each child *c ∊ C*(*v*)) and combining each child’s contribution through its incoming branch of length *t_c_*:

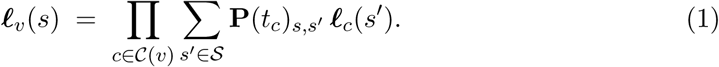

The factor **P**(*t_c_*)*_s,s_′* is the probability of a transition from state *s* at *v* to state *s*^′^ at its child *c*, along the branch of length *t_c_* joining them, so each inner sum is the probability of the data below child *c* given that the parent state is *s*; the outer product reflects the conditional independence of the subtrees rooted at the children of *v*, given *v*’s state. The name *pruning* reflects what this post-order pass avoids: a naive enumeration of all joint internal-node state assignments, which is exponential in tree size. Each subtree’s contribution to its parent is instead summarised in the single length-*S* vector ***ℓ****_v_* and the subtree discarded thereafter.

The recursion terminates at the root *r*, yielding ***ℓ****_r_ ∊* ℝ*^S^*, the per-state (conditional) likelihood of the full observed site configuration. The log-likelihood of the data conditional on root state *s* is then log ***ℓ****_r_*(*s*). Under a prior *π* on the root state, the posterior probability that the root is in state *s* given the observed site **x** and the tree *T* follows from Bayes’ theorem as

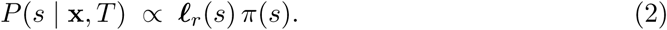

The kernel computes this marginal posterior by the sum-product recursion above. Note that the alternative max-product route, used by `PolarBEAR` [Liang and Dutheil, 2026], answers a slightly different question and is discussed in Appendix A.3. Worked examples of how the posterior responds to the observed tip pattern, from a near-certain assignment to a genuine tie, are given in Appendix A.2, alongside a gallery of tree and site configurations (Appendix A.1).

#### Re-rooting

Equation 2 is expressed at the root, since that is where the recursion terminates, whereas the focal node is in general an interior node of the tree at hand, so the tree is re-rooted there. The substitution models are time-reversible [Squartini and Arndt, 2008], so the likelihood conditioned on a node’s state is the same whichever node the recursion is rooted at, leaving every branch length as it is. This is the invariance behind Felsenstein’s pulley principle [Felsenstein, 1981], and the basis on which ancestral-sequence reconstruction reports marginals at interior nodes [Yang et al., 1995, Yang, 2007, Ashkenazy et al., 2012]. A non-root focal node therefore costs one re-rooting and leaves Equation 2 unchanged in form, with *π* read as the prior at the focal node itself. The construction extends to a point part-way along a branch, so the readout position is continuously adjustable between the two anchors (Appendix C.1). Nothing requires the focal node to be a population’s ancestor, so the kernel annotates a species-level tree in the same way as a population panel. Where a single genome represents each species and no population-level variation is available, the ingroup designation only defines the clade at whose MRCA the posterior is read.

#### Priors

The likelihood kernel returns log *P* (**x** *r* = *s, T*) for every candidate state *s* at the focal node, and turning this into a posterior requires a prior *π*(*s*) over that state. The `StationaryPrior` takes *π*(*s*) to be the base composition supplied to the kernel, and the uniform 1*/S* where none is given; under `F81, HKY` and `GTR` this coincides with *π***_Q_**(*s*), the stationary distribution of the substitution model. The uniform case is not a separate prior but this same one constructed without a base composition, so a run that supplies none is already using it. Under the uniform default, the per-state likelihood ***ℓ****_r_* is itself the posterior up to normalisation. The symmetric models (JC69, K2) have a uniform *π***_Q_** whatever the composition, so an empirical composition enters their inference through this prior alone. This is the natural prior in ARG mode, where the local-tree topology and branch lengths already carry the ingroup coalescent signal, leaving the prior to express what is known about the root state before the data are observed.

#### Recurrent substitution

The kernel evaluates **P**(*t*) = exp(**Q***t*) exactly, so any number of substitutions per branch, including reversions to an earlier state, is admitted. `PolarBEAR` [Liang and Dutheil, 2026] instead retains only the zero- and one-mutation terms and `EST-SFS` [Keightley and Jackson, 2018] additionally the two-mutation term of a truncated Poisson series. The difference is negligible under low mutation rates, where recurrence is rare, and matters only on hypermutable sites where back-substitution is appreciable (Appendix A.3). In the opposite direction, the *µ →* 0 limit reduces the kernel to maximum parsimony: each additional substitution carries a vanishing *O*(*µ*) weight, so the likelihood concentrates on the fewest-change history. Low *µ* with no homoplasy is the infinite-sites regime, a single mutation per site, where parsimony is itself correct.

#### Poly-allelic sites

Sites with more than two alleles at the ingroup tips require no special treatment in the likelihood kernel: the tip partial ***ℓ****_u_* is a one-hot indicator over the model’s full state set regardless of how many distinct alleles the site happens to carry, and the post-order combine step operates over the same *S*-dimensional alphabet at every node. The only special case is the adaptive weight, whose per-bin parameterisation *ω_i_* is biallelic by construction; for *k* ≥ 3 alleles it falls back to the Kingman baseline.

#### Substitution models

`Ancestree` provides a nested family of `SubstitutionModel` implementations, each supplying the rate matrix **Q** the kernel exponentiates: JC69 [Jukes and Cantor, 1969], a single rate with a uniform stationary distribution; K2 [Kimura, 1980], adding a transition/transversion ratio *κ*; `F81` [Felsenstein, 1981], making off-diagonal rates proportional to empirical base frequencies *ϕ*; `HKY` [Hasegawa et al., 1985], combining K2’s *κ* with F81’s base frequencies; and `GTR` [Tavaré, 1986], the general time-reversible model with five free parameters. Each reduces to a simpler member at special parameter values (e.g. HKY → F81 at *κ* = 1, and to JC69 at uniform *π*; GTR → HKY). The free parameters (*κ*, the GTR rates) can be supplied directly or jointly fitted with the branch lengths in `FixedTreeInference`. For reference, `EST-SFS` [Keightley and Jackson, 2018] implements the Jukes–Cantor, Kimura two-parameter, and six-rate models, and `PolarBEAR` [Liang and Dutheil, 2026] assumes the Jukes–Cantor model. The effect of misspecifying the substitution model is quantified in Appendix C.4.

### 2.2 ARG Inference

`ARGBasedInference` applies the likelihood kernel directly to an externally supplied ancestral recombination graph, reading the marginal tree at each site and scoring the posterior at that site’s focal node on the given genealogy. The mode uses the substitution model’s stationary root prior (**Priors**), and outgroups, where available, attach as extra tips. Its accuracy is bounded by the input graph: because the graph is fixed, ancestral-allele errors embedded in its construction cannot be corrected here, so it is best paired with an ARG inferred without conditioning on the ancestral state (Appendix B.5). Segments that have not fully coalesced can carry several roots. Where a focal node is given the kernel is run on the root subtending it; otherwise it is run on each coalesced root’s subtree in turn, and the posteriors averaged under a uniform prior over those roots. The mutation rate *µ* may be a single value or vary along the genome, each local tree then taking the interval-weighted mean over its own span. Inferred ARGs are benchmarked in Section 3.3.

### 2.3 Fixed-Tree Inference

Given genotypes for the ingroup and one or more outgroup haplotypes, but no inferred genealogy, one approach is to assume a fixed topology for how the ingroup and outgroups are related [Keightley and Jackson, 2018]. This approximation is reasonable when all outgroups are sufficiently diverged from the ingroup to be informative for ancestral-state inference (Appendix C.12). We refer to such a tree as an *outgroup-ladder tree* (`OutgroupLadderTree`).

#### The outgroup-ladder tree

The *n* ingroup haplotypes collapse to a single polytomy at the ingroup most recent common ancestor (MRCA), and the outgroups join the ingroup lineage in a nested ladder ordered by decreasing relatedness: each outgroup is assumed to coalesce with the ingroup lineage before it coalesces with any other outgroup (Figure 3). For *n*_out_ outgroups, the tree carries 2*n*_out_ − 1 free branch-length parameters 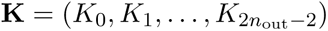, mapped one-to-one to the ladder edges shown in Figure 3, each in units of the expected number of substitutions per site, matching `EST-SFS`. The deepest branch, 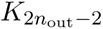, stretches across the species-tree root *r*, whose position along it is unidentifiable. The collapsed ingroup haplotypes carry no branches, entering instead through an SFS-based weight at I, described below.

**Figure 3:**
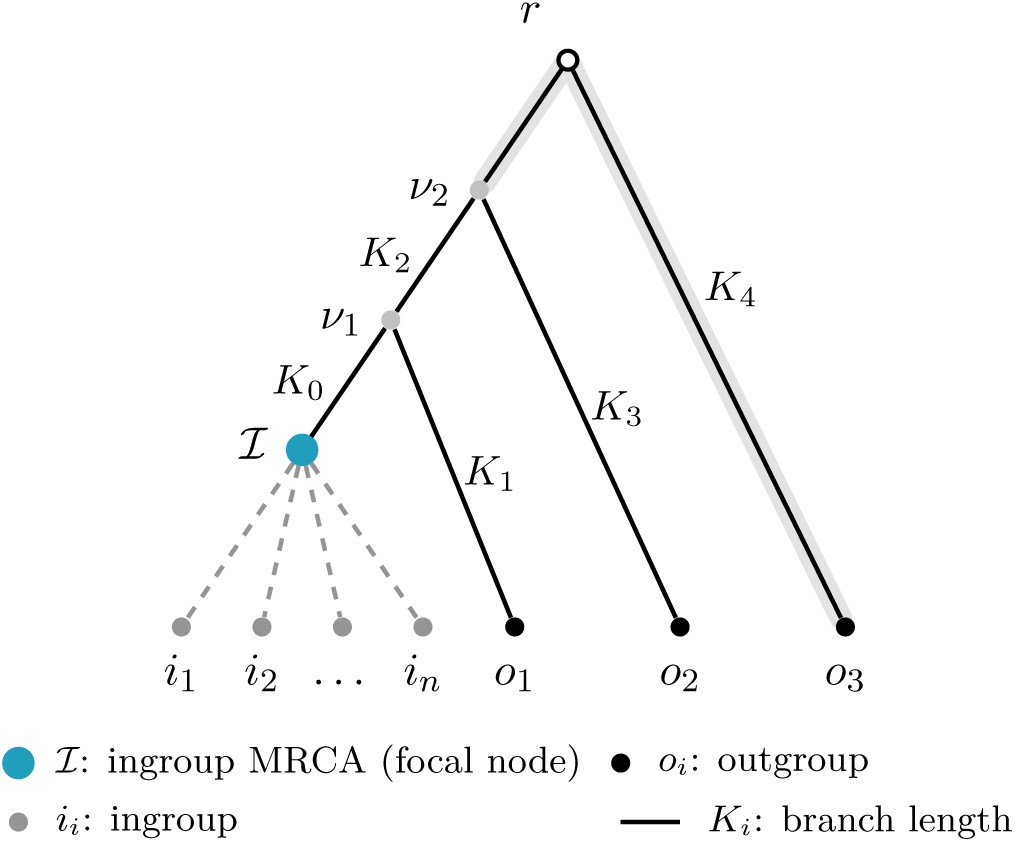
**Outgroup-ladder tree** with *n*_out_ = 3 outgroups. The outgroups are ordered by decreasing relatedness: the closest, *o*_1_, joins most recently, and the most distant, *o*_3_, sits at the end of the deepest branch *K*_4_. The ingroup haplotypes *i*_1_*,..., i_n_* (grey, dashed) collapse to a polytomy at and carry no branches, entering only through the per-site ingroup weight. The 2*n*_out_ − 1 = 5 branch lengths (*K*_0_*,..., K*_4_) are estimated by maximum likelihood. The species-tree root *r* is shown hollow at the apex, but it is not separately estimated: under the time-reversible model its position along the deepest branch is unidentifiable. The single length *K*_4_ therefore spans both segments either side of it, as the shaded band indicates.

#### Fitting the branch rates

`FixedTreeInference` estimates **K** by maximum likelihood, then emits per-site posteriors at the fitted rates. The marginal log-likelihood is a single weighted sum over the distinct site configurations the data realise, polymorphic and monomorphic alike:

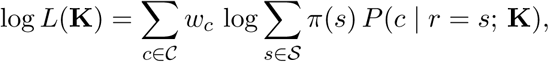

where *C* is the set of distinct site configurations, *w_c_* is the expected number of sites with configuration *c*, *s* ranges over the candidate alleles *S* at the root *r*, *π*(*s*) is the prior on *s*, and *P* (*c r* = *s*; **K**) is the probability of configuration *c* given root state *s*, evaluated by the likelihood kernel against the outgroup-ladder tree. A configuration *c* = (**n**, **o**) pairs the ingroup allele counts **n** = (*n_A_, n_C_, n_G_, n_T_*) with the alleles **o** = (*o*_1_*,..., o_n_*_out_) observed at the outgroup tips. Following Keightley and Jackson [2018], it captures everything a site’s likelihood depends on, so sites sharing a configuration collapse into one weighted term. In practice, the likelihood is maximised by multi-start L-BFGS-B with log-uniformly sampled initial rates.

#### Sub-sampling

Missing data and varying ploidy leave a site with *n*_obs_ ≤ *n* called ingroup haplotypes. The ingroup tuple **n** is therefore projected onto a common subsample of size *n*_sub_. Rather than draw one random sub-sample of size *n*_sub_ from the *n*_obs_ observations (which would assign the site to a single bin and inject sampling noise), a site with observed minor-allele count *k* has its contribution distributed across bins *j* ∈ {0*,..., n*_sub_} proportionally to the hypergeometric probability mass function

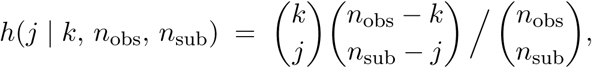

the mass a site with *k* minor-allele copies among *n*_obs_ called haplotypes contributes to bin *j* of the sub-sample of size *n*_sub_.

#### Monomorphic sites

A monomorphic site is a configuration in which every tip carries the same base. Such sites are absent from variant-only input and are supplied as per-base counts over the invariant positions. They carry most of the weight and so pin the total tree length Σ*_i_ K_i_*, while the polymorphic configurations split that length across branches. Poly-allelic ingroup polymorphisms are excluded from the branch-rate maximum-likelihood fit but still scored at inference time on the fitted tree, with no measurable effect on the recovered rates. Computationally, the fit streams the variant input rather than holding it in memory all at once. The number of sites the fit requires for stable rate estimates is examined in Appendix C.5.

#### Ingroup weights

Collapsing the ingroup to a polytomy discards its topological signal. What remains is the ingroup allele counts **n**, which enter as

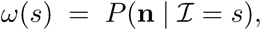

the likelihood of those counts given the state *s* at the ingroup most recent common ancestor *I*. The weight enters at *I* inside the pruning recursion, so it propagates up the branch to the focal node and fades with distance (Section C.1). The posterior is therefore proportional to the root-state prior *π*(*s*) (**Priors**) times the kernel evaluated with *ω* at *I*. Such a weight can improve the estimates, especially when few or no outgroups are present (Appendix C.3). `Ancestree` supplies two ingroup weights.

#### Kingman weight (KingmanIngroupWeight)

Under a neutral Kingman coalescent, that is, in a panmictic population at mutation–drift equilibrium, a derived allele at count *i* in an ingroup of *n* samples contributes to the expected SFS with weight proportional to 1*/i* [Watterson, 1975], so conditional on observing a biallelic site with minor-allele count *i*, the major allele (count *n − i*) is the ancestral one with probability (1*/i*)*/*(1*/i* + 1*/*(*n i*)) = (*n − i*)) = (*n* −*i*)*/n*. For a site with observed ingroup allele counts {*a*_1_ : *n*_1_*,..., a_k_* : *n_k_*} (allele *a_j_* observed *n_j_* times), summing to *n*_obs_, the number of called ingroup genotypes at the site, the weight at candidate state *a_j_* is thus

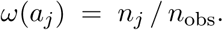

The same count-proportional formula extends naturally to *k >* 2 segregating alleles, and to ingroup-monomorphic sites, where all weight falls on the single observed allele, leaving the outgroup evidence to decide whether the site is fixed ancestral or fixed derived.

#### Adaptive weight (AdaptiveIngroupWeight)

The adaptive weight generalises the Kingman weight, replacing its closed form with a free parameter *ω_i_* per folded SFS class *i*. For each class *i*, the weight asks the data what fraction of sites with *i* copies of the minor allele have the major allele as the ancestral one, rather than imposing the neutral Kingman expectation. `EST-SFS` [Keightley and Jackson, 2018] adopts the same data-driven approach, estimating these per-frequency weights rather than fixing them at the neutral expectation. Under a neutral Kingman coalescent the answer is exactly (*n*_sub_ − *i*)*/n*_sub_, so the adaptive weight converges to Kingman in the neutral large-data limit. The adaptive weight is therefore useful where the SFS is expected to deviate from neutrality—under positive selection, sustained demographic distortion, or ascertainment bias. On most empirical datasets the fitted *ω_i_* closely resemble the Kingman weights and become unreliable on small ones, so the adaptive weight is best reserved for larger datasets (≳ 20 sites per SFS class; Appendix C.5).

The *ω_i_* are fitted in a second stage, after the tree-rate MLE has been computed, by maximising the per-bin log-likelihood

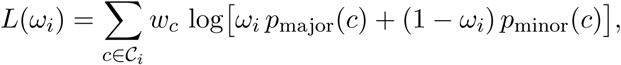

Here *i ∊* {0*,..., n*_sub_} indexes the sub-sample classes, *C_i_* ⊆ *C* is the set of configurations with *i* minor-allele copies, *w_c_* is the mass the sub-sampling assigns to configuration *c*, and *p*_major_(*c*) and *p*_minor_(*c*) are the likelihood-kernel values (Section 2.1, Eq. 1) of *c* under the major and minor allele as root, evaluated at the fitted tree. The bracket mixes two hypotheses, the major allele ancestral with weight *ω_i_* and the minor allele ancestral with weight 1 − *ω_i_*, so maximising over *ω_i_* asks how the data split class *i* between them. The boundary classes are fixed at *ω*_0_ = 1 and 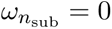, where only one allele is present. Folding 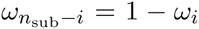 pairs each class with its mirror, leaving the free parameters *i ∊* {1*,...,* ⌈*n*_sub_*/*2⌉ − 1}, which are maximised independently; for even *n*_sub_ the self-paired middle class is fixed at 1*/*2. The per-bin parameterisation is biallelic by construction; poly-allelic sites and sites with *n*_obs_ *< n*_sub_ fall back to the Kingman baseline.

### 2.4 Local-Tree Inference

ARG mode needs a genealogy, yet most ARG-inference pipelines need ancestral states to build one. `LocalTreeInference` breaks this loop, inferring a dated local genealogy from genotypes alone and passing it to the same kernel, ideally with outgroups attached. Beyond the genotypes, the per-site mutation and recombination rates (*µ, ρ*) are the only inputs required, each supplied either as a single constant or as a rate map. Local trees are built per window in two stages. First, for each pair of haplotypes (*a, b*) a time-discretised pairwise-coalescent HMM—a PSMC-style model [Li and Durbin, 2011] with the pairwise TMRCA as the hidden state—infers the local coalescence time along the genome. Second, those times are clustered locally into dated trees, either once from their posterior means or over an ensemble of trees sampled from the same posterior, which the per-site posterior is then marginalised over.

#### The pairwise HMM

In HMM terms, two haplotypes are compared in fixed blocks of *B* base pairs, the observation at block *k* being their difference count *c_k_*, the number of sites at which they differ there, and the hidden state *z_k_* the block’s discretised TMRCA bin, so that *z_k_* = *i* means block *k*’s TMRCA lies in bin *i*. The emission probability is *e_i_*(*c_k_*) = *P* (*c_k_ z_k_* = *i*) and the transition probability from bin *i* to bin *j* between consecutive blocks is *A* = *P* (*z* = *j z* = *i*). The TMRCA is discretised into *T* bins on a log-spaced grid with edges *u*_0_ *< … < u_T_* and geometric bin midpoints 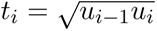, spanning a per-dataset range derived from the observed per-block divergences. The coalescent prior *ψ*is per-pair, giving the pair’s TMRCA distribution over the bins before any block is observed. Under a constant-size coalescent, the pairwise TMRCA of haplotypes (*a, b*) is exponentially distributed with mean their genome-average coalescence time 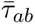 (estimated from the pair’s mean per-block divergence), so *ψ_j_* is the exponential’s mass in bin *j*,

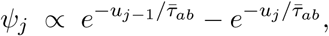

normalised to sum to one over the *T* bins. The per-block TMRCA posteriors are computed by the forward–backward recursion over this HMM. Because each window’s TM-RCA is a posterior mean over the bins rather than the single most-probable bin, the estimate interpolates between grid points rather than being confined to a single grid value, so accuracy is insensitive to the bin count *T*. The emission treats a block’s difference count as Poisson-distributed—that is, *c_k_* is Poisson with mean the bin’s expected divergence,

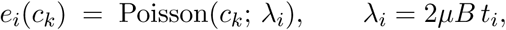

where *µ* is the per-site per-generation mutation rate, *t_i_* the TMRCA of bin *i*, and *B* the block length in base pairs. Missing genotypes and inaccessible spans shrink *B* to the sites at which both haplotypes are called, so that uncalled sites do not deflate the pair’s inferred coalescence time. A block with no called sites takes an emission constant across the *T* bins rather than being read as a run of zero differences. The pair’s TMRCA remains constant only within a non-recombining segment; a recombination breakpoint between blocks begins a new segment whose lineages re-coalesce independently, with a TMRCA drawn independently from the prior. Recombination enters the model as a reset: between two consecutive blocks the hidden state is redrawn, with probability 1 − *r_i_*, from a reset distribution *ν* over the same *T* bins, where 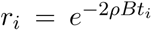 is the no-recombination probability at TMRCA *t_i_* across one block and *ρ* is the per-site recombination rate. A recombination map or an uncovered gap replaces *ρB* with that step’s recombination distance. This gives the per-block transition probability

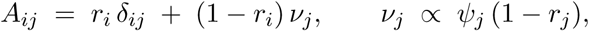

with *δ_ij_* the Kronecker delta and *i, j* indexing the *T* bins, each row summing to one. Contigs and declared breakpoints are severed by taking *ρB* → ∞. Stationarity determines the reset distribution *ν.* Imposing *ψA* = *ψ* gives 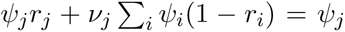 whose unique solution is the *ν* above. Every block then carries the same marginal TMRCA distribution *ψ*, as required of a coalescent process that is stationary along the genome. Because each transition row combines a stay-probability with a reset to the prior *ψ*, the forward recursion simplifies:

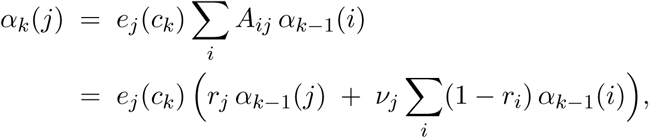

where the forward probability *α_k_*(*j*) = *P* (*c*_1:_*_k_, z_k_* = *j*) is that of the counts through block *k* together with *z_k_* = *j*. The bracket collects the only two ways to be in bin *j* at block *k*: remaining there from block *k* −1 (no recombination, probability *r_j_*) or resetting into it (probability *ν_j_*) from the reset mass Σ*_i_*(1 − *r_i_*) *α_k_*_−1_(*i*) Because that mass is a single scalar reused for every *j*, the step avoids the full *T × T* matrix–vector product. The backward probability *β* collapses by the same structure,

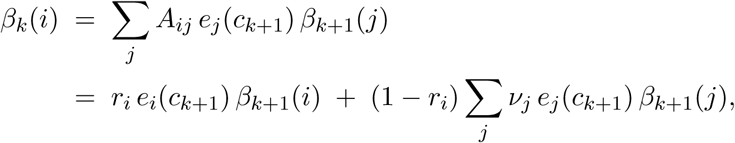

where the backward probability 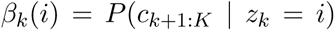 is that of the remaining counts given *z_k_* = *i*, and the sum 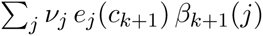 is again a single scalar shared Combining the two then gives the per-block smoothed posterior *γ_k_*(*i*) = *P* (*z_k_* = *i | c*_1:_*_K_*) ∝ *α_k_*(*i*) *β_k_*(*i*), the probability that block *k*’s TMRCA falls in bin *i* given the pair’s entire difference-count sequence. Its expectation over the geometric bin midpoints *t_i_*, 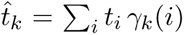, is the posterior-mean TMRCA for block *k*. A *window* is a contiguous run of blocks forming the genomic span assigned one local tree; averaging 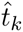 over its blocks gives a single pairwise coalescence time 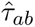 for haplotypes (*a, b*). The HMM itself runs over the whole chunk as one chain, windows entering only at this averaging step. In practice, to keep memory bounded, the genome is processed in chunks. To avoid edge effects at the cuts, each chunk is extended by a margin that is decoded and discarded, so the HMM has decorrelated from the cut before the first retained block.

#### Clustering into local trees

Within each window the pairwise-time matrix (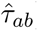) is then clustered by UPGMA [Sokal and Michener, 1958] into a single ultrametric (every tip equidistant from the root), dated tree: at each step it merges the two clusters *C_i_, C_j_* (sets of haplotypes, with |*C*| a cluster’s size) with the smallest average-linkage time

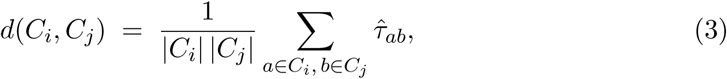

dating the new node to that time and updating distances to the merged cluster as the size-weighted mean. The Felsenstein likelihood kernel (Section 2.1) then scores each site against its window’s trees.

#### Sampling genealogies

The construction above gives each window a single genealogy, built from the posterior-mean times 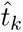, so the Felsenstein kernel is conditioned on a point estimate treated as known, and the spread of *γ_k_* about its mean is discarded. `LocalTreeInference` can instead marginalise the genealogy out, replacing that single tree by an ensemble of *M* genealogies drawn from the HMM posterior (*M* = 128 throughout this paper). A member is drawn by forward filtering and backward sampling [Chib, 1996, Rasmussen et al., 2014]. The forward recursion above is run once per pair and its probabilities *α_k_*(·) retained for every block rather than only combined with *β*. A bin path is then drawn backwards along the genome, starting from the last of the *K* blocks, whose bin is sampled from *α_K_* normalised over the *T* bins. For each *k < K* the bin at block *k* then follows from its conditional given the bin *j* already drawn at block *k* + 1, that is,

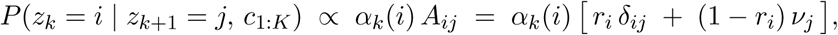

where *α_k_*(*i*) carries the evidence to the left of block *k* and *A_ij_* one transition step to the right. By the Markov property, *z_k_* is independent of the counts *c_k_*_+2:_*_K_* further right once *z_k_*_+1_ is known, so the backward probability is not required. Each member is a joint draw of the pair’s entire TMRCA path along the genome, carrying the between-block correlation the transition model implies, rather than a sequence of independent per-block draws. The drawn bin path *z*_1:_*_K_* assigns only one of the *T* discrete times per block, so a continuous TMRCA is obtained by drawing from the pair’s coalescent prior truncated to the sampled bin, Exp(1*/t̅_ab_*) with *t̅_ab_* the pair’s mean TMRCA over the whole chunk.

#### Ensemble averaging

Member *m* contributes, per window, the block-averaged pairwise times 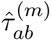, which UPGMA (Eq. 3) clusters into a dated genealogy *G_m_*. Every member is then scored by the Felsenstein kernel and the site likelihoods are averaged,

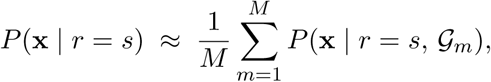

a Monte Carlo estimate of the site likelihood with the genealogy marginalised out. The root-state prior (**Priors**) is then applied to it as before. Because the average is taken over likelihoods rather than over per-member posteriors, each draw is weighted by how well its genealogy explains the site, and a member that explains it poorly contributes little. Haplotype pairs are sampled independently of one another and reconciled only by the subsequent clustering, so the ensemble draws from the product of the per-pair posteriors rather than from a joint posterior over genealogies. The accuracy of local-tree mode, its sensitivity to the window size, rate misspecification and phasing error, and the fidelity of its genealogies are examined in Section 3.4.

## 3 Results

We begin by confirming that `Ancestree` reproduces the tools it subsumes, `EST-SFS` and `PolarBEAR`, on simulated and on real data (Section 3.1). We then compare `Ancestree`’s three inference modes across outgroup counts, scenarios and site classes (Section 3.2), examining local-tree mode in detail (Section 3.4). Ancestral alleles inferred on genealogies from dedicated ARG-inference tools are then scored against those inferred on the true ARG (Section 3.3), and we show how collapsing the per-site posteriors biases the unfolded site-frequency spectrum derived from them (Section 3.5). Further benchmarks and worked examples are collected in Appendix A.

### Choice of focal node

Throughout the benchmarks the posterior is evaluated at the focal node (Section 2.1) determined by the site’s pattern of polymorphism: the ingroup MRCA *I* at sites polymorphic within the ingroup, and the panel root at sites monomorphic within the ingroup, whose shared allele may be ancestral or derived. A polymorphism within the ingroup is a statement about the state at *I*, and reading it at a deeper node would render the posterior sensitive to substitutions on the branch between the two. Where the ingroup is monomorphic, by contrast, the shared allele fixes the state at *I* while leaving open whether that state is ancestral, a question only the outgroup evidence at the deeper node resolves. Appendix C.1 varies the focal node continuously between the two anchors and quantifies the accuracy attained at intermediate positions.

### Common simulation setup

Most simulations use `msprime` ancestral simulations under the JC69 substitution model (**Substitution models**) unless stated otherwise [Baumdicker et al., 2022]. Effective population sizes are *N_e_* = 3 × 10^4^ for the ingroup and ancestral populations; the recombination rate is *r* = 10^−8^ and the base mutation rate *µ* = 1.25 × 10^−8^ per generation per site. Each outgroup population contributes a single haploid sample, so its *N_e_* does not enter the simulation through within-population coalescence—only the reported outgroup-to-ingroup split times matter.

### Metrics

Every score compares a site’s inferred posterior *p* over the alleles *a* against a reference *q*, either the simulated truth or another tool’s output, at one of two levels: the MAP estimate, or the full posterior. *MAP agreement* is the fraction of sites at which the two put their largest mass on the same allele, termed *MAP recovery* where the reference is the truth. At the posterior level, the *Brier score*

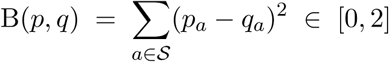

is averaged over sites, and is 0 for identical posteriors and 2 where each is confident on a different allele, so lower is better. Against the simulated truth *q* places all its mass on the true allele. Unlike MAP agreement it reads the whole posterior, and in expectation is lowest when the reported posterior is the one the data support, penalising confident errors more heavily.

### 3.1 Agreement with Existing Tools

Fixed-tree mode should reproduce `EST-SFS`, and ARG mode `PolarBEAR`, wherever their respective modelling assumptions hold. On a 40-haplotype ingroup simulated with three nested outgroups, satisfying the fixed outgroup-ladder tree `EST-SFS` assumes, `FixedTreeInference` recovers `EST-SFS`’s branch rates to within 1–3% and matches its per-site estimates with zero MAP disagreements, their mean Brier scores agreeing to four decimal places (Appendix B.1). Given the same ARG, `ARGBasedInference` reproduces `PolarBEAR`’s per-site MAP estimates exactly on the ingroup-biallelic sites, their mean Brier scores agreeing to six decimal places (Appendix B.2; see also Appendix A.3).

#### Real data

On real data no ground truth exists, so we instead measure inter-tool agreement on the 1000 Genomes MSL (Mende in Sierra Leone) chromosome-1 panel [1000 Genomes Project Consortium, 2015] that the `PolarBEAR` study uses: 174 haplotypes, with orangutan and macaque as outgroups (Appendix B.3). Each mode receives the evidence its reference tool uses. `PolarBEAR` does not accept outgroups, so both genealogy-based modes are run on the ingroup alone, ARG mode reading the same `gamma-SMC` [Schweiger and Durbin, 2023] ARG that the `PolarBEAR` study reconstructs for this panel, and local-tree mode inferring its genealogies from the ingroup genotypes. Fixed-tree mode, like `EST-SFS`, fits the outgroup ladder to the two outgroups. The equivalences seen in simulation hold here too, with ARG mode reproducing `PolarBEAR` on its informative sites and fixed-tree mode closely tracking `EST-SFS`. Local-tree mode agrees closely with `PolarBEAR` as well, though less tightly than ARG mode does (Table 1). Across the five methods, the results separate into a genealogy-based family (ARG, local-tree and `PolarBEAR`) and an outgroup-based one (fixed-tree and `EST-SFS`), with MAP agreement 0.97–1.00 within the first and 0.999 within the second but only ~0.90 across them, a gap that reflects the distinct evidence each family draws on. Only the outgroup-based family has access to the outgroups here, so most disagreements are sites where both outgroups carry the ingroup-minor allele, which those methods take as ancestral while the genealogy assigns the ingroup MRCA the major allele. Where they disagree instead, the two outgroups carry different alleles, and the outgroup-based posterior is correspondingly flatter.

**Table 1:**
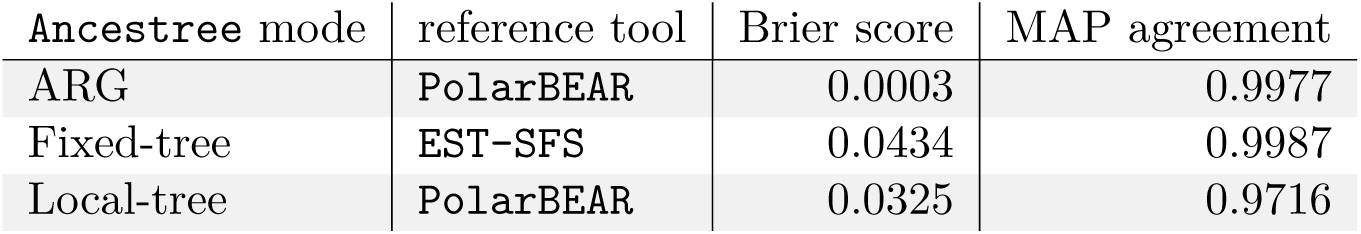
Each Ancestree mode reproduces its reference tool on the real MSL chr1 panel. Both scores compare each mode against the tool it subsumes, over the positions both emit. The full pairwise matrix across all five methods is in Appendix B.3 (Figure B1).

### 3.2 Comparison of the Inference Modes

The comparisons above used data satisfying the reference tools’ assumptions, where agreement is expected by construction. Here we instead evaluate all three of `Ancestree`’s inference modes across a panel of simulation scenarios, each of which deliberately breaks a different modelling assumption (recurrent mutation, incomplete lineage sorting, a misspecified outgroup topology, and selection).

#### Ingroup polymorphism

We classify sites by their ingroup allele pattern. A site is referred to as *ingroup-polymorphic* when the ingroup carries two or more alleles, biallelic in the usual case and poly-allelic when three or more segregate, and otherwise *ingroup-monomorphic*. An ingroup-monomorphic site is either fixed for the ancestral allele, and then trivially resolved at the ingroup MRCA, whose state is simply the allele every sample carries, or fixed for the derived allele, a divergence site whose ancestral state is recoverable only from the outgroups.

#### Inference modes

We compare the package’s three inference modes: ARG mode, run on the true simulated local trees (the achievable ceiling for a genealogy-based method); local-tree mode, which infers the genealogy from genotypes alone (Section 2.4); and fixed-tree mode, the outgroup-ladder fit (Section 2.3). The two genealogy modes use the stationary root prior. Fixed-tree collapses the ingroup to a polytomy, and instead uses the Kingman ingroup weight to exploit information from the ingroup allele frequencies (**Kingman weight**). All modes use JC69 (**Substitution models**) and the true phase. Section 3.4 examines local-tree mode’s performance under injected phasing switch error.

#### Simulation scenarios

We use five scenarios at realistic, human–ape-scale depths, the deepest outgroup splitting from the ingroup at *T_n_* 1.5 10^6^ generations with *N_e_* = 3 × 10^4^. Each scenario simulates *n* = 20 ingroup haplotypes and 10 outgroups, from which smaller subsets are taken below, selected evenly across the ten available splits by depth so that each subset spans the full divergence range rather than dropping the deepest outgroups first. Four are neutral `msprime` coalescent simulations, each run as ten independent replicates, and one, *purifying selection*, is a forward SLiM [Haller and Messer, 2019] simulation run as one hundred replicates. The *baseline* scenario is a neutral coalescent whose nested-outgroup topology is the one fixed-tree mode assumes (**The outgroup-ladder tree**), with the outgroups deep enough that incomplete lineage sorting is negligible, their splits log-spaced from 3 10^5^ to 1.5 10^6^ generations. It is the reference case that the other four scenarios each perturb.

The *hypermutation* scenario follows the baseline, with every site’s mutation rate raised tenfold, which roughly corresponds to the elevation reported for CpG sites in humans [Nachman and Crowell, 2000]. At this rate, a substantial fraction of sites sustain more than one mutation, so recurrent and back-mutations—and the homoplasy they create—are no longer negligible, breaking the infinite-sites assumption that the other scenarios largely satisfy. This also makes inference considerably harder, because the observed alleles no longer pin down a single mutation event: the same allele can arise on several branches or revert, so the ancestral state is no longer identified by the minimal-change reconstruction and the likelihood must integrate over competing mutation histories. The *strong ILS* scenario uses shallow splits (3 10^4^ to 2.1 10^5^ generations) under which the ingroup is non-monophyletic across 89% of the genome. This violates the fixed-tree assumption, since the local genealogy departs from the species tree at most loci, and it also makes the outgroups much less informative, because incomplete lineage sorting leaves polymorphisms shared between ingroup and outgroups.

The *outgroup clade* scenario lets the outgroups coalesce with one another first and join the ingroup together, at a single split 3 10^5^ generations back, breaching the outgroup-ladder assumption of fixed-tree mode. It also leaves less independent outgroup information, since the outgroups share much of their history with each other rather than diverging separately toward the ingroup. The *purifying selection* scenario simulates one hundred independent 100 kb replicates in which ingroup mutations are deleterious and semi-dominant, drawing their selection coefficient from a gamma distribution (mean −0.01, shape 0.2), while the outgroups evolve neutrally. Forward simulation is far costlier than the coalescent, so this scenario runs at a smaller effective population size, *N_e_* = 10^3^, with the rates scaled up tenfold and the split times down by the same factor to leave the scaled parameters unchanged. Each replicate is then recapitated to a complete coalescent history. Selection skews the site-frequency spectrum away from the neutral expectation—holding deleterious derived alleles at low frequency and rarely allowing them to reach high frequency—so fixed-tree mode’s Kingman weight, calibrated to a neutral SFS, is misspecified and the frequency-based signal it supplies becomes unreliable.

#### Recovery across the frequency spectrum

Figure 4 reports, for each of the three inference modes and each of the five simulation scenarios, the mean Brier score within each bin of the unfolded spectrum, one panel per outgroup count. Recovery worsens as the derived allele becomes more common, and worsens further still at the two ingroup-monomorphic tails of the spectrum, the fixed-ancestral bin (*i* = 0) and the fixed-derived bin (*i* = *n*), which differ substantially in difficulty. The fixed-derived bin, holding the divergence sites, is the hardest in every panel, since no ingroup sample carries the ancestral allele and the outgroups must supply all of the evidence for a configuration that is itself rare. At both tails, the focal node is the panel root rather than the ingroup MRCA, whose state a monomorphic ingroup already fixes. That choice divides the two classes: graded at *I* a fixed-ancestral site would be trivially correct and a divergence site maximally wrong, so only the deeper focal node can separate them, at the cost of leaving the fixed-ancestral class no longer trivial (Appendix C.1). The difficulty of the fixed-ancestral sites at the panel root comes from the outgroup branches, on which a mutation can make the ingroup allele look derived.

**Figure 4:**
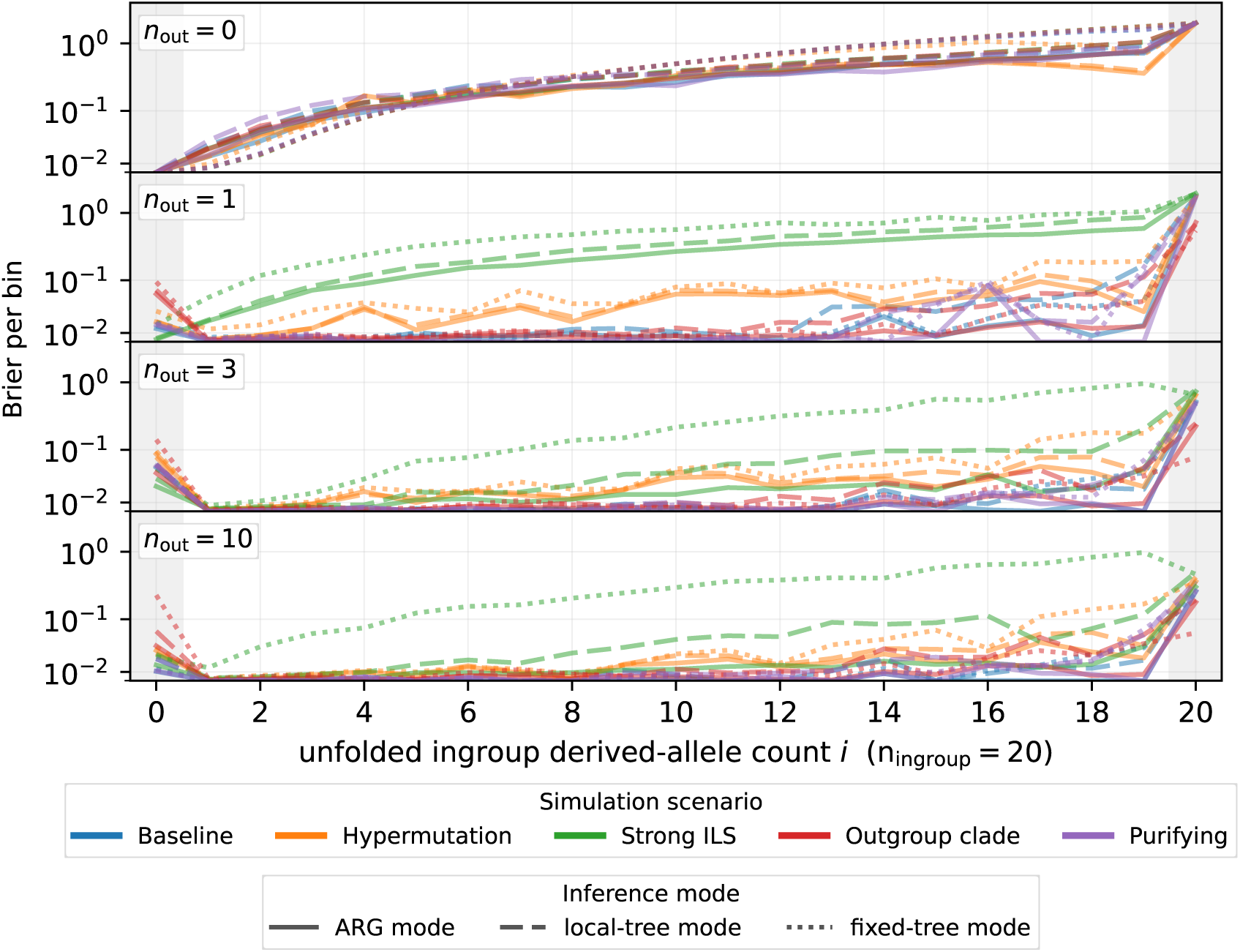
**Brier score across the unfolded SFS** for the three inference modes (linestyle) on all five scenarios (colour), one panel per outgroup count *n*_out_ ∊ {0, 1, 3, 10}. Lower is better. X-axis: unfolded ingroup derived-allele count *i ∊* {0*,...,* 20}, with the two ingroup-monomorphic tails shaded. Each bin is graded at the focal node its site class refers to: the ingroup MRCA where the ingroup is polymorphic, and the root of the whole panel at the two monomorphic tails.

#### Without outgroups

With no outgroup, every inference mode performs poorly across the whole spectrum and the five scenarios become nearly indistinguishable, the inference resting on the ingroup alone while what separates the scenarios lies mostly in their outgroups. An ingroup-monomorphic site then carries no evidence at all, the panel root being the ingroup MRCA whose state the single observed allele determines, so every mode places all its posterior mass on that allele and thereby resolves every fixed-ancestral site correctly and every divergence site incorrectly. Across the polymorphic interior, the lowest-frequency bins are the least poorly scored. A derived allele at low frequency subtends a small clade nested well below the root, from which the genealogy modes can identify the derived allele relatively well. Fixed-tree mode, reading the allele frequency alone, attains a marginally lower Brier score than either genealogy mode, at the cost of a substantially higher Brier score in the high-frequency bins.

The difficulty of recovering the ancestral allele grows with the derived-allele frequency, in what `PolarBEAR` terms a *non-informative genealogy* [Liang and Dutheil, 2026]: a common derived allele subtends most of the sample, so its mutation sits on a branch adjacent to the root and the two polarisations—placing that mutation on either side of the root—have similar likelihood (Appendix A.2). Such alleles are also rare *a priori*, the neutral expectation the Kingman ingroup weight encodes in fixed-tree mode (**Kingman weight**), so a confident assignment there requires correspondingly more outgroup evidence. Already well below mid-frequency, the ordering of the modes reverses, the topology remaining more informative than the frequency alone and giving ARG and local-tree mode an edge through the rest of the spectrum (Appendix C.2). Fixed-tree mode instead places little posterior mass on a common derived allele, a bias that is strongest under *purifying selection*, where most derived alleles segregate at low frequency.

#### Adding outgroups

A single outgroup already brings the polymorphic interior to a low Brier score that degrades only a little toward highest frequencies, the exceptions being *strong ILS* and *hypermutation*. Under strong ILS, the gain from a single outgroup is small for every mode, because that outgroup shares much of its polymorphism with the ingroup, and fixed-tree mode is the most affected. A third outgroup improves recovery substantially, including under strong ILS, and the improvement is by far the greatest at the divergence sites, which rest on outgroup evidence alone. The sites fixed for the ancestral allele do not benefit from the additional outgroups. With no outgroup, the panel root coincides with the ingroup MRCA, so they are trivially resolved, whereas adding outgroups moves the focal node deeper than the ingroup and makes them no longer trivially identifiable. Appendix C.1 varies the focal node between the ingroup MRCA and the panel root, and reports ancestral-allele recovery at every placement along that path.

Extending to ten outgroups brings diminishing returns across the polymorphic interior while the divergence sites keep improving, though they remain poorly resolved. The reason is that at an ingroup-monomorphic site the mutation may lie anywhere on the tree, and one close to the panel root could sit either on the branch above the most distant outgroup or on the branch subtending every other tip, two placements implying opposite ancestral states. The likelihood tends to favour the first, the distant outgroup branch offering more mutational opportunity, which leaves divergence sites confidently misassigned. Under *strong ILS*, the outgroups keep sharing polymorphism with the ingroup, and under *hypermutation*, each further outgroup branch brings more opportunity for recurrent and back-mutation alongside the evidence it carries. Both scenarios therefore remain poorly recovered however many outgroups are added, especially at high derived frequencies.

#### Across scenarios

The simulation scenarios differ in how far they depart from the *baseline*, but in each the two genealogy modes move together, local-tree mode closely tracking true-ARG mode throughout. *Hypermutation* degrades every mode across the entire spectrum, pervasive recurrent and back-mutation leaving many sites genuinely ambiguous. *Strong ILS* instead opens a wide gap between the genealogy modes and fixed-tree across the whole spectrum. The local gene tree there disagrees with the species tree at most loci, so fixed-tree mode is wrong wherever the genealogy departs from the topology it assumes and overweights the outgroup evidence accordingly, while the genealogy modes, which read the local tree itself, are far less affected. Local-tree mode nonetheless trails true-ARG mode visibly here, reconstructing the local trees being hardest where incomplete lineage sorting is pervasive. Under the *outgroup clade*, the outgroups share a common stem, on which a single mutation leaves every outgroup carrying an allele different from the ingroup’s, and the fitted ladder reads that as several independent observations rather than one event. Fixed-tree mode therefore overweights the outgroup evidence where the ingroup is monomorphic and is biased toward assigning those sites a derived state, which lowers its Brier score at the divergence sites and raises it at the sites fixed for the ancestral allele. *Purifying selection* differs little from the *baseline* once outgroups are present. It stands out only at *n*_out_ = 0, where the neutral Kingman calibration misreads the skewed spectrum. Figure 5 carries the same comparison averaged over different site classes.

**Figure 5:**
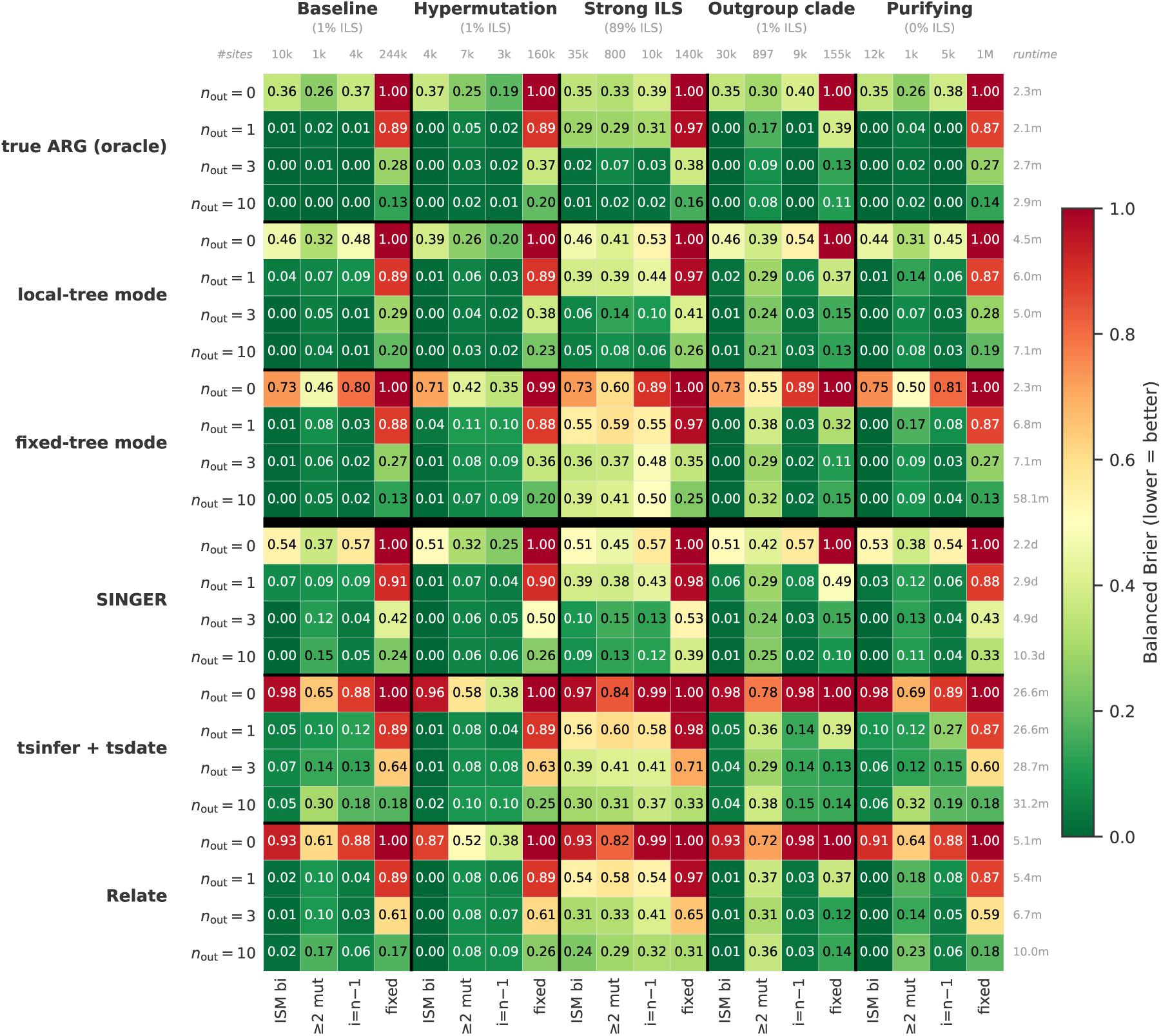
Inference modes and ARG-inference pipelines across scenarios, *n*_out_, and site classes. Balanced Brier (lower = greener = better) per cell, on a [0, 1] colour scale, for the true-ARG reference, local-tree mode, fixed-tree mode, `SINGER`, `tsinfer` and `Relate`, each varied over *n*_out_ ∊ {0, 1, 3, 10} over the five scenarios, each split into four site classes. The labels above each column give the number of sites in that class, those to the right of each row its runtime. `tsinfer` and `Relate` are oriented by a fixed-tree pass over the *n*_out_ outgroups rather than by the ground truth, while `SINGER` marginalises over that orientation at `-polar` 0.5. Every method is scored on the site set common to all, local-tree mode over an ensemble of *M* = 128 sampled genealogies. Versions: `tsinfer` 0.5.1 with `tsdate` 0.2.7, `Relate` 1.2.4, `SINGER` 0.1.8-beta.

### 3.3 Comparison with ARG-Inference Methods

Local-tree mode infers the genealogy from genotypes alone. To situate it among methods that also target the local genealogy, we compare it against three established ARG-inference pipelines that differ in what they require of the input—`tsinfer, Relate` and `SINGER`—over the scenarios and outgroup counts of Section 3.2 (Figure 5).

#### Site classes

Each cell of Figure 5 scores one method at one outgroup count on one scenario, over sites stratified into four classes rather than into frequency bins. Two of the classes divide the ingroup-polymorphic sites by mutation count: those carrying a single mutation, biallelic within the ingroup and the bulk of the panel (ISM bi), and those carrying more than one and therefore homoplastic (≥`2 mut`). The two remaining classes are single frequency bins, each folded with its mirror: the highest-frequency segregating bin paired with the singleton bin (i=n − 1), and the ingroup-monomorphic bins, fixed for the derived allele paired with fixed for the ancestral one (`fixed`). The mutation count separating the first two classes is taken over the full simulated tree with all outgroups present, so a site keeps its class as the number of outgroups is varied.

#### Balanced Brier

The rare high-frequency-derived and divergence sites are outnumbered many-fold by low-frequency polymorphisms and by sites fixed for the ancestral allele (Appendix C.6). A statistic restricted to the rare classes is therefore uninformative about the abundant ones, where even a small error rate affects many sites. To correct for this class imbalance, every cell reports a *balanced* Brier, 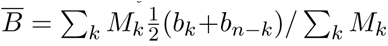 with *M_k_* = *m_k_* + *m_n_*_−_*_k_*, where *b_i_* is the mean Brier over the sites in derived-count bin *i*, *m_i_* is the number of such sites, and *k* = min(*i, n − i*) indexes the folded bins. Folded bins thus enter at equal weight, but the pairs in proportion to their site counts. To bound the computational cost and keep the rows comparable, every heatmap grades all of its methods on one common covered site set, each chunk contributing a genomic window that holds at most about 5,000 ingroup-segregating sites.

#### Orientation-dependent pipelines

`tsinfer` [Kelleher et al., 2019], dated with `tsdate` [Wohns et al., 2022], and `Relate` [Speidel et al., 2019] both require the ancestral allele as input to build their genealogies. We include the *n*_out_ outgroups among the samples each builds its ARG from, and orient the input not by the ground truth but by a fixed-tree pass over those same outgroups, then score the resulting ARG with ARG mode. Both tools behave similarly, and neither exceeds the accuracy of the fixed-tree pass that orients it, the ancestral-state assignment largely inheriting that orientation rather than correcting it. This is most damaging when few outgroups are available and the fixed-tree pass carries little information to orient the sites. At *n*_out_ = 0 it reduces to the site-frequency prior, which mis-orients the high-frequency-derived sites, and both tools then score worse than fixed-tree mode itself. Notably, their genealogies are thus limited by the ancestral states they are handed, a mis-oriented site biasing the tree built around it (Appendix C.9).

#### Orientation-free sampling

`SINGER` [Deng et al., 2025] is a full Bayesian ARG sampler that, unlike `tsinfer` and `Relate`, does not need the ancestral allele fixed. Its -polar flag sets the probability that the reference allele is ancestral, one value shared by every site, and at `-polar` 0.5 the orientation is left uninformative, so the sampler marginalises over it throughout. Its sample panel holds both the ingroup and the *n*_out_ outgroups, and we run thirteen independent Markov-chain Monte Carlo chains (300 iterations each, thinned every 20 with the first five draws discarded as warm-up, leaving 130 posterior ARG draws per site). The per-site kernel likelihood is marginalised over the sampled ARGs in likelihood space, as in the local-tree ensemble (see **Ensemble averaging**). Where the fixed outgroup topology is violated, it is more robust than fixed-tree mode, and its accuracy tracks local-tree mode’s closely across the scenarios, local-tree mode being the more accurate of the two in most cells.

#### Accuracy and cost

`SINGER`’s sampling from the full ARG posterior costs orders of magnitude more computation than the local-tree build, without a corresponding gain in accuracy. Fixed-tree mode is comparable to local-tree mode wherever its outgroup ladder holds, but it is the one mode whose runtime grows steeply with the number of outgroups, every additional outgroup adding a branch to the ladder it fits. Local-tree mode stays within minutes at every outgroup count and is the most accurate of the methods that infer the genealogy, so no method here recovers the ancestral allele from a plain VCF at lower computational cost. Additional comparison rows and the fitting details are given in Appendix B.5.

### 3.4 Local-Tree Mode Robustness

Local-tree mode infers dated local genealogies per window directly from a plain VCF via a pairwise-coalescent HMM (Section 2.4), then applies the ARG-mode kernel to it. We therefore examine the sensitivity of ancestral-allele recovery to the hyperparameters and to breaches of the modelling assumptions, and how faithfully the local trees themselves are recovered.

#### Setup

We first assess the sensitivity of local-tree mode’s ancestral allele recovery to its hyperparameters: the window size, the assumed recombination rate driving the HMM’s TMRCA resets, and the assumed mutation rate *µ*, which rescales every inferred TMRCA. Because the HMM pairs ingroup haplotypes, and unphased input exposes that pairing to switch errors, we additionally vary the switch-error rate. The underlying dataset is a single neutral ingroup-only `msprime` ARG (*n* = 20 haplotypes, *L* = 10^8^ bp, *N_e_* = 3 × 10^4^, *µ* = 1.25 × 10^−8^, *r*_true_ = 10^−8^, JC69; 265,079 polymorphic sites over 174,446 true local trees). From a common reference configuration (window = 8 SNPs, *r* = *r*_true_, *µ* = *µ*_true_, perfect phasing), we vary each input in turn. For each configuration we reconstruct the local trees from genotypes, infer the ancestral allele at each site, and score it against the ground truth. Local-tree mode is run throughout with the ensemble of *M* = 128 sampled genealogies used elsewhere, at a window of 8 SNPs over blocks of 4 SNPs unless stated otherwise. Running the same kernel on the genuine simulated genealogy through `ARGBasedInference` sets a true-ARG ceiling that isolates the cost of inferring the trees from the cost of using them. The analysis is ingroup-only (*n*_out_ = 0), so the ancestral state rests entirely on the inferred ingroup genealogy, making this the configuration most exposed to tree-inference error and hence the appropriate worst case for the sensitivities below. Appendix C.8 repeats the analysis with one and three outgroups.

#### Window size

A window is the contiguous run of SNPs constrained to share one inferred local tree (see **The pairwise HMM**). We vary it over more than three orders of magnitude, with the recombination and mutation rates set to their true values (Figure 6). The Brier score improves as the window narrows—finer windows resolve the genealogy at a recombination-faithful scale—and approaches, without reaching, the true-ARG ceiling. The window width sets the scale at which a single genealogy is imposed: a wide window forces sites separated by recombination breakpoints onto a common tree, mismatching most of them, whereas a single-SNP window lets each site draw on a genealogy local to it. Narrowing it deprives no tree of data, the HMM borrowing information along the whole sequence and the window setting only how finely its posterior is condensed into discrete trees. The cost is computational, one tree being inferred per window. Ancestral-allele recovery over the window and the HMM’s emission block jointly is shown in Figure C14.

**Figure 6:**
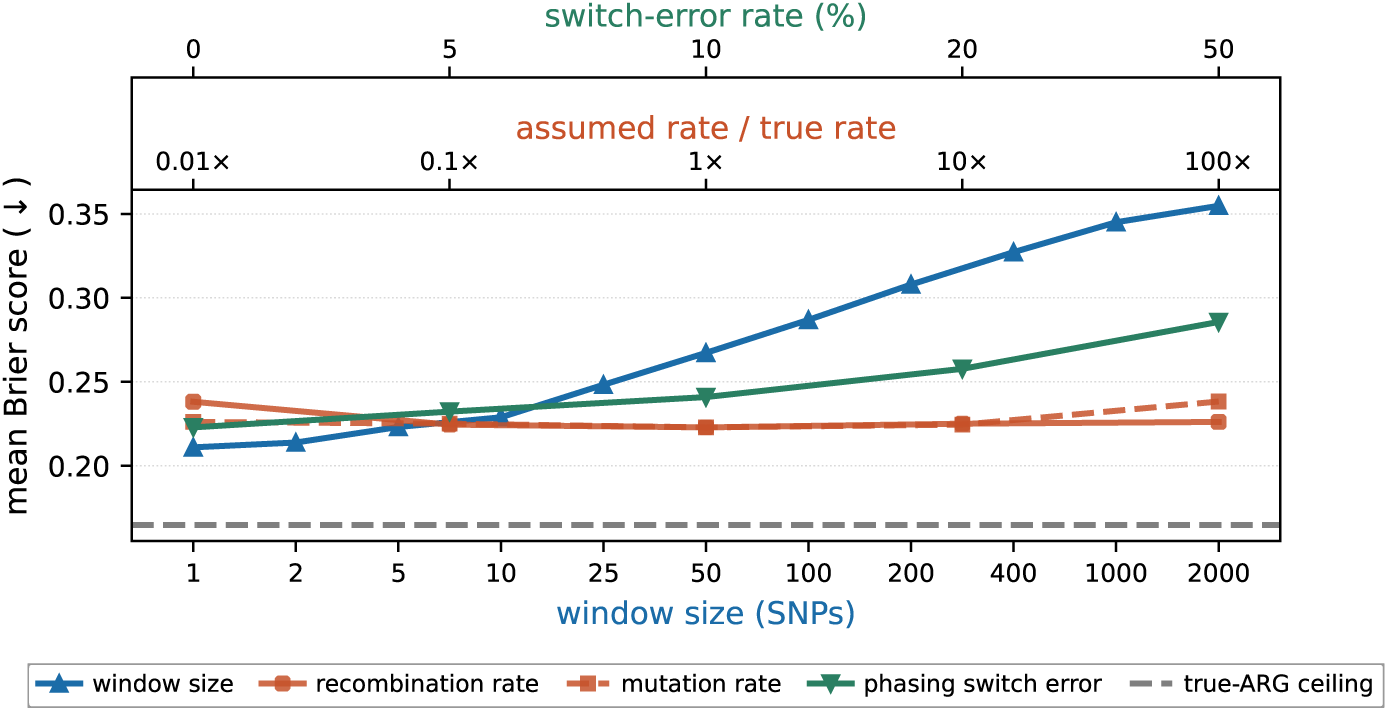
Ancestral-allele recovery against window size, rates and phasing. Mean Brier score (lower is better) as each input is varied in turn. The window is varied over three orders of magnitude at the true rates, recovery improving as it narrows. At the reference window of 8 SNPs, the assumed recombination and mutation rates are varied as multiples of the truth, leaving recovery essentially unchanged across four orders of magnitude, while injected phasing switch error degrades it only gently, remaining modest even at *s* = 0.5, where the phase is random. Every curve is produced with the ensemble of *M* = 128 sampled genealogies. The dashed line marks the true-ARG ceiling, the score ARG mode attains on the simulated genealogy itself, so the gap to it quantifies the cost of tree inference.

#### Rate misspecification and phasing

Holding the window at its reference value, we first misspecify the recombination and mutation rates from 0.01 to 100 times the truth (Figure 6). The assumed recombination rate exerts only a negligible effect, and misspecifying the mutation rate is essentially harmless, a global rescaling leaving the scale-invariant UPGMA topology intact. We also inject a phasing switch error, pairing consecutive ingroup haplotypes into pseudo-diploids and flipping the orientation of a pair at each heterozygous site with probability *s*, so that a rate of *s* = 0.5 leaves every heterozygous site independently oriented and amounts to random phasing. Accuracy degrades only gently under it, because the ancestral state is governed by the coarse genealogical structure—the ingroup topology and the allele-frequency configuration it induces—rather than by exact phase. The window size is therefore the most important determinant of how well that structure is recovered.

#### Genealogy recovery

The figures above score the final ancestral-allele assignment, and the gap to the true-ARG ceiling already attributes part of the error to tree inference rather than to the assignment itself, but it does not reveal how faithfully the genealogy is reconstructed. To examine that directly, we compare the inferred local trees against the simulated truth position by position, varying the window size (Figure 7). Three families of metrics serve the comparison, each averaged along the genome with every local tree weighted by the number of base pairs it spans. Rank agreement is the Spearman rank correlation between inferred and true pairwise TMRCAs (*TMRCA rank ρ*), which captures whether coalescences are ordered correctly regardless of absolute times. Absolute-time calibration is the mean log-ratio log(*t*^^^*/t*) (*TMRCA log-bias*) together with the slope of a *t*^^^ *t* least-squares regression (*TMRCA slope*), an unbiased, correctly-scaled estimator reading 0 and 1 respectively. Tree distance is the normalised Robinson–Foulds distance [Robinson and Foulds, 1981] (*Robinson–Foulds, norm.*), which reads topology alone and is 0 for identical splits, and the Kendall–Colijn distance [Kendall and Colijn, 2016] (*Kendall–Colijn*) at *λ* = 1, where the parameter *λ* ∈ [0, 1] tunes the metric from pure topology (*λ* = 0) to fully branch-length- (time-) aware (*λ* = 1). We also include the downstream mean polarisation Brier at each window size (*mean Brier*), relating genealogy recovery to ancestral-allele recovery.

**Figure 7:**
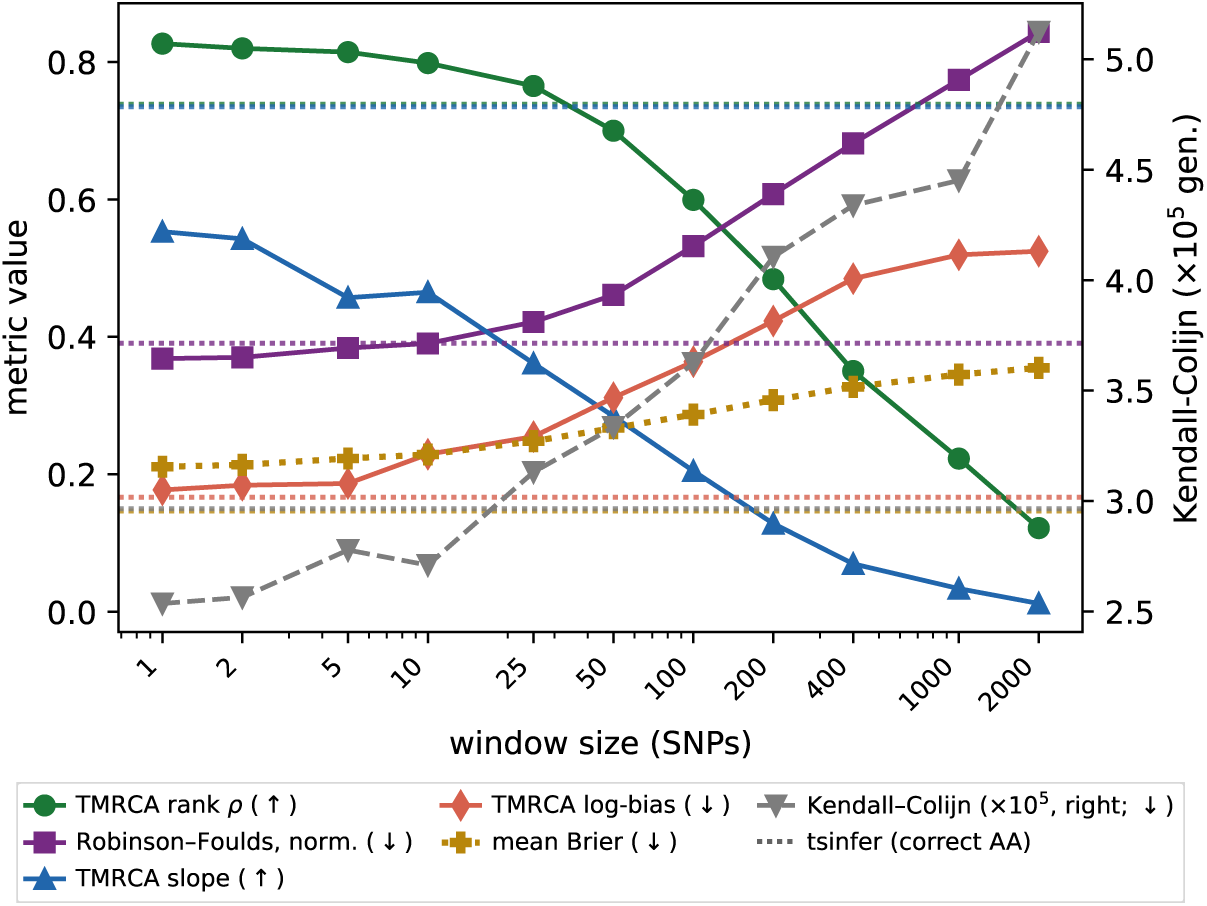
**Genealogy recovery vs. window size**: inferred local trees vs. the simulated truth on the 20-haplotype, 100 Mb sim (perfect phasing, correctly specified rates), with each local tree weighted by the number of base pairs it spans. All metrics are computed from the posterior-mean tree, except the mean Brier score, which comes from the *M* = 128 ensemble the mode scores with. Each metric’s better direction is marked in the legend. The dotted horizontals are a `tsinfer` ARG reference (correct ancestral allele; node times dated with `tsdate` at the true mutation rate).

The genealogy-recovery metrics (Figure 7) track ancestral-allele recovery: narrowing the window improves the inferred genealogy itself, the TMRCA rank correlation (TM-RCA rank *ρ*) rising from 0.12 at 2000 SNPs to 0.83 at the narrowest windows and the normalised Robinson–Foulds distance (Robinson–Foulds, norm.) falling from 0.84 to 0.37. The window’s effect on ancestral-allele recovery therefore acts mostly through genealogy reconstruction. The absolute TMRCAs, by contrast, are range-compressed at every window—the regression slope (TMRCA slope) reaches only ~0.55 at the narrowest windows and falls towards zero as they widen, while the log-bias (TMRCA log-bias) stays positive throughout—with the shortest coalescences pulled up and the deepest pulled down. This is the shrinkage of a posterior mean towards its prior, and comparable range compression is reported for other Bayesian coalescent-time estimators [Schweiger and Durbin, 2023]. A `tsinfer` ARG built from the same genotypes (dated with `tsdate` [Wohns et al., 2022] at the true mutation rate) provides a window-independent reference (dotted horizontals). At narrow windows local-tree mode matches or exceeds it on topology (Robinson–Foulds 0.37 vs. 0.39, TMRCA rank 0.83 vs. 0.74), with `tsinfer` better-calibrated only on absolute times (regression slope 0.73 vs. ~0.55). Here `tsinfer` is supplied the ground-truth ancestral allele, but yields only a small downstream-Brier edge (0.15 vs. 0.21); local-tree mode reaches comparable accuracy from genotypes alone, with no prior knowledge of the ancestral allele.

### 3.5 The Posterior-Weighted SFS

Bayesian ancestral-allele inference returns a calibrated posterior at every site rather than a single best estimate. A common downstream analysis aggregates these posteriors into an unfolded site-frequency spectrum, and we demonstrate here how two common ways of collapsing the posterior each bias that spectrum, since both discard information in a frequency-dependent way. We illustrate them on the baseline coalescent scenario (Section 3.2; 20 ingroup haplotypes under JC69), here provided with no outgroup (*n*_out_ = 0) so that the ancestral state is informed by the ingroup genealogy alone. ARG-mode posteriors are scored against the simulator’s true ingroup derived-allele count at the biallelic segregating sites (Figure 8).

**Figure 8:**
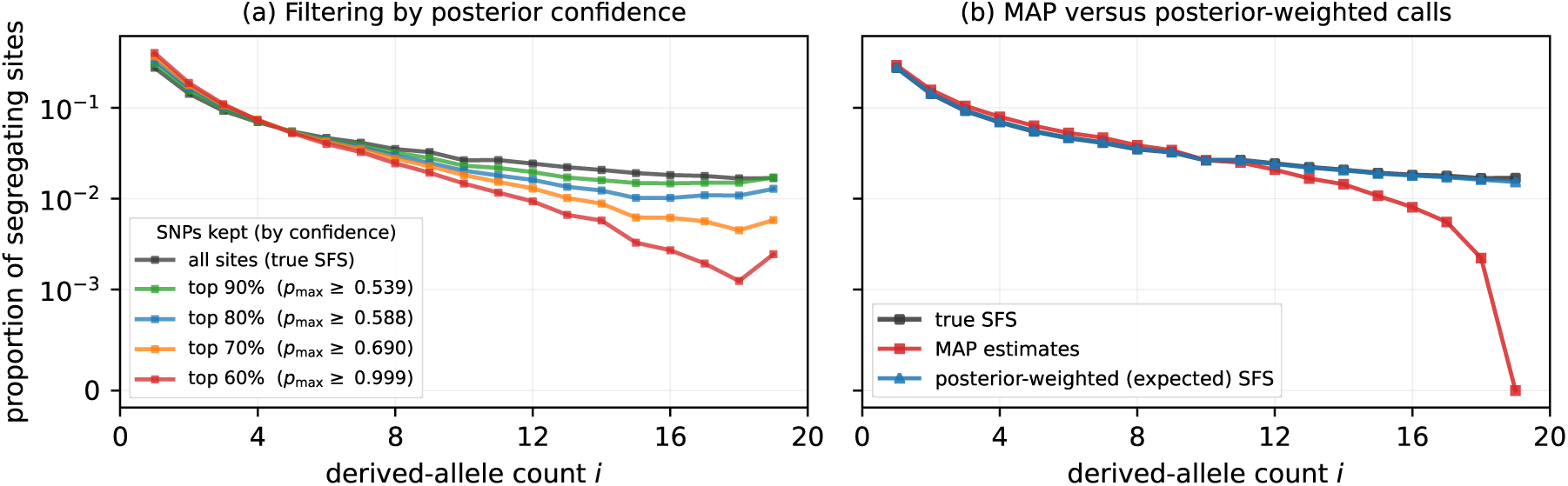
Confidence filtering and MAP estimates distort the unfolded site-frequency spectrum. ARG-mode posteriors on the synthetic baseline scenario with no outgroup (*n*_out_ = 0); each curve is the proportion of segregating sites at every true ingroup derived-allele count *i*, on a logarithmic axis. **(a)** Sites are binned by their true derived-allele count, so the black curve (all sites retained) is the true spectrum; retaining only the most-confident fraction of sites (largest posterior mass *p*_max_) depletes the high-frequency-derived tail and correspondingly inflates the low-frequency bins, because confidence falls with derived-allele frequency. **(b)** Committing each site to its MAP ancestral allele (red) similarly depletes the high-frequency tail relative to the truth (black), whereas the posterior-weighted (expected) spectrum (blue) almost completely recovers the true shape.

First, discarding low-confidence sites when computing the SFS is not frequency-neutral: posterior confidence falls as the derived allele becomes more common, so retaining only the most-confident sites preferentially removes high-frequency-derived variants and erodes the high-frequency tail. Second, even without any filtering, committing each site to its maximum-a-posteriori allele depletes the same region of the spectrum: where the derived allele is at high frequency the ancestral state is genuinely uncertain, and the MAP estimate resolves that uncertainty in favour of the major allele, moving sites out of the high-frequency-derived bins. Both distortions arise from the same high-frequency uncertainty, so the no-outgroup case shown here is the most severe: supplying outgroups raises the certainty of the high-frequency assignments and correspondingly attenuates the bias (Appendix C.7).

Retaining every site and weighting it by its ancestral-state posterior instead removes both distortions: each biallelic site adds its posterior probability for each candidate ancestral allele to the derived-count bin that allele implies—splitting an uncertain site between bins *i* and *n − i*—so the per-site posteriors sum into a posterior-weighted expected unfolded spectrum. This recovers the true shape across the whole frequency range, including the high-frequency tail that MAP estimates deplete (Figure 8b), and it does so with far fewer outgroups than reliable site-wise assignments would require, since the aggregate spectrum tolerates per-site uncertainty that a MAP estimate cannot.

## 4 Discussion

`Ancestree` unifies two approaches to ancestral-allele inference—outgroup-based fixed-tree reconstruction and outgroup-free ARG-based reconstruction—within a single probabilistic framework, and adds a third, local-tree mode that bridges them by inferring the per-site genealogy from genotypes alone.

### Local-tree mode as default

Our benchmarks show that local-tree mode improves on fixed-tree mode without outgroups, is comparable to it once outgroups are available, and approaches the true-ARG ceiling (Figure 5). It also remains accurate under phasing error and under misspecified mutation and recombination rates (Section 3.4), is the more robust of the two wherever the fixed-tree topology is violated, and its runtime scales more favourably with the number of outgroups. Its pairwise-coalescent HMM scales quadratically in the number of samples (Figure C16) but stays practical up to a few hundred individuals, and subsampling ingroup individuals should sacrifice little power (Appendix C.2). It is therefore an accurate and practical default, and fills a gap left open by current tools: genealogy-aware ancestral-allele inference from plain genotype data. Its annotations can in turn supply the fixed ancestral states required by ARG builders such as `tsinfer` and `Relate`.

### Outgroups and posteriors

The comparison across simulation scenarios and outgroup configurations (Section 3.2) yields several practical conclusions. The ingroup topology carries only a limited signal about the ancestral state, so recovery rests mainly on the outgroup evidence. This is especially true at sites where the ingroup is fixed for the derived allele, which carry no within-ingroup signal at all. Outgroup depth and independence matter as much as number: separately diverging outgroups are more informative than a clade of close relatives, and they should fall outside the ingroup’s variation without being so deep that additional mutations erode the signal. Homoplasy degrades recovery throughout, although enough outgroups restore most of it. How much outgroup evidence is needed depends on what the annotation is for. An aggregate spectrum needs far fewer than site-wise inference, since weighting each site by its ancestral-state posterior integrates over the per-site uncertainty, so the expected spectrum recovers its true shape with one or a few outgroups, whereas site-wise posteriors keep improving as further outgroups are added. One response to a diffuse posterior is to retain only the sites whose posterior is concentrated, but for summary statistics such as the SFS this filtering introduces a bias: confidence falls with derived-allele frequency, so discarding low-confidence sites biases the SFS (Section 3.5). Since every per-site output is a full posterior over the four nucleotide states, we recommend integrating over that uncertainty downstream wherever possible, and restricting to a confident subset only where the sites themselves, rather than summaries of them, are the object of interest.

### Implementation

We designed the `Ancestree` implementation to be computationally efficient and to scale to genome-scale datasets across its three inference modes. Each bounds memory differently: fixed-tree mode streams the variant data (flat in sequence length), local-tree mode chunks the genome to cap its quadratic pairwise-HMM matrix, and ARG mode shares a single in-memory ARG across cores (Appendix C.13). The package reads and writes VCF, Zarr and `tskit` ARG data alike, facilitating integration into existing pipelines. It provides both a Python API and a command-line interface, documented online with worked examples (`ancestree.readthedocs.io`).

### Limitations and outlook

`Ancestree`’s ARG mode is bounded by the quality of the input ARG; in particular, ancestral-allele errors embedded during the ARG’s construction cannot be corrected by re-inferring ancestral alleles from that ARG. In practice it is therefore best paired with externally inferred SMC ARGs that do not take the ancestral allele as input, rather than with ARG builders such as `tsinfer` and `Relate` that condition on the ancestral allele (Appendix B.5). Uncertainty should be quantified not only in the ancestral state but also in the genealogy it is read from. Local-tree mode does so at little additional cost, marginalising each window’s site likelihood over *M* genealogies sampled per haplotype pair (Figure C12), which approximates the joint posterior over ARGs and substantially improves ancestral-state calibration relative to a single point-estimate genealogy (Figure C11). A natural avenue for future work is thus propagating ancestral-allele uncertainty into ARG inference itself, as `SINGER` [Deng et al., 2025] does for the genealogy. More broadly, by relying on one likelihood kernel across three interchangeable modes, `Ancestree` lets the available genealogy—assumed, supplied, or inferred—rather than the choice of tool dictate how each dataset is polarised, and returns per-site posteriors that downstream analyses can propagate.

## Abbreviations

ARG: Ancestral Recombination Graph
ASR: Ancestral-Sequence Reconstruction
F81: Felsenstein 1981 (substitution model)
GTR: General Time-Reversible (substitution model)
HKY: Hasegawa–Kishino–Yano (substitution model)
HMM: Hidden Markov Model
ILS: Incomplete Lineage Sorting
ISM: Infinite-Sites Model
JC69: Jukes–Cantor 1969 (substitution model)
JIT: Just-In-Time (compilation)
K2: Kimura Two-Parameter (substitution model)
L-BFGS-B: Limited-memory BFGS with Bound constraints
MAP: Maximum A Posteriori
MLE: Maximum Likelihood Estimate
MRCA: Most Recent Common Ancestor
MSL: Mende in Sierra Leone (1000 Genomes population)
PSMC: Pairwise Sequentially Markovian Coalescent
SFS: Site-Frequency Spectrum
SMC: Sequentially Markovian Coalescent
SNP: Single-Nucleotide Polymorphism
TMRCA: Time to the Most Recent Common Ancestor
UPGMA: Unweighted Pair Group Method with Arithmetic mean
VCF: Variant Call Format

## Code availability

The software is freely available at github.com/Sendrowski/Ancestree, and documentation as well as usage examples can be found at `ancestree.readthedocs.io`.

## Data availability

The real-data analyses use the MSL chr1 panel and pre-computed ancestral-state posteriors distributed with `PolarBEAR` [Liang and Dutheil, 2026] at doi.org/10.17617/3.9JG13K. All simulated data are generated by the workflow in the software repository, which also reproduces every figure and table in this manuscript.

## A Illustrations and Method Details

### A.1 Worked-Example Gallery

Eight worked examples (Figure A1) illustrate what the kernel returns under representative configurations, each panel running the likelihood kernel directly on a small tree and site. They span the configurations the library handles natively: (1) a baseline ARG-mode polymorphic site, (2) a poly-allelic site with three observed alleles, (3) a missing outgroup tip handled by all-ones partial marginalisation, (4) a monomorphic-ingroup site evaluated on the deep-rooted ladder tree (`OutgroupLadderTree`), (5) a homoplastic site where the same allele appears on two independent branches and the full exp(**Q***t*) kernel returns an equiprobable posterior, (6) a root polytomy, (7) a transition-biased site under K2 with *κ* = 10, and (8) the standard fixed-tree workflow on `OutgroupLadderTree` with a polymorphic ingroup.

**Figure A1:**
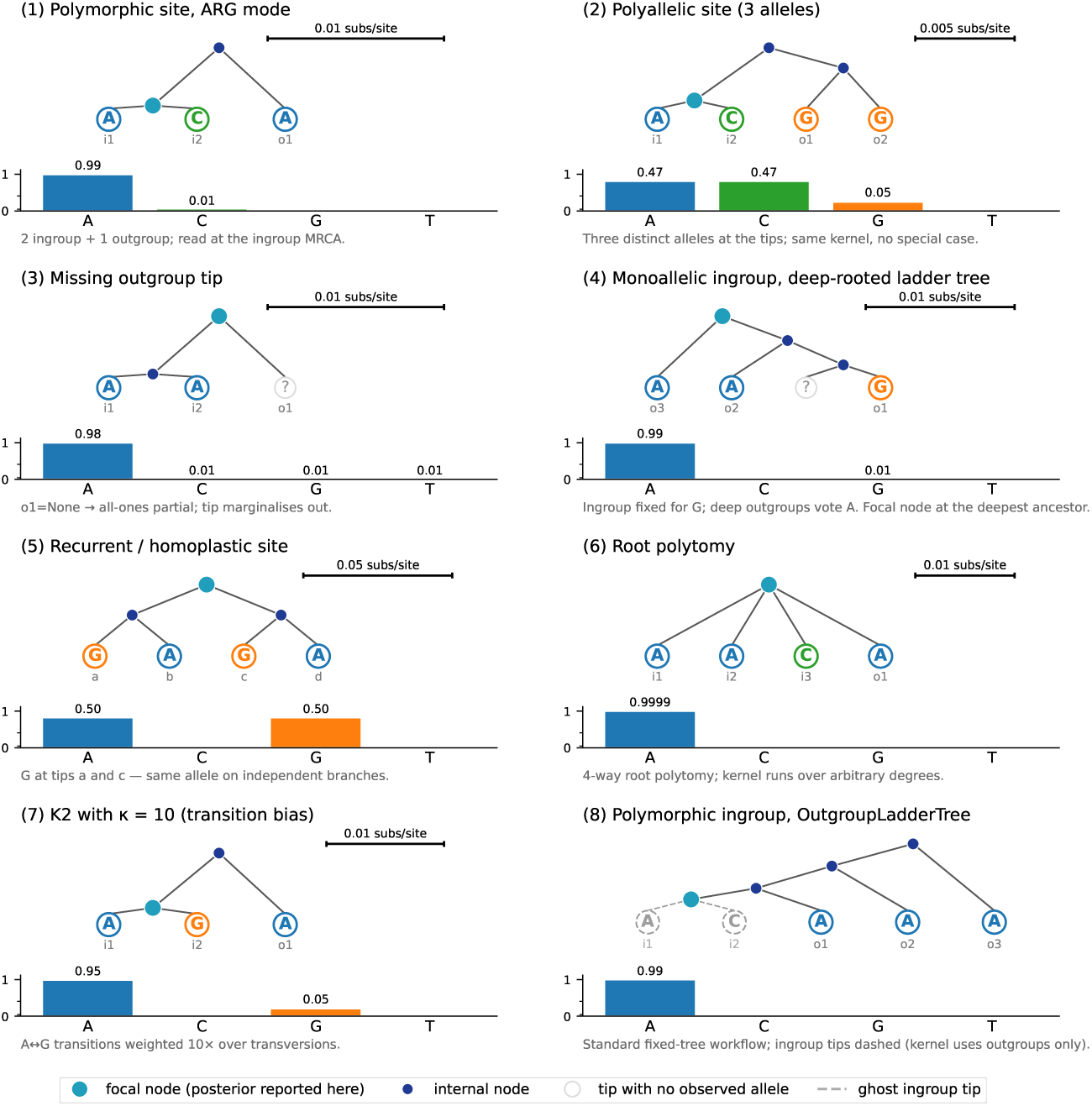
Worked-example gallery. The eight (tree, site) configurations enumerated in the text, with the posterior over {*A, C, G, T*} at the inference root shown as a four-bar strip beneath each tree. Tree branches drawn as straight diagonals proportional to fitted branch length; per-panel scale bar (top-right) gives the substitutions-per-site corresponding to the visual span. For `OutgroupLadderTree` (case 8), the outgroups form a nested ladder on the right, joining the ingroup lineage successively, whereas the ingroup—collapsed into the root by construction—is shown as a dashed cherry on the left; only the outgroup tips enter the kernel.

### A.2 Root-State Posteriors without Outgroups

Without outgroups the root-state posterior is determined by the observed allele pattern alone, which we illustrate on a single six-tip ingroup genealogy with varying tip alleles (Figure A2). A low-frequency derived allele is oriented almost certainly—a singleton, or its near-fixed mirror—because only one orientation is parsimonious on the tree. When the two deepest clades carry different alleles the root is genuinely ambiguous, the posterior splitting evenly between them, since either allele requires a single change on the backbone branch. A homoplastic pattern, the same allele on two separate clades, is again resolved confidently, the kernel preferring the orientation that invokes fewer independent mutations.

**Figure A2:**
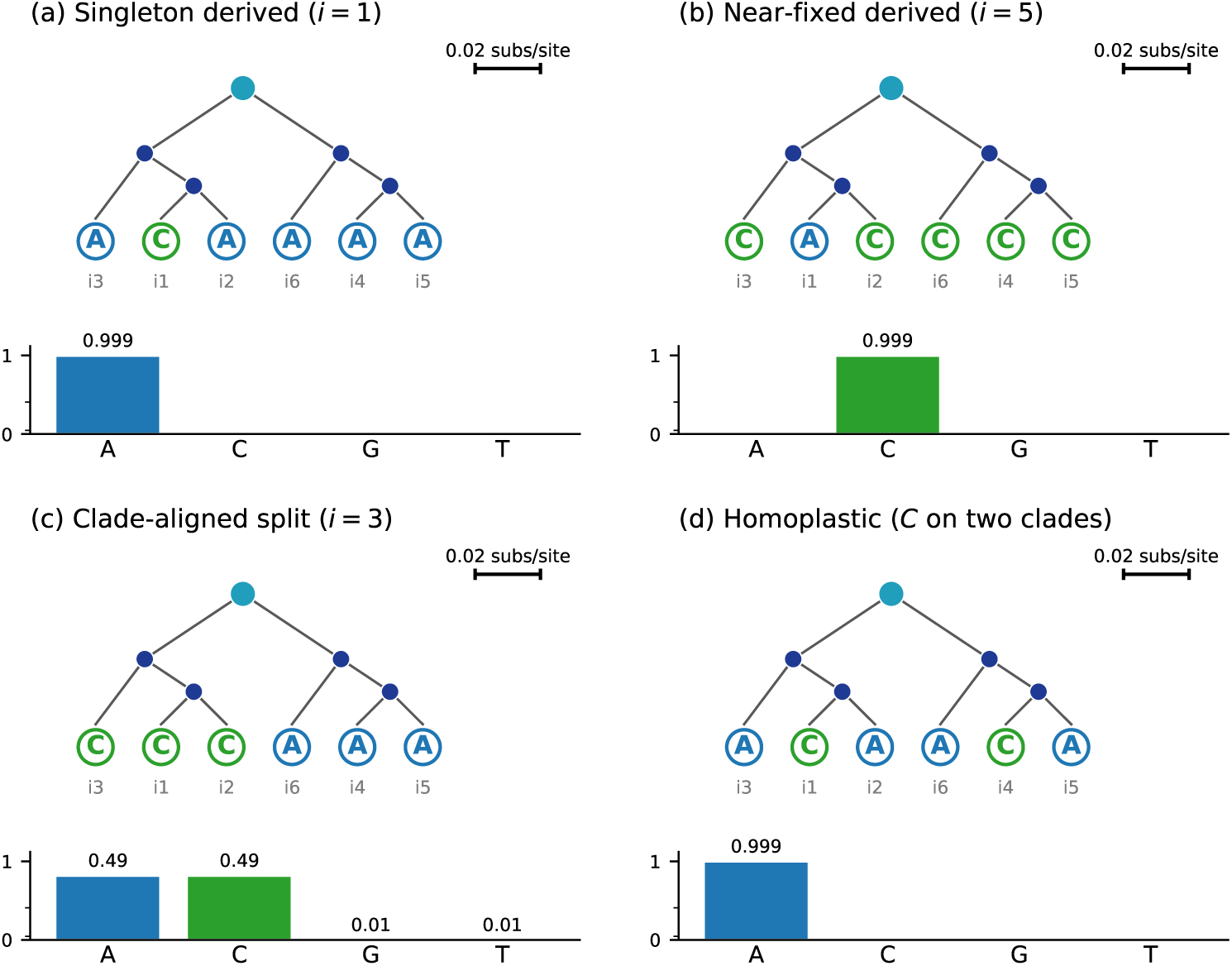
Root-state posteriors for four example site patterns on one fixed ingroup-only genealogy. The same six-tip ingroup tree is shown for different tip alleles; beneath each is the posterior over {*A, C, G, T*} at the inference root. **(a)** A singleton derived allele and **(b)** its near-fixed mirror are oriented almost certainly. **(c)** When the two deepest clades carry different alleles the root is equiprobable between them. **(d)** A homoplastic pattern is resolved towards the orientation requiring fewer independent mutations.

### A.3 Edge Cases

#### Sites outside the common envelope

The main benchmarks restrict to the biallelic, parsimony-resolvable, ingroup-polymorphic sites that all three tools accept (Section 3), since outside that intersection the reference tools filter or mishandle the input. Table A1 reports kernel-level posteriors on four classes that fall outside it, on a balanced quartet ((`i0, i1`), (`i2, i3`)) under JC69, a 4-way star for the polytomy class, each shown first without outgroups and then with three outgroups attached as a nested ladder. `Ancestree` and `PolarBEAR` both return a full (`A, C, G, T`) vector and agree throughout except on the *tri-allelic* site, where `Ancestree`’s sum-product marginal breaks the tie toward the ingroup-majority A (0.60) without an outgroup and `PolarBEAR`’s max-product keeps the three-way 0.33 tie. `EST-SFS` instead reports only the scalar *P* ^maj^, the posterior that the ingroup-major allele is ancestral, and only with outgroups.

**Table A1:**
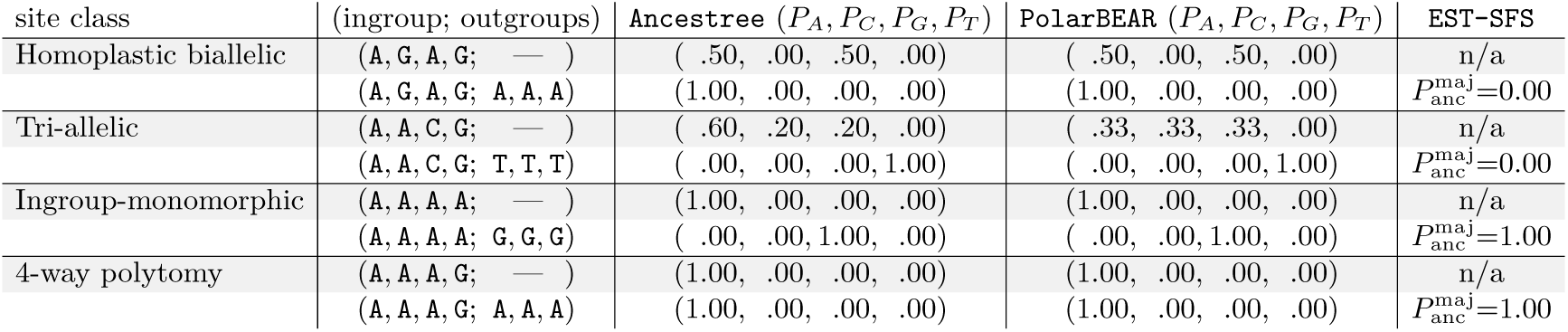
Kernel-level posteriors on four site classes, each without and then with three outgroups. `Ancestree` (sum-product) and `PolarBEAR` (max-product) return a full (*A, C, G, T*) posterior; `EST-SFS` reports only the scalar *P* ^maj^, the posterior that the ingroup-major allele is ancestral, and only with outgroups (n/a otherwise).

| site class | (ingroup; outgroups) | **Ancestry** ($P_A, P_C, P_G, P_T$) | **PolarBEAR** ($P_A, P_C, P_G, P_T$) | **EST-SFS** |
| --- | --- | --- | --- | --- |
| Homoplastic biallelic | (A, G, A, G; — ) | (.50, .00, .50, .00) | (.50, .00, .50, .00) | n/a |
| | (A, G, A, G; A, A, A) | (1.00, .00, .00, .00) | (1.00, .00, .00, .00) | $P_{\text{anc}}^{\text{maj}}=0.00$ |
| Tri-allelic | (A, A, C, G; — ) | (.60, .20, .20, .00) | (.33, .33, .33, .00) | n/a |
| | (A, A, C, G; T, T, T) | (.00, .00, .00, 1.00) | (.00, .00, .00, 1.00) | $P_{\text{anc}}^{\text{maj}}=0.00$ |
| Ingroup-monomorphic | (A, A, A, A; — ) | (1.00, .00, .00, .00) | (1.00, .00, .00, .00) | n/a |
| | (A, A, A, A; G, G, G) | (.00, .00, 1.00, .00) | (.00, .00, 1.00, .00) | $P_{\text{anc}}^{\text{maj}}=1.00$ |
| 4-way polytomy | (A, A, A, G; — ) | (1.00, .00, .00, .00) | (1.00, .00, .00, .00) | n/a |
| | (A, A, A, G; A, A, A) | (1.00, .00, .00, .00) | (1.00, .00, .00, .00) | $P_{\text{anc}}^{\text{maj}}=1.00$ |

#### Marginal versus joint root states

The kernel can read the root state in two ways, which answer distinct questions. Summing over each child’s states marginalises the internal nodes, so the normalised root vector is the marginal posterior *P* (*r* = *s* **x**), the probability of state *s* after averaging over all internal-node histories. Replacing that sum with a max*_s_′* gives the max-product algorithm, which returns the single most-probable joint assignment over the internal nodes and reads the root state from it, the route `PolarBEAR` [Liang and Dutheil, 2026] takes. The two can differ, since a node’s marginal MAP need not match its state in the joint-MAP assignment, but they coincide wherever the genealogy and the observed alleles pin the ancestral history down, which is almost everywhere: the per-site MAP estimates agree at almost every MSL chr1 position both emit (Appendix B.4). `Ancestree` uses sum-product, because the marginal posterior scores each candidate ancestral allele on its own rather than conditioning on one most-probable history, and it reports the full per-state vector rather than its largest entry alone. Where two states are exactly equally likely, an argmax must fall back on an arbitrary tie-break, whereas the posterior simply reports them as equiprobable.

## B Validation and Baselines

### B.1 Agreement with EST-SFS (Fixed-Tree Mode)

`EST-SFS` [Keightley and Jackson, 2018] is the standard fixed-tree ancestral-state inference tool: a maximum-likelihood fit of an outgroup-ladder tree under JC69 with a truncated-Poisson per-branch transition probability, combined with an implicit Kingman prior on the ingroup site-frequency spectrum. We assess whether `FixedTreeInference` reproduces `EST-SFS`’s per-site ancestral-state posteriors and fitted branch lengths on data satisfying its assumptions (biallelic, low-recurrence).

#### Setup

We simulated one `msprime` ARG with one ingroup population (*n* = 40 haploid samples) and three nested-split outgroup populations at split times {5 × 10^4^, 10^5^, 1.5 × 10^5^} generations (closest first, one haploid sample each), sequence length *L* = 2 × 10^6^, JC69 mutations—1,656 ingroup-polymorphic sites among the 11,151 segregating in the full alignment. Both `Ancestree` and `EST-SFS` run with JC69 inference and use 10 independent random initialisations to optimise the branch lengths. We compare results for *n*_out_ ∊ {1, 2, 3} (closest-first subset of the three available outgroups). The same simulated VCF is passed to both tools.

#### Results

Table B1 reports the per-branch ladder rates *K*^^^*i* fitted by each tool, and Table B2 the per-site agreement metrics and runtime. On data satisfying the `EST-SFS` assumptions the two tools are equivalent: the ladder rates match to within 1–3% at every cell, and the per-site estimates reach 99.94% MAP recovery with zero MAP disagreements.

**Table B1:** Per-branch ladder rates. One row per branch index *i* ∈ {0, 1*,...,* 2(*n*_out_ − 1)} of the outgroup ladder (Section 2.3). *K*^^^*i* is each tool’s ML fit on the same VCF (subs/site).

| $n_{\text{out}}$ | $i$ | $\hat{K}_i$ (subs/site) | | rel. diff |
| --- | --- | --- | --- | --- |
|  |  | EST-SFS | Ancestrree |  |
| 1 | 0 | $1.05 \times 10^{-3}$ | $1.06 \times 10^{-3}$ | +1.0% |
| 2 | 0 | $4.28 \times 10^{-4}$ | $4.38 \times 10^{-4}$ | +2.4% |
| | 1 | $6.19 \times 10^{-4}$ | $6.19 \times 10^{-4}$ | +0.0% |
| | 2 | $1.62 \times 10^{-3}$ | $1.62 \times 10^{-3}$ | +0.0% |
| 3 | 0 | $4.28 \times 10^{-4}$ | $4.38 \times 10^{-4}$ | +2.4% |
| | 1 | $6.19 \times 10^{-4}$ | $6.19 \times 10^{-4}$ | +0.0% |
| | 2 | $5.13 \times 10^{-4}$ | $5.13 \times 10^{-4}$ | +0.0% |
| | 3 | $1.10 \times 10^{-3}$ | $1.10 \times 10^{-3}$ | −0.0% |
| | 4 | $2.10 \times 10^{-3}$ | $2.10 \times 10^{-3}$ | −0.0% |

**Table B2:**
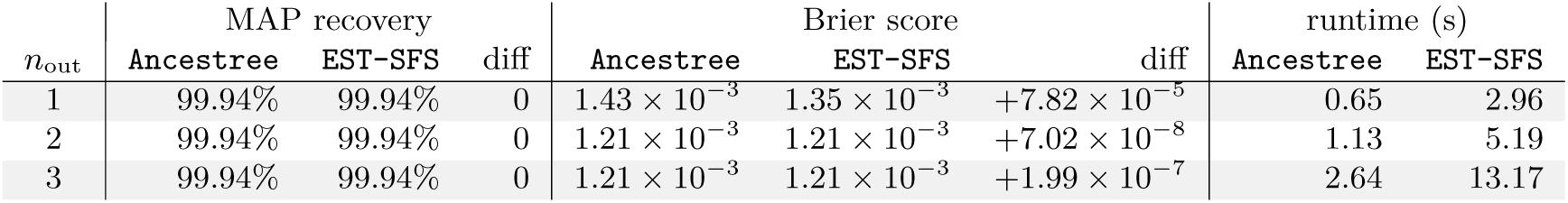
Per-site agreement and runtime. One row per *n*_out_, with metrics computed on the ingroup-biallelic sites. Each diff column is `Ancestree` minus `EST-SFS`, in percentage points for MAP recovery and in Brier score at the posterior level.

### B.2 Agreement with PolarBEAR (ARG Mode)

`PolarBEAR` [Liang and Dutheil, 2026] is an ARG-based ancestral-state tool: it reads per-site local trees from an inferred ARG and computes max-product ML ancestral-allele probabilities under JC69, without outgroups. On the ingroup-biallelic sites compared here, `ARGBasedInference` coincides with its first-order JC kernel. We simulated one `msprime` ARG with 20 diploid individuals, that is 40 sample haplotypes, from a single panmictic population (*L* = 2 × 10^6^) and no outgroups, and gave both tools the same. trees file. On that shared ARG the two reproduce each other’s per-site MAP estimates exactly, recover the simulated ancestral allele at the same sites, and score the same mean Brier to six decimal places (Table B3).

**Table B3:** Agreement with PolarBEAR on a 20-sample, 2 Mb panmictic `msprime` simulation. Both tools read the same ARG and score the same 3,348 ingroup-biallelic sites, agreeing on every MAP estimate. MAP recovery and the Brier score are taken against the simulated truth. The diff row is `Ancestree` minus `PolarBEAR`, and runtime excludes data load and output write.

| tool | MAP recovery | Brier score | runtime |
| --- | --- | --- | --- |
| **Ancestrtree** ARG mode | 88.3% | 0.1361 | 0.79 s |
| **PolarBEAR** | 88.3% | 0.1361 | 1.02 s |
| diff | 0 | $+8.60 \times 10^{-8}$ | −0.23 s |

### B.3 Agreement on the MSL chr1 Panel

Real data offer no ground truth, so we measure inter-tool agreement instead, on the published 1000 Genomes MSL chr1 panel that the `PolarBEAR` study uses.

#### Setup

The panel comprises 87 human MSL diploid individuals (174 haplotypes) and two outgroup species—ponAbe2 (orangutan) and macFas5 (crab-eating macaque)—across the whole of chromosome 1. The pseudo-VCF distributed with `PolarBEAR` covers 157,591,304 aligned positions across chr1. Of those, 156,451,219 (99.28%) are ingroup-monomorphic (ingroup fixed at a single allele) and 1,140,085 (0.72%) are ingroup-polymorphic SNPs—1,137,196 biallelic (99.75% of polymorphic) and 2,889 poly-allelic (0.25%). We run `Ancestree` in all three modes, both for direct comparability against each reference tool and to contrast the outgroup-based and genealogy-based paths on the same panel. Fixed-tree mode attaches the two outgroups and fits the `EST-SFS`-style outgroup ladder under HKY with the Kingman SFS prior. ARG mode uses no outgroups and reads the same `gamma-SMC` [Schweiger and Durbin, 2023] ARG as `PolarBEAR` under JC69, matching that tool’s configuration. Local-tree mode also uses JC69 and no outgroups, and infers the local trees from the ingroup genotypes alone.

#### Agreement

With no ground truth on real data, we measure pairwise agreement between the tools at the positions both methods score (Figure B1), reporting both the Brier agreement and the MAP agreement (**Metrics**). Both kernels score every site, `PolarBEAR` reducing its output to a published subset only afterwards (Coverage, below), so we compare the two over its full per-site output. Across the full output ARG mode and `PolarBEAR` agree at Brier 0.0003 (MAP 0.9977)—the residual confined to the non-informative class, where the genealogy alone cannot orient the site and the two tie-break it differently. Fixed-tree mode tracks `EST-SFS` at Brier 0.0434 (MAP 99.87%). Notably, local-tree mode, which infers the genealogy from a plain VCF, agrees with `PolarBEAR` at Brier 0.0325 (MAP 0.9716) and with ARG mode on the `gamma-SMC` ARG at Brier 0.0320 (MAP 0.9724). Its shortfall against the full-ARG methods is therefore genealogy-inference noise rather than a systematic gap. More broadly, the agreement separates into two blocks: the genealogy-based methods (`Ancestree` ARG, `Ancestree` local-tree, `PolarBEAR`) agree at 0.97–1.00 and the outgroup-based methods (`Ancestree` fixed-tree, `EST-SFS`) at 0.999, while across the two families it falls to ~0.90. The two families read different evidence, so the gap is expected rather than a defect: `Ancestree`’s own modes disagree at the same level (90.5%, fixed-tree against ARG).

**Figure B1:**
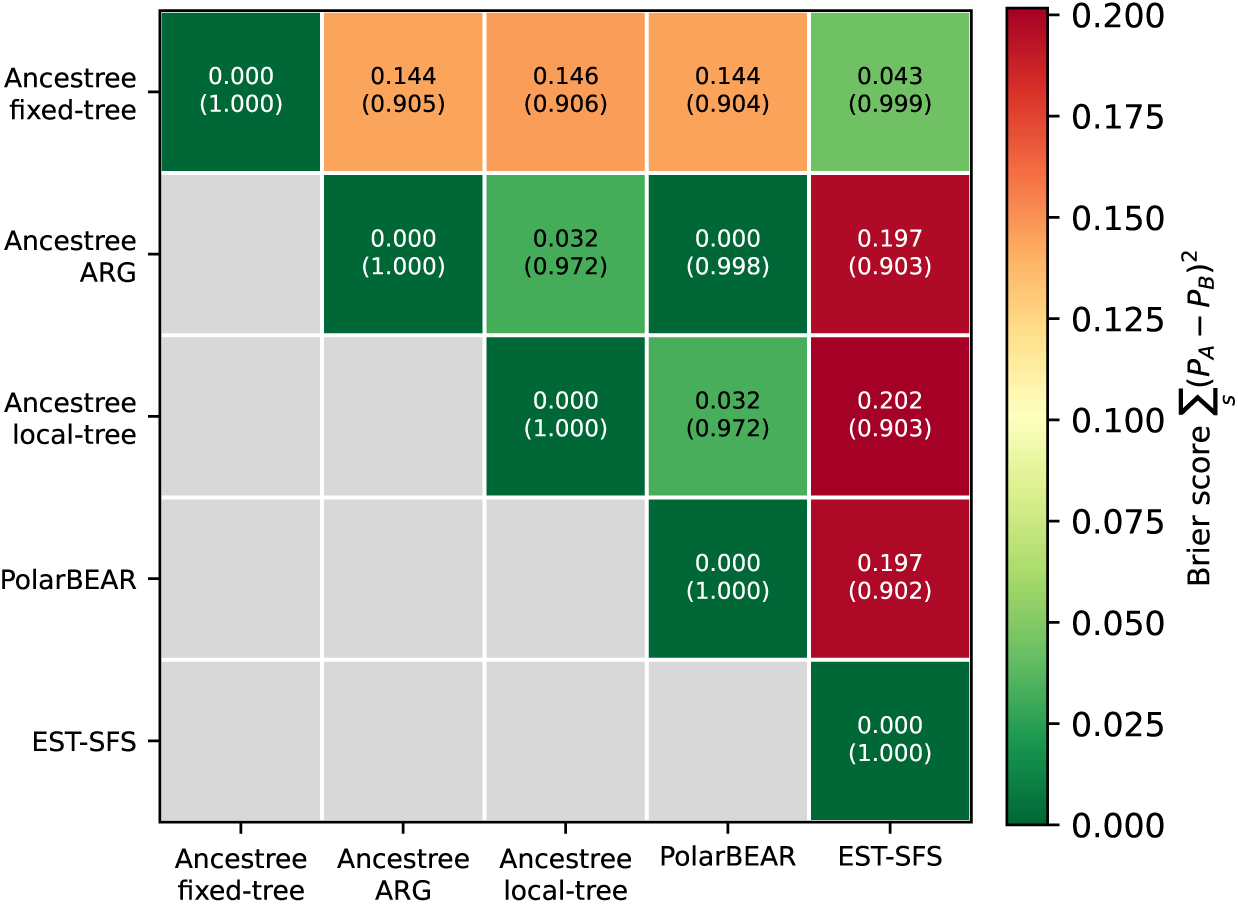
Pairwise agreement on the MSL chr1 panel. Over the positions both methods score, the top number (and the colour) is the Brier score 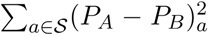 of the two per-site posteriors, averaged over sites, so lower is better, and the parenthetical is the MAP agreement—the fraction at which the two methods’ MAP ancestral allele coincides. The methods fall into two blocks: genealogy-based (`Ancestree`-ARG, `Ancestree`-Local-tree, `PolarBEAR`) and outgroup-based (`Ancestree`-VCF, `EST-SFS`), agreeing highly within each block and less across the two families.

#### Coverage

`Ancestree` scores every one of the 1,140,085 ingroup-polymorphic MSL SNPs, and `PolarBEAR`’s kernel reaches essentially the same set—it scores all but eight (1,140,077). `Ancestree` additionally scores the ingroup-monomorphic divergence sites.

#### Runtime

The fixed-tree and ARG passes are less expensive than either reference tool, and the local-tree HMM the most expensive—it scales quadratically in the number of samples (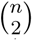 haplotype pairs)—so its serial cost runs to hours where the others take minutes (Table B4). The pairs are independent, so distributing them across cores annotates the whole chromosome in minutes.

**Table B4:**
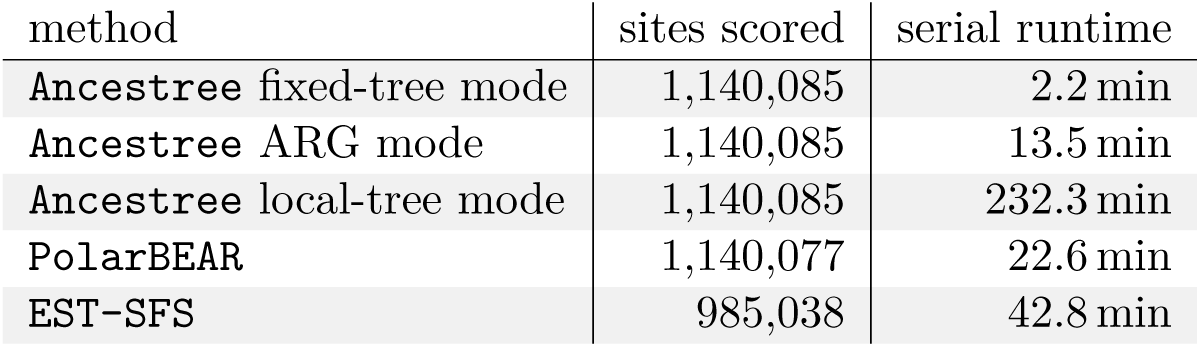
Coverage and serial runtime on the MSL chr1 panel. Sites scored per tool, and the serial cost of annotating the panel, taken for a sharded run as the sum of the shards’ own wall times so that every row is one definition. `Ancestree` and `PolarBEAR` reach essentially the whole SNP set, `EST-SFS` the biallelic sites at which both outgroups are called.

| method | sites scored | serial runtime |
| --- | --- | --- |
| **Ancestry** fixed-tree mode | 1,140,085 | 2.2 min |
| **Ancestry** ARG mode | 1,140,085 | 13.5 min |
| **Ancestry** local-tree mode | 1,140,085 | 232.3 min |
| **PolarBEAR** | 1,140,077 | 22.6 min |
| **EST-SFS** | 985,038 | 42.8 min |

### B.4 Frequency Prior versus Genealogy on Informative Sites

To gauge how much the local genealogy adds over the ingroup-frequency prior alone on real data, we restrict to the 792,598 MSL chr1 sites `PolarBEAR` marks as having an informative genealogy and compare each `Ancestree` mode against its per-site posterior and MAP estimate there (Table B5). ARG mode, reading the same `gamma-SMC` [Schweiger and Durbin, 2023] ARG, reproduces `PolarBEAR`’s MAP estimates exactly, and local-tree mode stays very close. Fixed-tree mode with no outgroups—which reduces to the Kingman ingroup weight alone (**Ingroup weights**)—still matches `PolarBEAR`’s MAP estimate at 96.6% of these sites (Brier score 0.0575). The genealogy is considerably more informative than the frequency prior where the derived allele is common, although such sites are rare (Appendix C.2).

**Table B5:** Agreement with PolarBEAR on its informative-genealogy sites. (792,598 MSL chr1 positions). Brier agreement and MAP agreement of each `Ancestree` mode against `PolarBEAR` (**Metrics**), both taken with `PolarBEAR`’s posterior in place of the one-hot truth. Fixed-tree mode with no outgroups is the Kingman ingroup weight alone.

| agreement with **PolarBEAR** | Brier score | MAP agreement |
| --- | --- | --- |
| ARG mode | $1.1 \times 10^{-7}$ | 1.0000 |
| local-tree mode | 0.0115 | 0.9926 |
| fixed-tree mode (Kingman prior, no outgroups) | 0.0575 | 0.9662 |

### B.5 Parsimony and Simple Baselines

Figure B2 reproduces the main-text heatmap (Figure 5) but sets the three `Ancestree` modes against additional inference modes: the `tsinfer`, `SINGER`, and `Relate` ARG-inference pipelines and two non-probabilistic baselines the likelihood kernel generalises—maximum parsimony and a majority-outgroup vote. The ARG-inference methods are compared directly in the main text (Section 3.3); this appendix gives the complete comparison across all methods and the full method details.

*Maximum parsimony* is the kernel’s own *µ →* 0 limit. Empirically its heatmap row is nearly indistinguishable from true-ARG mode: under infinite sites the integral is already dominated by that reconstruction, so taking *µ* to zero scarcely moves the posterior, and parsimony is essentially the kernel’s zero-rate limit.

*Majority outgroup* (MajorityOutgroupInference) takes the most frequent canonical allele among the outgroup tips as ancestral, breaking ties in favour of the closest outgroup, and by ad-hoc rule places a fixed confidence (here 0.95) on that allele with the remaining mass spread uniformly over the other three states (at *n*_out_ = 0 it falls back to the most frequent ingroup allele). It performs well wherever outgroups are informative, because neutral outgroups retain the ancestral allele, so their majority allele is the ancestral state in the absence of recurrent mutation. But its confidence is a fixed constant rather than a data-driven posterior, so it does not quantify uncertainty.

The `tsinfer` row places a realistic ARG-inference pipeline alongside: `tsinfer` requires the ancestral allele as input to build its ARG, so rather than the ground truth we orient it by a fixed-tree pass (using its *n*_out_ outgroups). It consumes that orientation as a fact rather than as a posterior, so it inherits the pass’s errors instead of correcting them. Local-tree mode takes no ancestral-allele input, yet reaches comparable genealogy quality (Figure 7) and accuracy from genotypes alone, with a far lighter pairwise-HMM build. At *n*_out_ = 0, `tsinfer` performs poorly: it must be oriented to build its ARG, and without an outgroup the orienting fixed-tree pass collapses to the SFS prior alone, which mis-orients the high-frequency-derived sites, after which the inferred genealogy confidently confirms the wrong orientation; it also cannot place the ingroup-invariant fixed sites without an outgroup, so its *n*_out_ = 0 fixed-class cells approach 1, the balanced Brier of an uninformative posterior, as they do for every method in Figure B2.

The two `SINGER` rows [Deng et al., 2025] add a full Bayesian ARG sampler, which infers the genealogy from the genotypes, does not require the ancestral allele as input, and supports uncertainty in the ancestral-allele probability (via the `-polar` flag). Both use the `SINGER` configuration of Section 3.3 (sample panel, Markov-chain averaging, and subsampled scoring window); both rows receive the same fixed-tree-oriented input, and the rows differ only in the confidence `SINGER` is told to place on that orientation: the *`SINGER`* row treats it as uninformative (`-polar` 0.5), while *`SINGER` (fixed-tree)* takes it as near-certain (`-polar` 0.99). The unoriented *`SINGER`* row (`-polar` 0.5) is comparable to local-tree mode and more robust than fixed-tree mode under strong incomplete lineage sorting. The *`SINGER` (fixed-tree)* row performs comparably, the two differing by under 0.01 in every site class. `SINGER` aborts deterministically on particular (chunk, seed) combinations, and such seeds are skipped and replaced so that every cell averages over the same number of chains. Under purifying selection at *n*_out_ = 1 it aborts at nearly every seed. The affected chunks are excluded from the results, which then average over fewer sites.

The `Relate` row places a second ancestral-allele-requiring ARG builder alongside `tsinfer`: it is oriented by the same fixed-tree pass (using its *n*_out_ outgroups), and it likewise performs poorly at *n*_out_ = 0 and, given that orientation, inherits its error rather than improving on it.

The fixed-tree mode (Kingman weight) and fixed-tree (adaptive) rows are near-indistinguishable across every scenario, including the purifying one (Figure B2): the fitted per-bin weights *ω_i_* converge to their neutral Kingman values wherever the prior is exercised (**Priors**).

**Figure B2:**
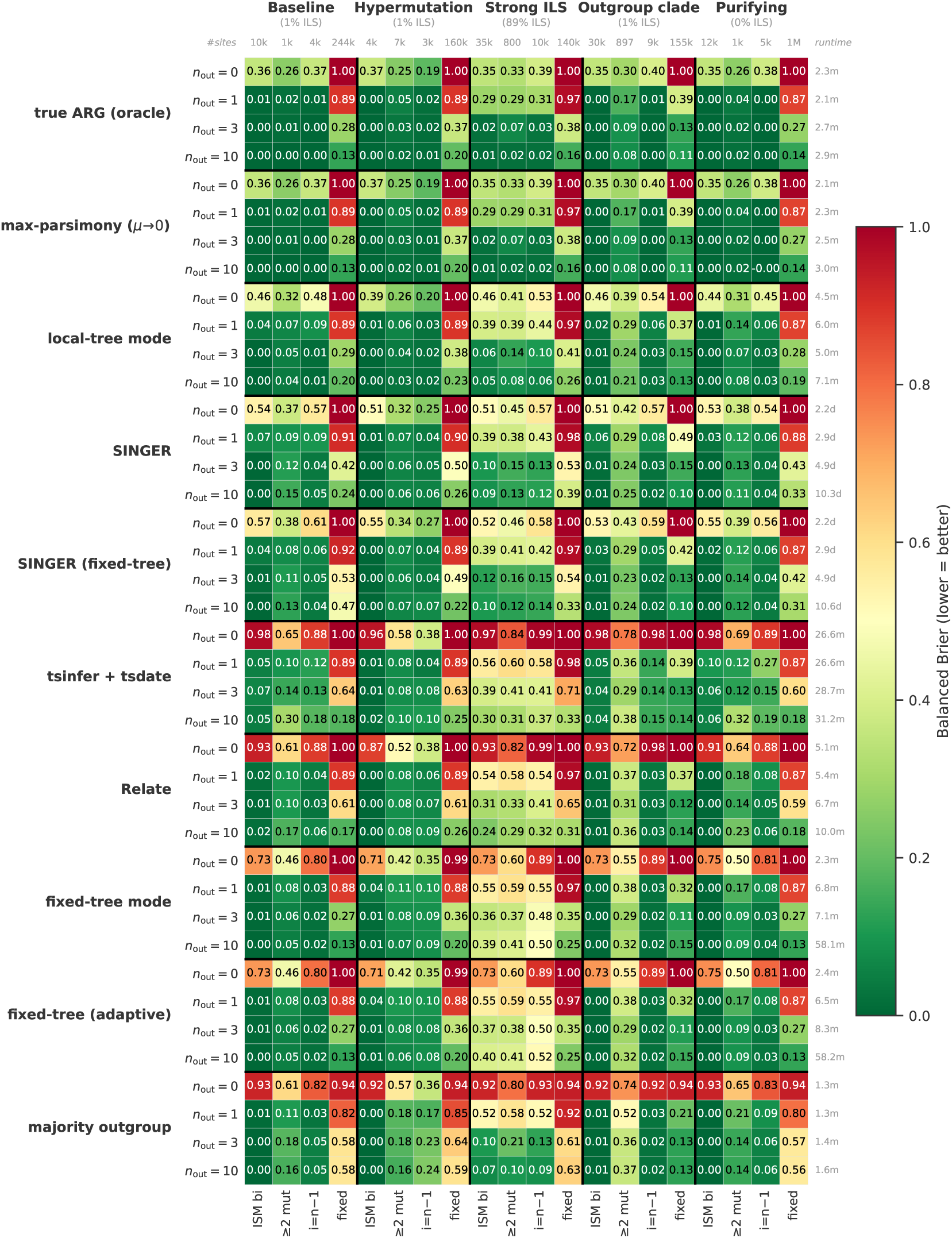
Simple baselines vs. the full likelihood across scenarios, *n*_out_, and site classes. Balanced Brier (lower = better; constructed as in Figure 5) for ten method blocks × *n*_out_ ∊ {0, 1, 3, 10} over the five scenarios, each split into four site classes; the right strip gives per-row runtime. The first two rows (true ARG, max-parsimony) run on the true local trees and so use oracle genealogy; the rest infer the genealogy or use none.

## C Extended Benchmarks and Robustness

### C.1 Where the Ancestral State is Read

The kernel evaluates Felsenstein’s recursion toward a single node, the focal node, and reports the posterior there. Which node that is must be stated explicitly, rather than taken from the tree’s rooting. Two nodes are natural. The ingroup most recent common ancestor is the node an unfolded site-frequency spectrum is typically polarised about: the ancestral state of an ingroup polymorphism is the state at the MRCA of the ingroup sample. The panel root is the most recent common ancestor of every sample, deeper than whenever outgroups are included, and is the informative node for an allele the ingroup has fixed. Here we vary the focal node continuously between the two anchors. We parameterise it as the fraction of the path from to the deepest node the panel provides, which is scale-free and hence comparable across panels whose divergences differ. Figure C1 grades each position against its own node, Figure C2 against the node each site class names.

**Figure C1:**
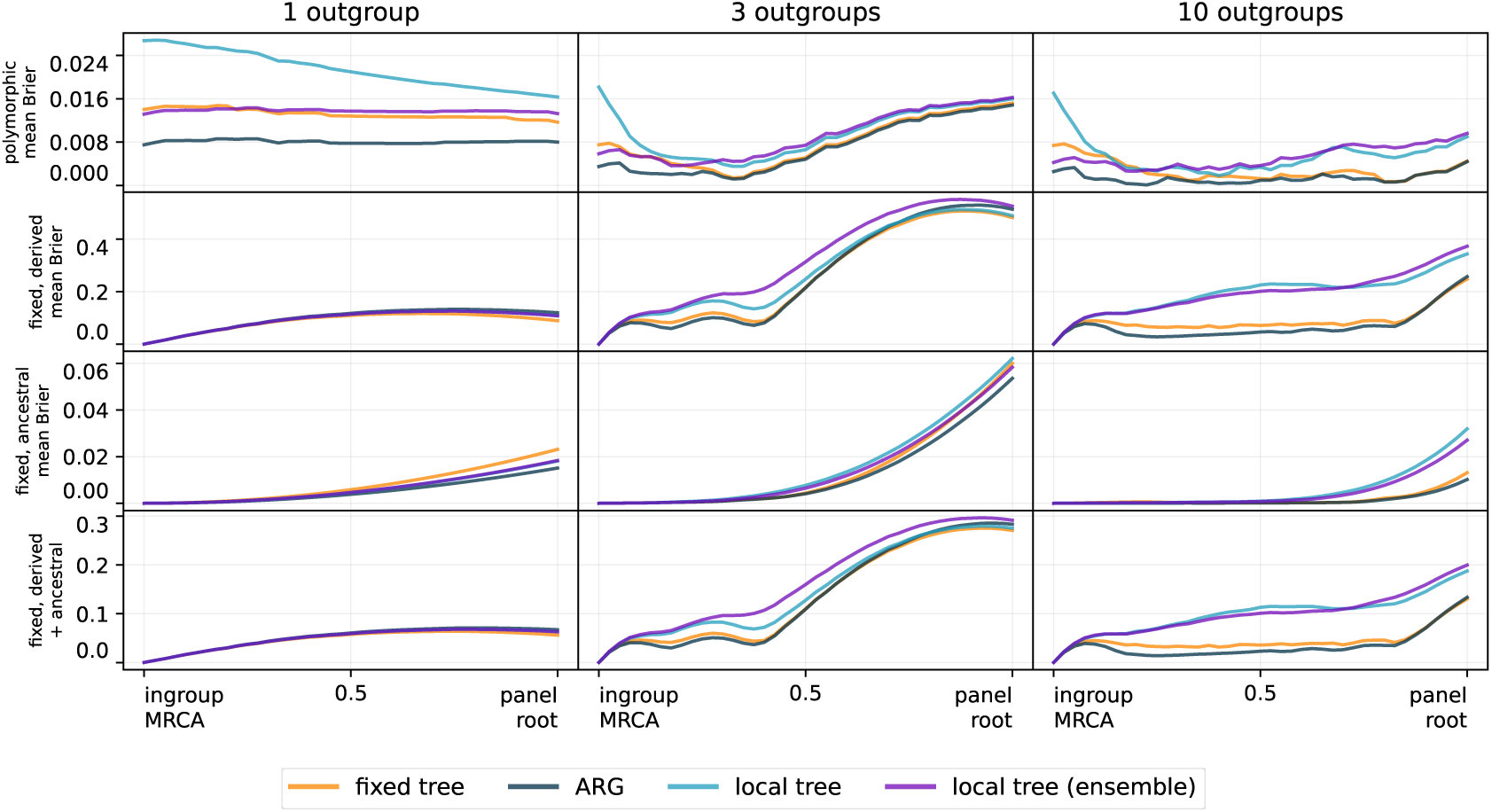
**Accuracy against the focal node**, varied from the ingroup MRCA to the panel root, for one, three and ten outgroups and for all three inference modes. Each panel reports the mean Brier score, for which lower is better, with the ingroup-polymorphic sites in the upper row and the ingroup-monomorphic sites below, on a scale shared within each row. The bottom row weights the two ingroup-monomorphic classes equally despite their unequal site counts, giving a class-balanced mean over the sites the ingroup has fixed.

Figure C1 varies that fraction on the baseline coalescent scenario, separately for the sites that are polymorphic within the ingroup and those that are not. Each position is graded against the simulated state at that same position, so each point measures how well the state at that node is recovered. The positions are therefore not comparable as answers to one question: moving up the path changes the question being asked, and a deeper node is both further from the polymorphism and better supplied with outgroup evidence. Because each position in Figure C1 is graded against its own node, the figure cannot show what is lost by reading in the wrong place.

Figure C2 holds the question fixed instead, grading each site class at the node its definition names, ingroup polymorphism at and sites the ingroup has fixed at the panel root, while the focal node varies along the same path. The resulting cost is asymmetric. Reading an ingroup polymorphism deeper than grows more expensive with each outgroup, as the divergence branch separates the reading from the polymorphism. Reading a fixed difference at is more costly again, driving the Brier score to its maximum, since the posterior there merely restates the allele the ingroup has fixed.

**Figure C2:**
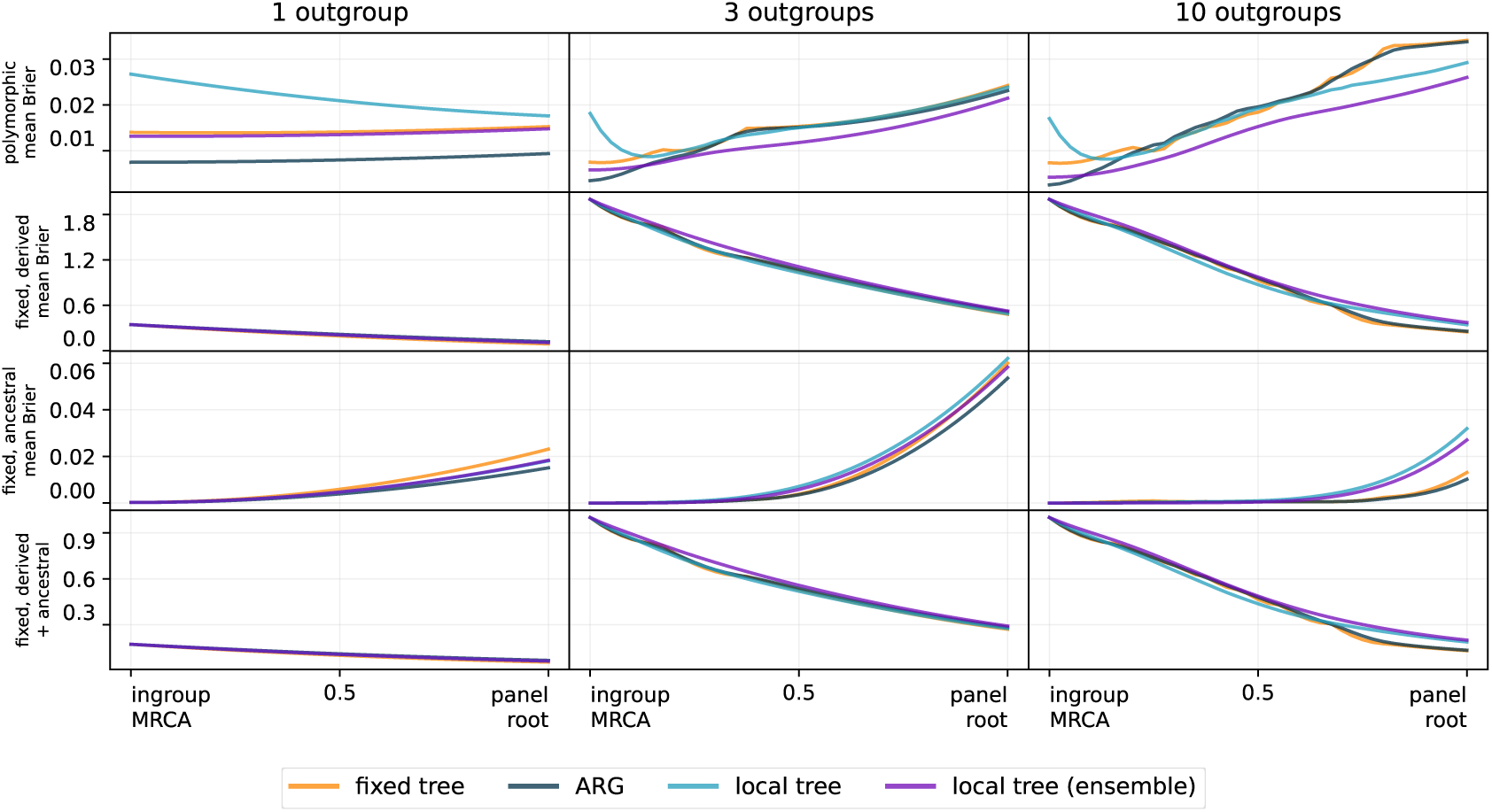
The cost of reading at the wrong node. The focal node runs from the ingroup MRCA to the panel root, for one, three and ten outgroups. Each of the upper three rows grades one site class at the node its definition names, so that, unlike Figure C1, the vertical axis is comparable along the path. The bottom row weights the two ingroup-monomorphic classes equally despite their unequal site counts. Each panel reports the mean Brier score, for which lower is better.

For ingroup-polymorphic sites accuracy varies only mildly over the path, and is best at *I* itself, the node their definition names. Sites the ingroup has fixed for the derived allele are best read at the panel root, and those fixed for the ancestral allele at, so no single node suits both.

### C.2 Ingroup-Size Saturation

To test whether ARG ingroup-only mode improves as the ingroup grows, we take one `msprime` ARG (*n* = 1000 haploid, *L* = 10^6^, *µ* = 1.25 × 10^−8^, *N_e_* = 3 × 10^4^, no outgroups), score it with `ARGBasedInference` in pure ingroup-only mode, and in each cell subsample the ingroup to *n ∊* {20, 50, 100, 200, 500, 1000}, projecting every site onto the *n* = 20 reference grid by hypergeometric down-sampling (Figure C3). Per-bin accuracy turns out to be flat across ingroup sizes: once the ingroup is large enough to resolve the local topology, additional samples add little further information about which allele is ancestral. With no outgroup, the fixed-tree reference’s posterior collapses to the pure Kingman SFS prior, which assigns *P* (allele of count *a* is ancestral) = *a/n*, so its MAP is right only when the derived allele is the minor one (*i < n/*2) and falls to zero on high-frequency-derived sites, while its Brier score 2(*i/n*)^2^ grows quadratically with *i*, the unfolded derived-allele count on the reference grid. That reference is the floor a genealogy-free method reaches, and reading the local tree improves on it across the whole interior, most of all in the high-frequency-derived bins, which the prior alone mis-assigns.

**Figure C3:**
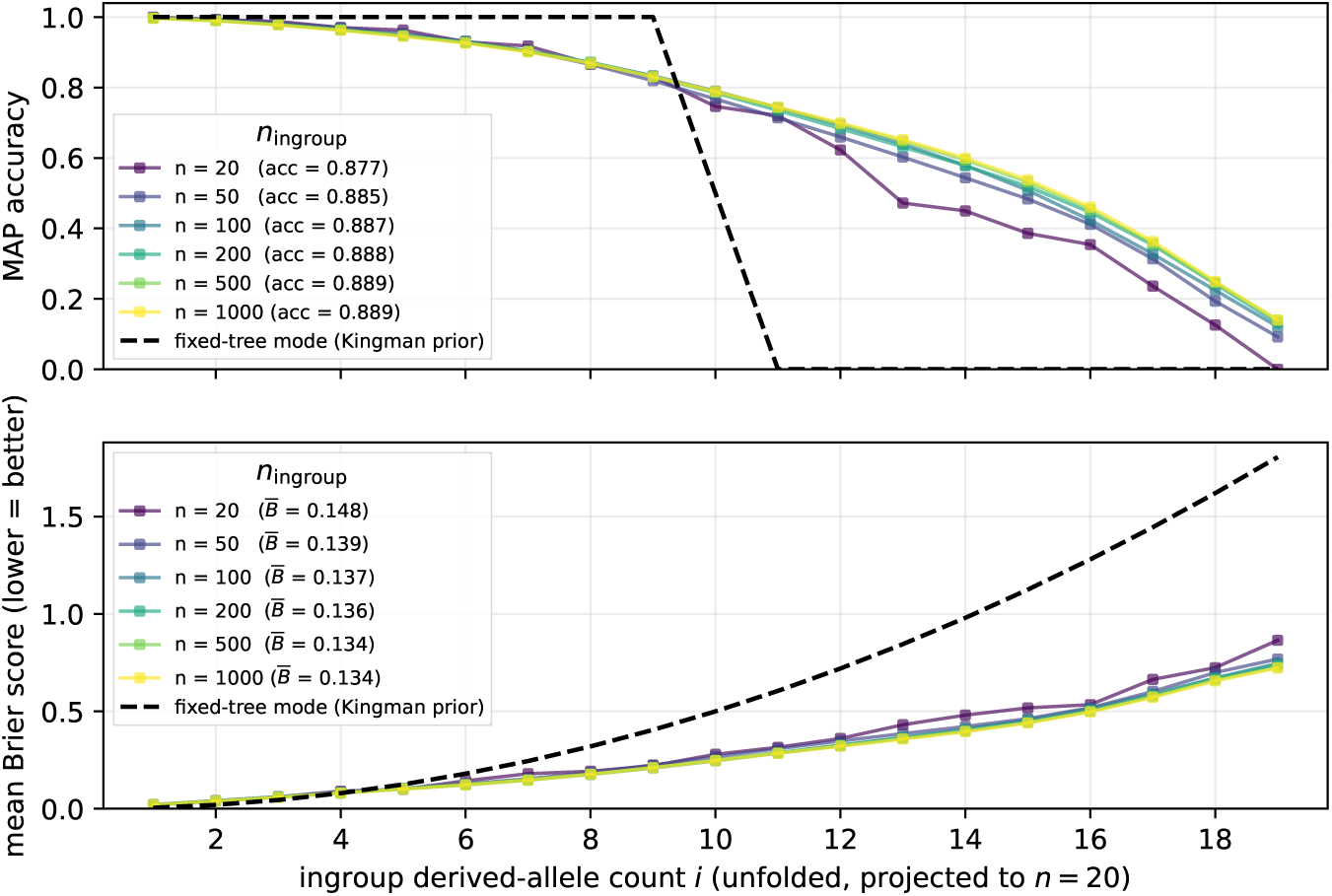
**Per-bin MAP accuracy and mean Brier score across ingroup sizes in ARG ingroup-only mode**, over the unfolded SFS projected to *n* = 20; one line per *n*, with the dashed black line the no-outgroup fixed-tree (pure Kingman-prior) reference.

### C.3 Prior Choice under Demographic Distortion

Fixed-tree mode can weight the candidate ancestral states by the ingroup allele frequencies (**Ingroup weights**). We compare the two such weights against the root prior applied alone: the Kingman weight, the neutral coalescent expectation *P* (root = *a*) = *n_a_/n*; the adaptive weight, which fits one polarisation probability per frequency class from the data; and the stationary root prior applied without any ingroup weight, which is uniform over ancestral states under JC69 and carries no orientation information (**Priors**). We evaluate all three under three demographies that share the outgroup ladder but differ in the recent history of the ingroup: a constant-size baseline, and a recent expansion and contraction in which the ingroup changes size exponentially by a factor of ten over the most recent 1.2 10^5^ generations—growing from a tenth of its present size, or declining from ten times it—and is held constant before that. The expansion enriches the low-frequency tail and the contraction the high-frequency derived tail. Each setting’s posterior-weighted (expected) unfolded SFS (Section 3.5) is scored against the ground truth over the ingroup-polymorphic sites, with no, one, and three outgroups (Figure C4).

**Figure C4:**
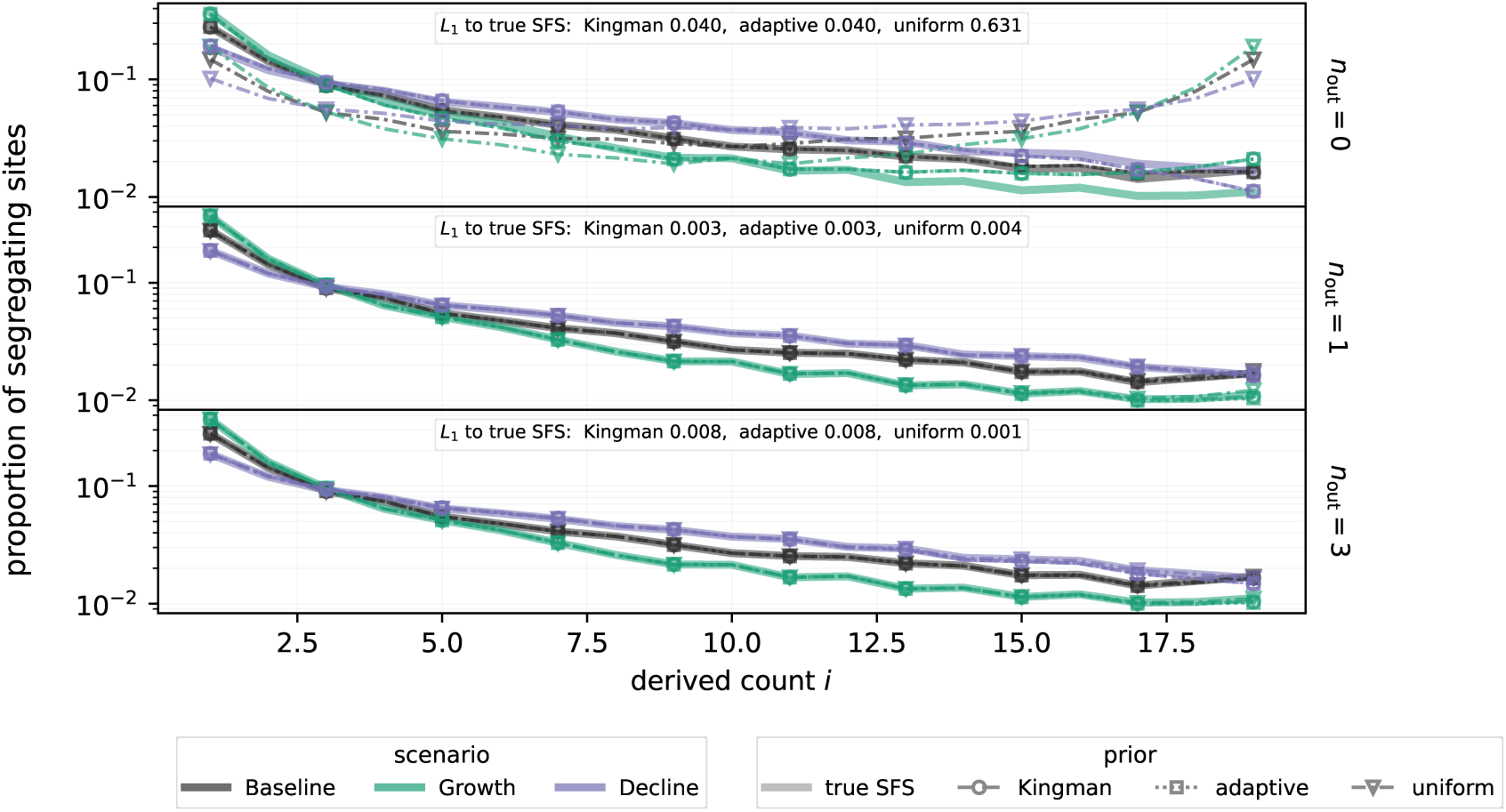
The ingroup weight governs ancestral-state recovery without an outgroup. Its influence falls off rapidly as outgroups are added. Posterior-weighted (expected) unfolded SFS under the Kingman and adaptive ingroup weights and under the stationary root prior alone, against the ground-truth spectrum, with no, one, and three outgroups (top to bottom), for three demographies. With no outgroup the posterior is the normalised weight and the reconstructions diverge: the stationary prior carries no orientation information and folds the spectrum symmetrically about its midpoint, while the count-based Kingman weight reconstructs an approximately neutral spectrum regardless of the truth. A single outgroup fixes the orientation through the Felsenstein likelihood and the three converge onto the ground truth; with three the agreement is complete.

Without an outgroup the weight alone determines the reconstruction, and none recovers a distorted truth reliably: the stationary prior folds the spectrum symmetrically, while the Kingman weight reproduces a near-neutral one irrespective of the demography. Outgroups suppress this dependence rapidly, since the outgroup allele is at most sites simply the ancestral allele, so that one outgroup fixes the orientation through the Felsenstein likelihood and the three settings converge onto the truth, becoming indistinguishable with three. The adaptive weight offers no advantage at any outgroup count, as it cannot be fitted without an outgroup and carries too little influence with one. From one outgroup onward the stationary prior alone reconstructs the distorted spectrum at least as faithfully as the Kingman weight, and more faithfully with three, imposing no neutral expectation for the data to override. Dropping the ingroup weight is therefore the more accurate choice here, although the ordering is dataset-dependent.

### C.4 Substitution-Model Misspecification

To ascertain how much choosing the wrong substitution model biases branch-length inference in fixed-tree mode, and to check whether ARG mode is affected likewise, we run the five models `Ancestree` provides—JC69, K2, F81, HKY, GTR—against simulated ground truth. A single `msprime` ARG (*n* = 20, three outgroups at (3, 9, 15)×10^5^ generations, *L* = 10^7^, *µ* = 1.25×10^−8^) is mutated under HKY with a 4:1 transition/transversion bias and an AT-rich stationary distribution (*κ* = 4, *ϕ* = (0.3, 0.2, 0.2, 0.3)). JC69 is then misspecified on both axes, K2 on base composition and F81 on the transition bias. Neither form of misspecification affects inference. The fitted divergences agree with one another to within 0.4 percentage points and per-site accuracy is unchanged (ARG 0.993, fixed-tree 0.984), the choice of substitution model being negligible at realistic outgroup divergence times (Table C1).

**Table C1:** Substitution-model misspecification. Fixed-tree ingroup-to-outgroup divergences (sums of ladder branches, not individual rates *K_i_*) fitted under each model, as percentage deviations from the truth (0.0064, 0.0193, 0.0322) substitutions per site.

| model | div<sub>1</sub> | div<sub>2</sub> | div<sub>3</sub> | max $\vert \cdot\vert $ |
| --- | --- | --- | --- | --- |
| JC69 | +3.4% | +1.0% | +0.2% | 3.4% |
| K2 | +3.5% | +1.2% | +0.5% | 3.5% |
| F81 | +3.4% | +1.0% | +0.3% | 3.4% |
| HKY | +3.6% | +1.2% | +0.6% | 3.6% |
| GTR | +3.5% | +1.2% | +0.6% | 3.5% |

### C.5 Site-Count Requirements

To establish how many polymorphic sites `FixedTreeInference` needs before its free parameters stabilise, we take one `msprime` ARG (*n* = 20, three nested-split outgroups at (3, 9, 15) × 10^5^ generations, *N_e_* = 3 × 10^4^, *µ* = 1.25 × 10^−8^, JC69) and sub-sample *L*_sites_ ∊ {10^2^, 3 × 10^2^*,...,* 10^5^} polymorphic sites (Figure C5); The number of target sites the optimiser is told the panel holds, n_target_sites, scales with *L*_sites_, so that the polymorphism rate it sees stays constant. On the fixed-tree side (left subplot), the mean relative deviation of the fitted ladder rates from the full k-site reference decays as 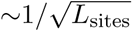, from 15–32% at *L*_sites_ = 100 to 0.5–1% at 10^5^. Adding outgroups does not raise the per-parameter cost—the *n*_out_ = 3 curve sits below *n*_out_ = 1, as the extra tips tend to concentrate the per-site likelihood faster than the extra ladder rates add variance. The adaptive weight (**Adaptive weight**) instead learns a per-SFS-bin root-state weight *ω_i_* from the data, replacing the fixed neutral Kingman value, with each site projected onto an *n*_sub_-haplotype sub-sample by hypergeometric weighting. Recovery of these *ω_i_* degrades at small *L*_sites_ as *n*_sub_ grows (more bins, fewer sites each), though with a diminishing penalty. In practice, a few thousand polymorphic sites suffice for reliable recovery.

**Figure C5:**
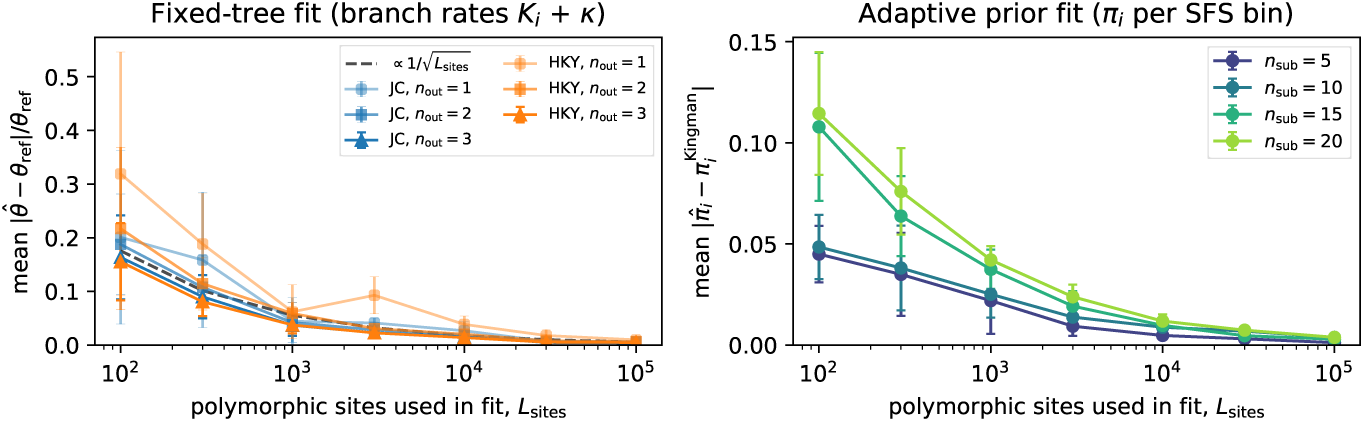
Parameter-recovery error vs. polymorphic-site count. *Left:* fixed-tree ladder recovery (mean absolute relative deviation of the fitted *K_i_*, and *κ* under HKY, from the full-data reference), one line per (model, *n*_out_ ∊ {1, 2, 3}. *Right:* adaptive-prior per-bin recovery (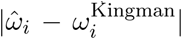 over fittable bins; centre bin excluded), one line per ingroup sub-sample *n*_sub_ ∈ {5, 10, 15, 20}. Error bars ±1 s.d. over five seeds.

### C.6 Spectra of the Simulation Scenarios

Figure C6 shows the polymorphic SFS of each of the five scenarios of Section 3.2. The spectra share the expected coalescent decay but differ markedly in polymorphic budget—by more than an order of magnitude between the hypermutation and purifying-selection scenarios—with SLiM also the most rare-variant-skewed.

**Figure C6:**
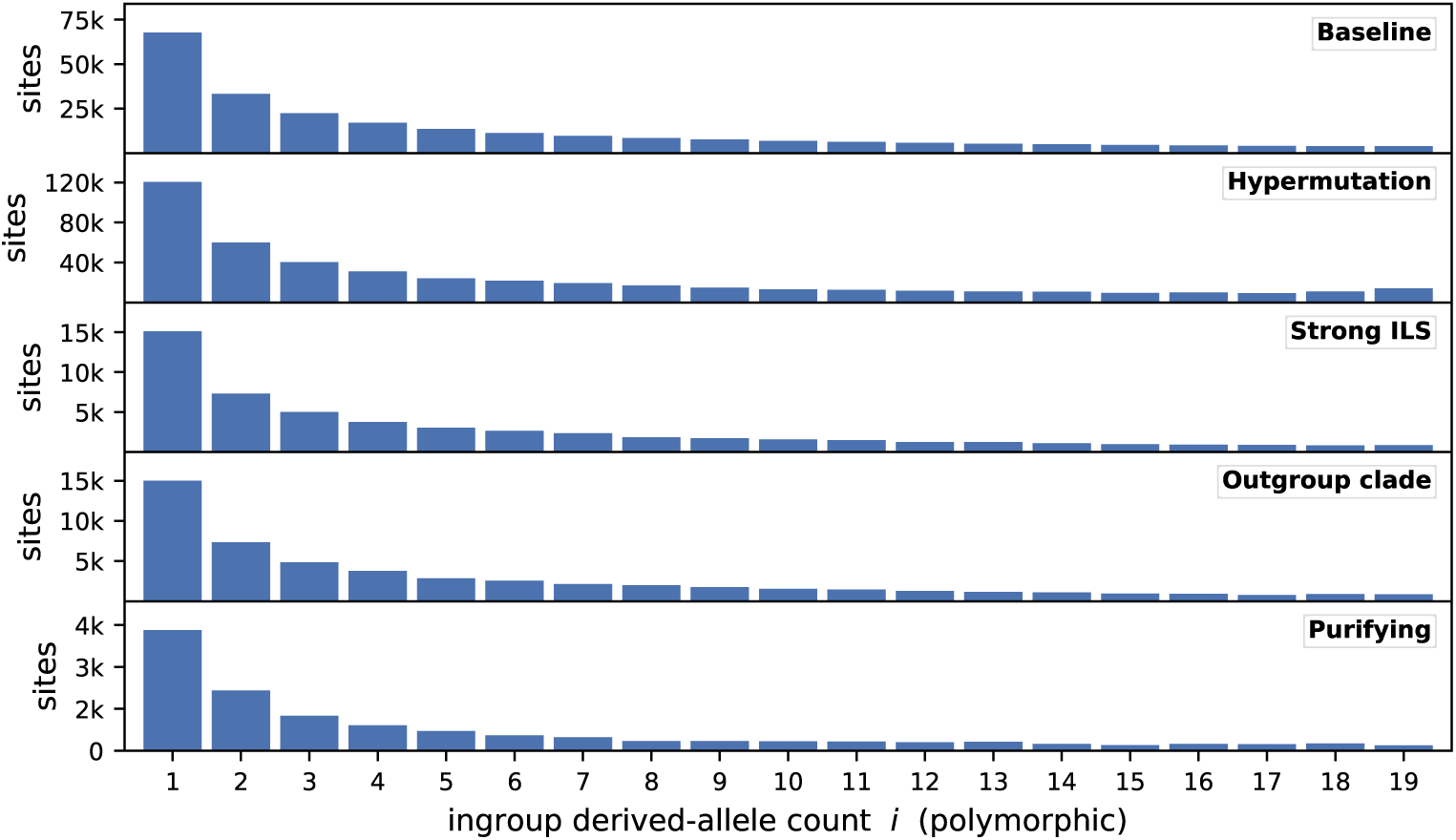
Polymorphic SFS per simulation scenario. Counts of segregating sites by unfolded ingroup derived-allele count *i* ∈ {1*,..., n* − 1}.

### C.7 Expected SFS Recovery

Figure C7 compares the true, MAP and posterior-weighted ingroup spectra across the scenarios and outgroup counts.

**Figure C7:**
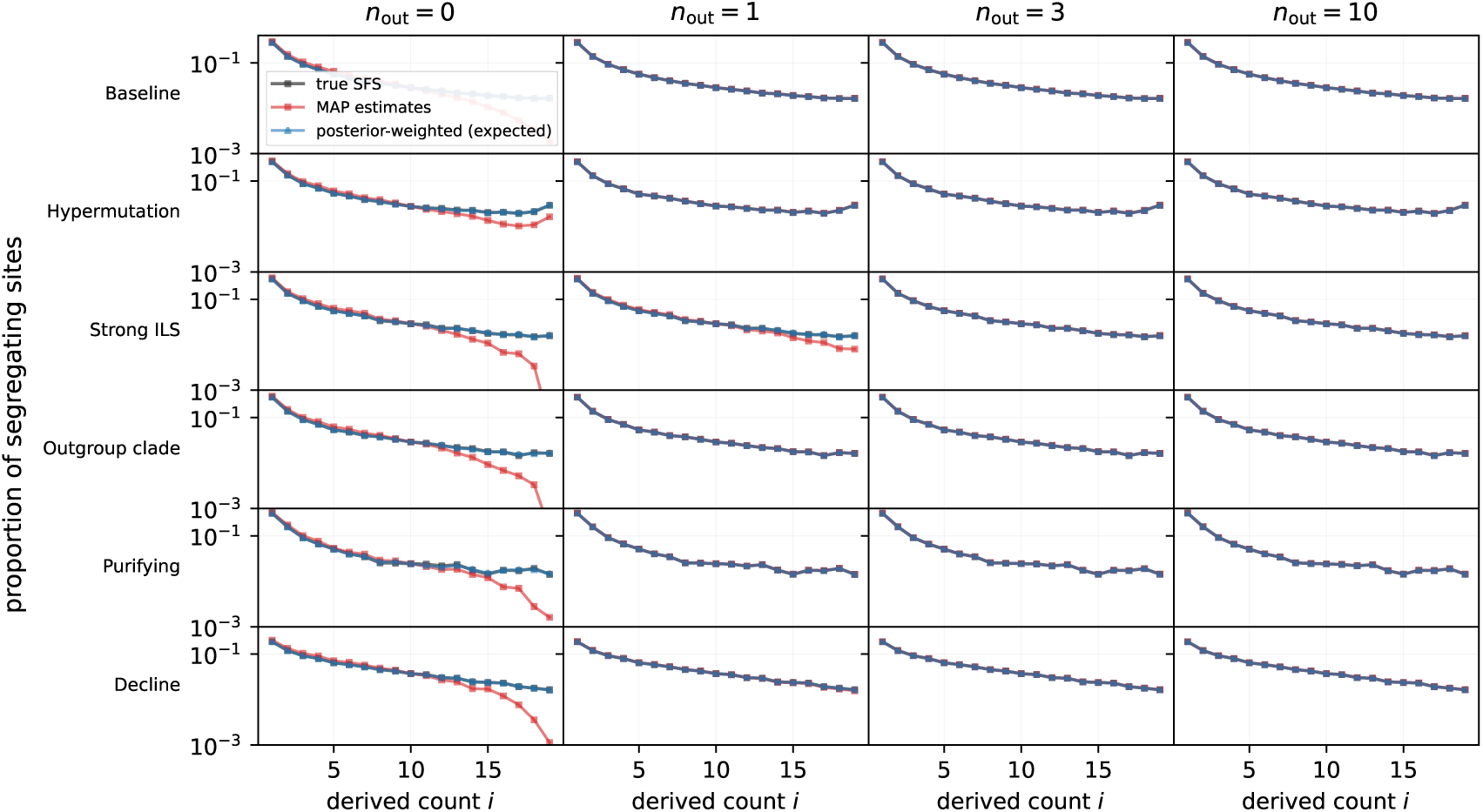
The posterior-weighted (expected) spectrum recovers the unfolded SFS across scenarios and outgroup counts. True, MAP, and posterior-weighted (expected) ingroup spectra for ARG-mode posteriors, arranged by scenario (rows) and outgroup count *n*_out_ (columns). The rows comprise the robustness scenarios of Section 3.2 together with a population-decline scenario, whose enriched high-frequency tail is where MAP attribution is most vulnerable. Committing to a MAP estimate depletes the high-frequency tail, most severely at *n*_out_ = 0; the posterior-weighted spectrum remains close to the truth, and the residual discrepancy is eliminated once one or more outgroups are supplied.

### C.8 Local-Tree Mode with Outgroups

The local-tree analysis in the main text (Section 3.4) uses an ingroup-only panel (*n*_out_ = 0), the worst case for ancestral-allele recovery, since without an outgroup every error in the reconstructed local trees passes straight into the inferred ancestral state. Here we repeat the same analyses with outgroups added to that panel, simulating an outgroup ladder under the baseline coalescent demography (Section 3.2) with the outgroup populations diverging from the ingroup lineage at 3 10^5^, 9 10^5^ and 1.5 10^6^ generations (about 10, 30 and 50 × *N_e_*), the *n*_out_ = 1 panels using the most closely related outgroup and the *n*_out_ = 3 panels all three. The outgroups enter local-tree mode as additional haplotypes in the inferred local tree, which the kernel weights no differently for being outgroups, their labels serving only to place the focal node at the ingroup MRCA. The Brier score is computed over the ingroup-polymorphic sites, so the panels are directly comparable.

**Figure C8:**
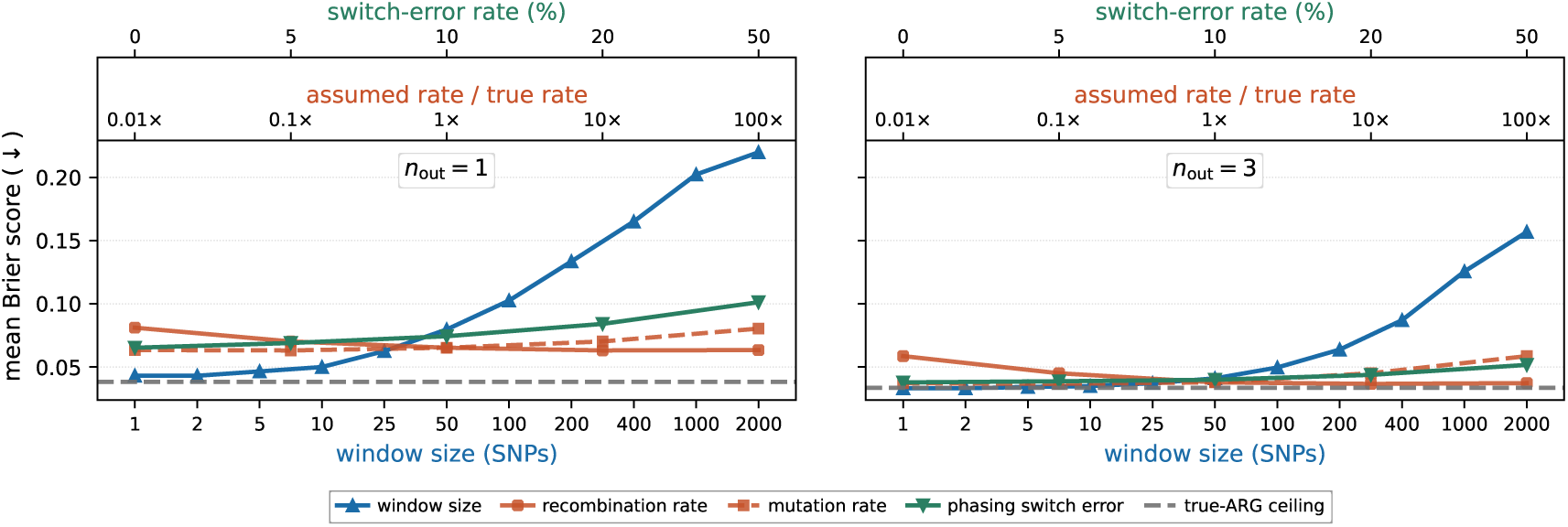
**Window size, rates and phasing with outgroups**, for *n*_out_ = 1 (left) and *n*_out_ = 3 (right) on a shared Brier axis, laid out as the ingroup-only Figure 6. Each input is varied against a common extent and read off its own colour-matched axis: the window size below, the assumed recombination and mutation rates as multiples of the truth above, and the phasing switch error above that, up to *s* = 0.5, random phase. The dashed line is the true-ARG ceiling, and every curve is produced with the ensemble of *M* = 128 sampled genealogies.

**Figure C9:**
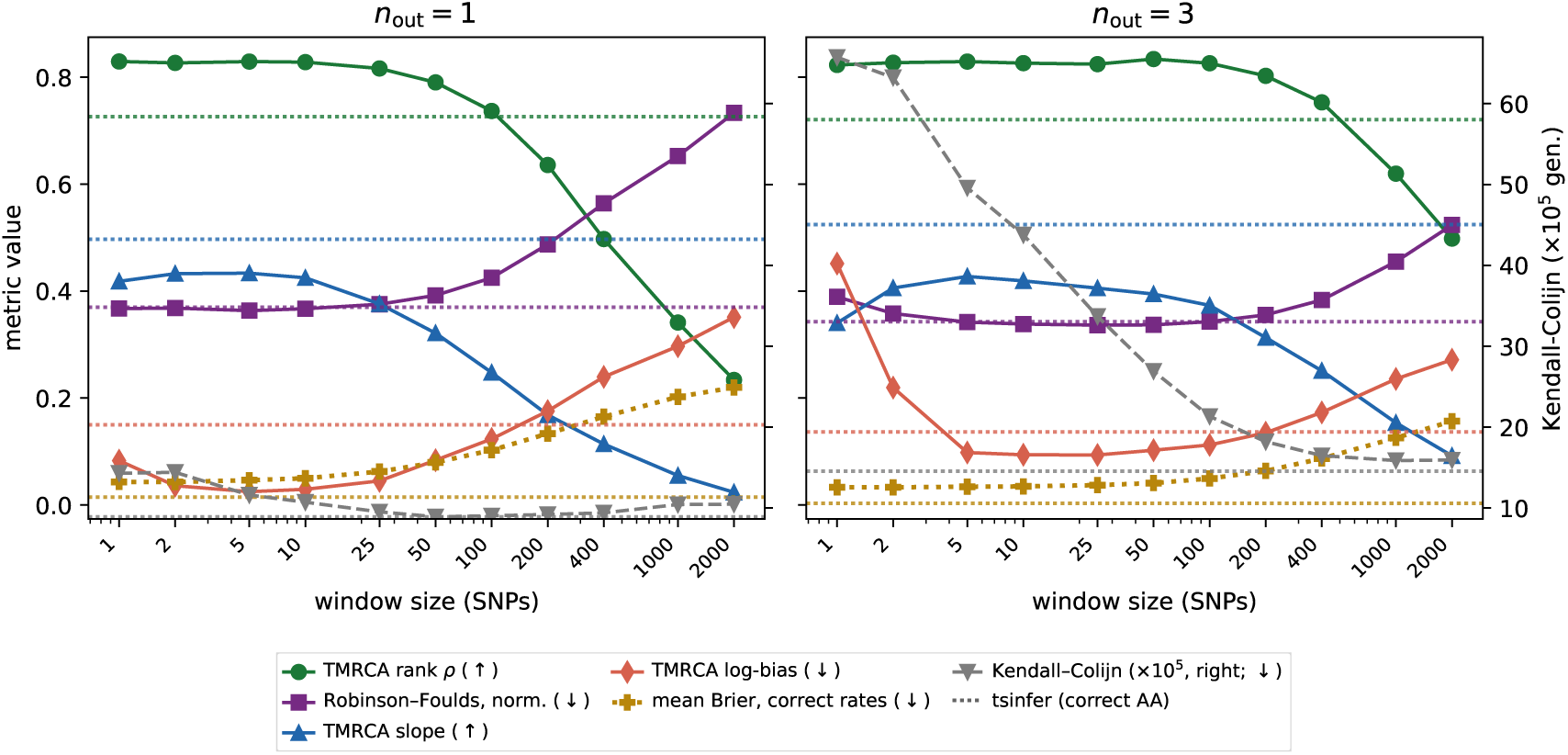
**Genealogy recovery with outgroups**, for *n*_out_ = 1 (left) and *n*_out_ = 3 (right) on a shared metric axis; the counterpart of the ingroup-only Figure 7 on a 20 Mb sim. TMRCA rank correlation, normalised Robinson–Foulds and Kendall–Colijn distances and TMRCA calibration against window size, with the `tsinfer` (correct-ancestral-allele) reference dashed. All metrics are computed from the posterior-mean tree, except the mean Brier score, which comes from the *M* = 128 ensemble. Genealogy recovery is essentially unchanged from the ingroup-only case, so the accuracy gain comes from the outgroup signal rather than from better tree reconstruction. The larger Kendall–Colijn distance at *n*_out_ = 3 reflects the deeper outgroup tree, the metric being absolute and branch-length-aware.

A single outgroup already changes the picture qualitatively (Figures C8 and C9). The mean Brier falls significantly at the reference window relative to the ingroup-only case, and by more at the narrowest windows. The rate-misspecification and phasing sensitivities, already mild without outgroups, become negligible. The reason is that once one or more outgroups are present, the ancestral state is determined primarily by the ingroup–outgroup topology—a strong, coarse signal that does not require resolving the fine ingroup genealogy—so the window size, the assumed rates and the phasing, which act only on that fine structure, have little influence. The genealogy-recovery metrics are themselves essentially unchanged from the ingroup-only case, since the deep outgroup tips are easily placed; the improvement comes from the added ancestral-state information, not from a better-reconstructed genealogy.

### C.9 Mis-Orientation and the Inferred Genealogy

`tsinfer` and `Relate` take the ancestral allele as input, so a mis-oriented site conditions the genealogy each tool builds. On the ingroup-only genealogy simulation of Section C.8, a 20 Mb neutral coalescent panel of 20 ingroup haplotypes (mutation rate *µ* = 1.25×10^−8^ and recombination rate *r* = 10^−8^ per site per generation, effective population size *N_e_* = 3 × 10^4^), we build two ARGs per tool from identical genotypes, one oriented by the simulated ancestral state and one mis-orienting 5% of the 102,913 biallelic sites. A site carrying *i* of 20 derived alleles is mis-oriented with probability proportional to *i/*20, since mis-orientation tends to be more likely for high-frequency derived alleles. Figure C10 scores the oriented and mis-oriented ARGs against the true local trees, binned by the distance from each site to the nearest mis-oriented one. Recovery degrades on every metric, most at the mis-oriented sites themselves, and most of that degradation is gone within a few hundred base pairs.

**Figure C10:**
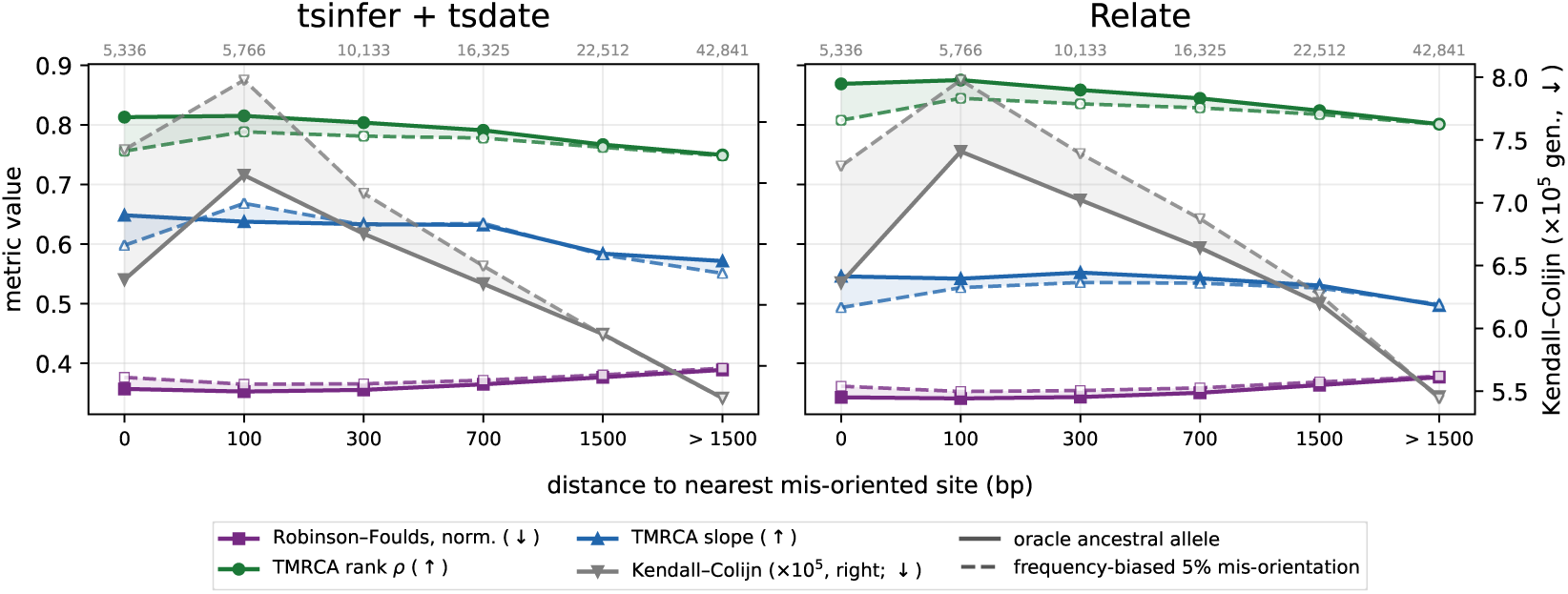
**Genealogy recovery against distance to the nearest misoriented site**, for `tsinfer` with `tsdate` and for `Relate`, built from identical genotypes under the simulated ancestral state (solid line) and under a frequency-biased 5% mis-orientation (dashed line); shading is the difference. Each metric’s better direction is marked in the legend, and site counts per bin are given above each panel.

### C.10 Ensemble Size

Local-tree mode marginalises each window’s posterior over an ensemble of genealogies drawn from the pairwise-HMM posterior rather than scoring a single point estimate (Section 2.4). Figure C11 varies the number of draws over the five robustness scenarios of Section 3.2. The scenarios saturate at different rates. Most plateau within a few tens of draws, while purifying selection alone still gains materially at the largest ensembles. Runtime scales well with the ensemble size and falls close to proportionally with the number of workers (Figure C12). Collapsing the posterior to a point estimate narrows the marginal distribution of pairwise coalescence times, which the draws restore (Figure C13).

**Figure C11:**
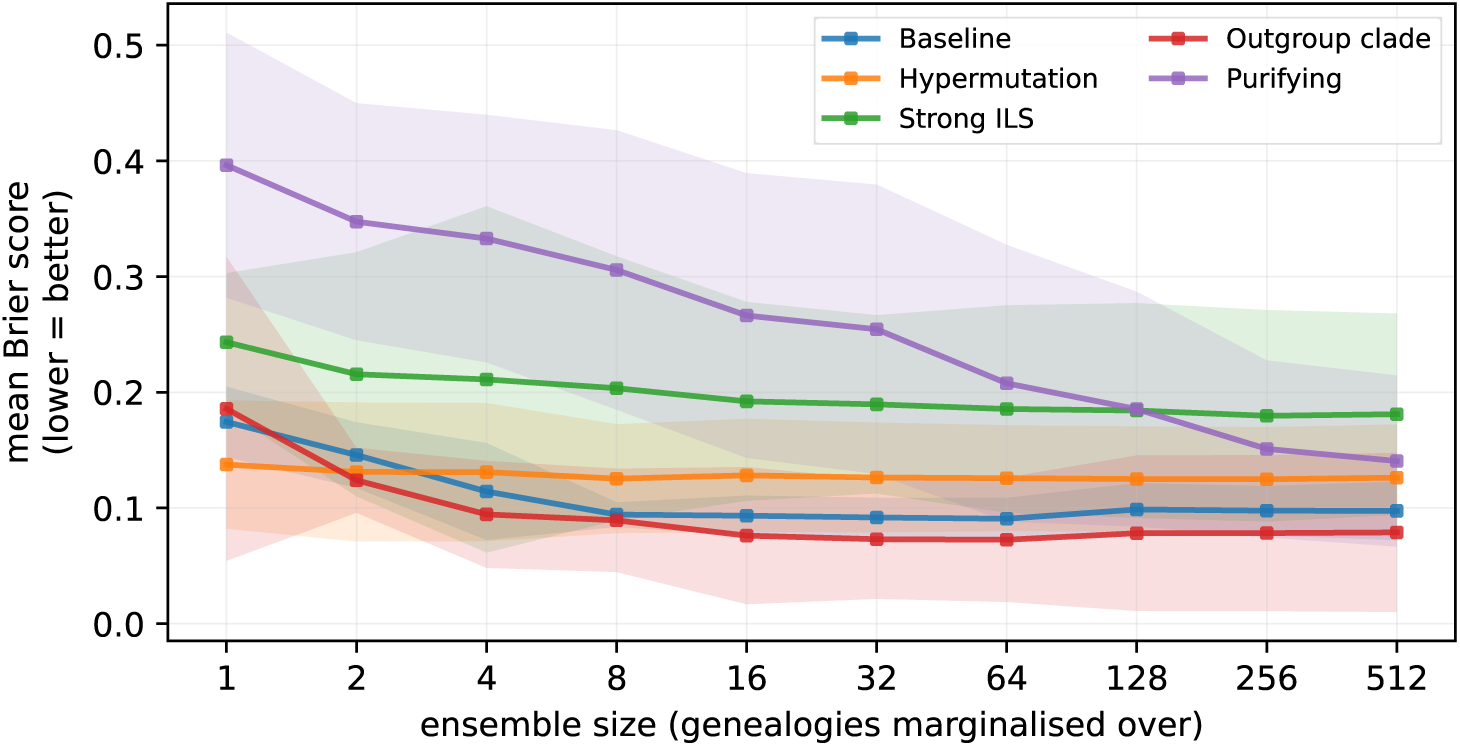
**Ground-truth recovery against ensemble size**, for the five robustness scenarios at *n*_out_ = 3. Mean Brier score against the simulated ancestral state, lower is better; points are the mean over independent chunks and the band is a 95% confidence interval of that mean (three chunks per scenario, ten for purifying selection). The x-axis doubles at each step.

**Figure C12:**
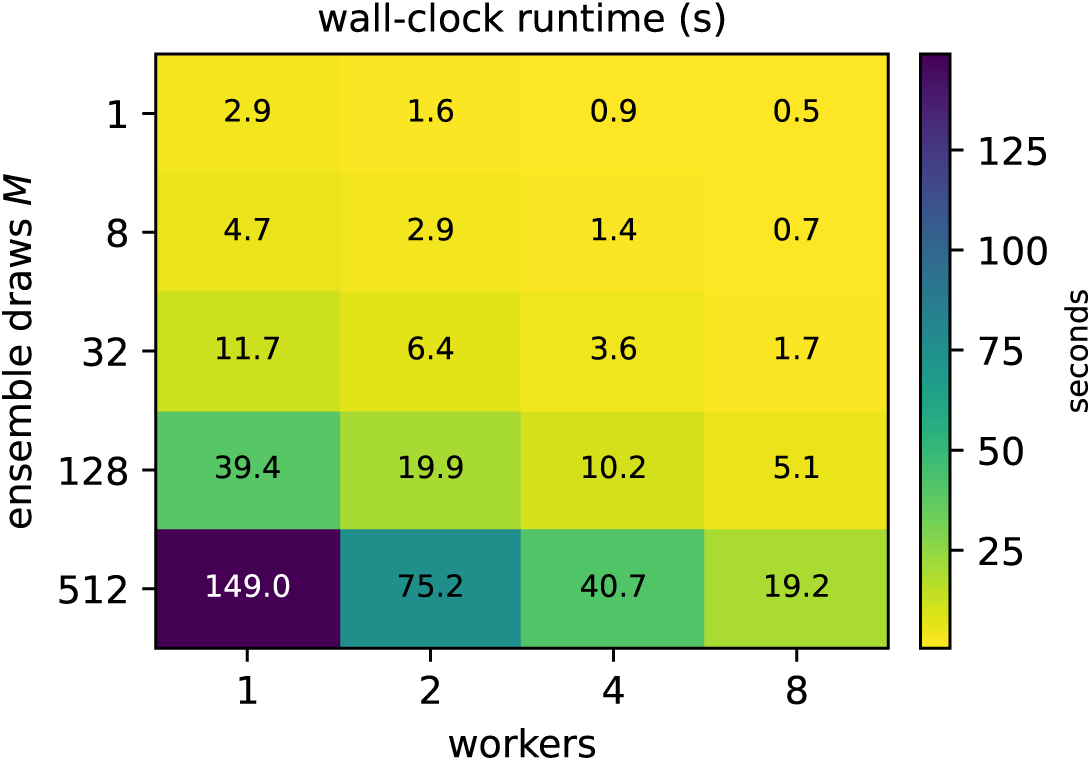
Local-tree runtime against ensemble size and number of workers. Time in seconds on 100 haplotypes over 2 Mb with 250 kb chunks.

**Figure C13:**
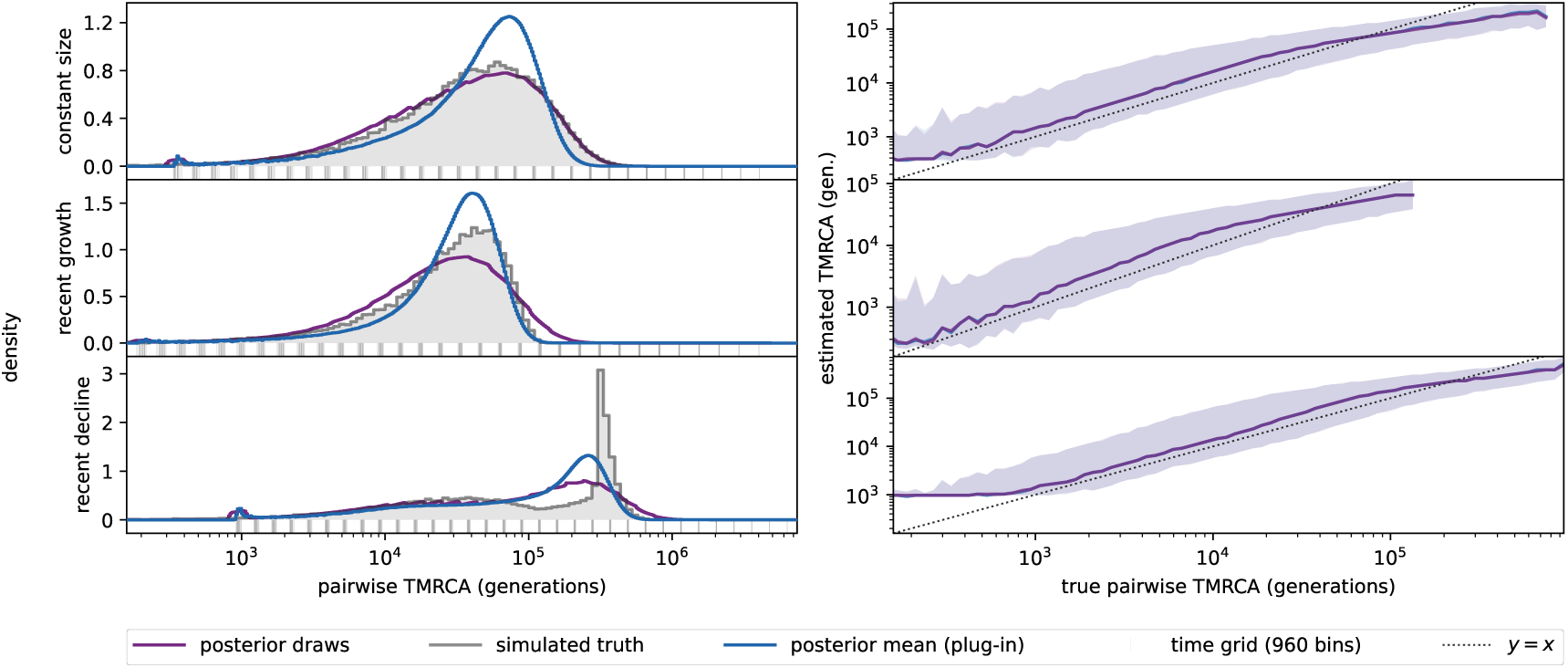
**Pairwise TMRCA recovery across demographies**, for a constant-size, a recently expanding and a recently contracting ingroup. **Left:** the marginal distribution of pairwise TMRCA under the simulated truth, the plug-in posterior mean and the ensemble draws, with the inference grid drawn as a rug. **Right:** each estimate against the truth, as the median and the central 90% over window–pair values. Collapsing the posterior to its mean narrows the marginal, which the draws restore.

### C.11 Window and Emission Block

The window and the HMM emission block are varied jointly in Figure C14.

**Figure C14:**
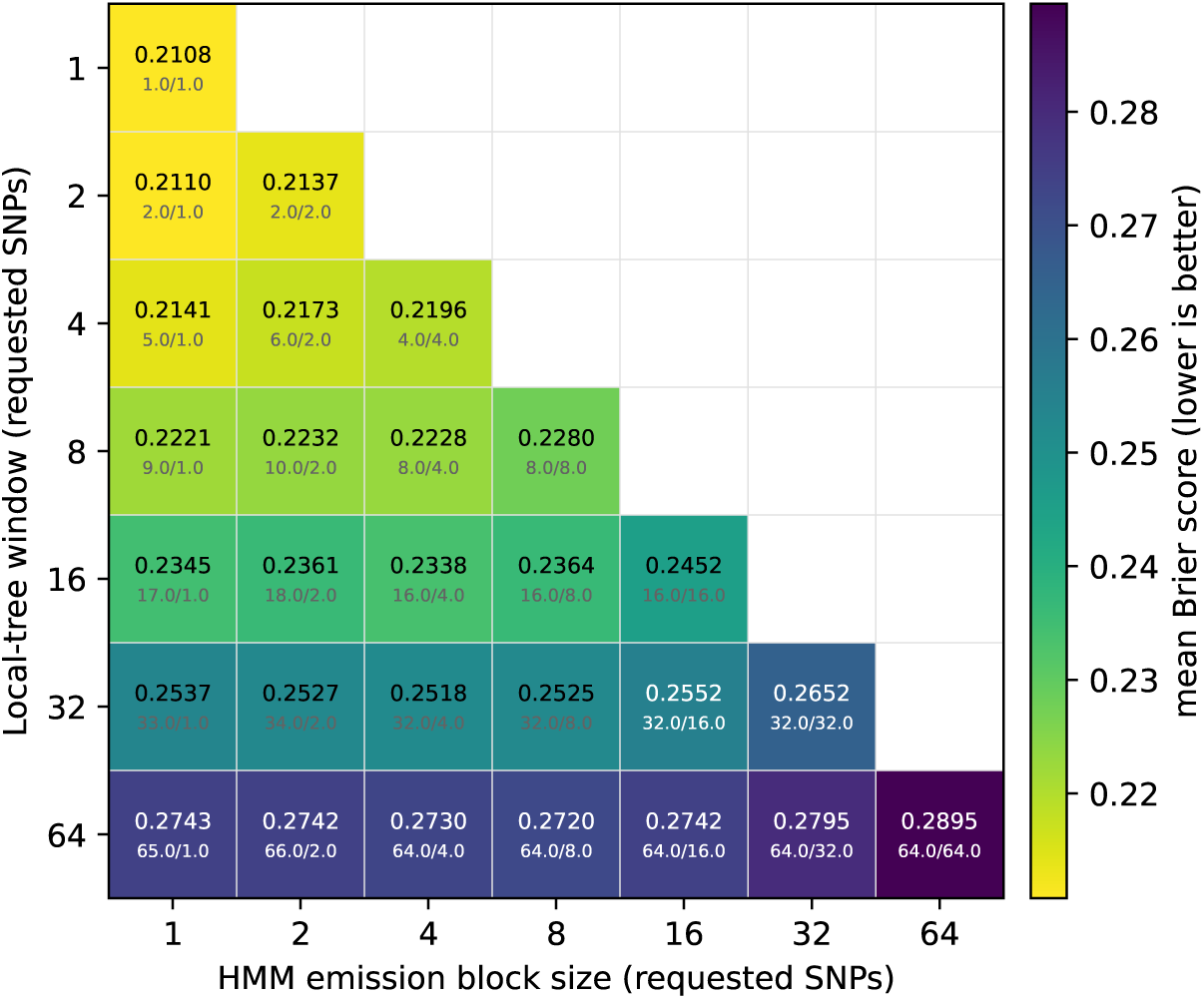
Ancestral-allele recovery over the window and the HMM emission block. Mean Brier score (lower is better) per cell, against the simulated truth on the same neutral ingroup-only `msprime` simulation as Figure 6 (*n* = 20 haplotypes, *L* = 10^8^ bp, *N_e_* = 3 × 10^4^); the true-ARG ceiling is 0.165. Cells are annotated with the score above the effective window and block in SNPs, which differ from the requested widths because a window is rounded up to a whole number of blocks. Only block ≤ window is shown, a wider block collapsing onto the diagonal. The window dominates, while the block matters through the number of blocks a window averages over, so recovery is worst on the diagonal where a window holds a single block.

### C.12 Outgroup Placement

To investigate optimal outgroup placement, Figure C15 varies the divergence of the deepest outgroup jointly with how the shallower outgroups are spaced beneath it and how many there are. Divergence times matter: recovery degrades where the outgroups are close enough to share the ingroup’s polymorphism, and again where they are distant enough for recurrent mutation to erode the signal, and is best over a broad range in between, of order *τ* ∼ 10, about 0.015 substitutions per site.

**Figure C15:**
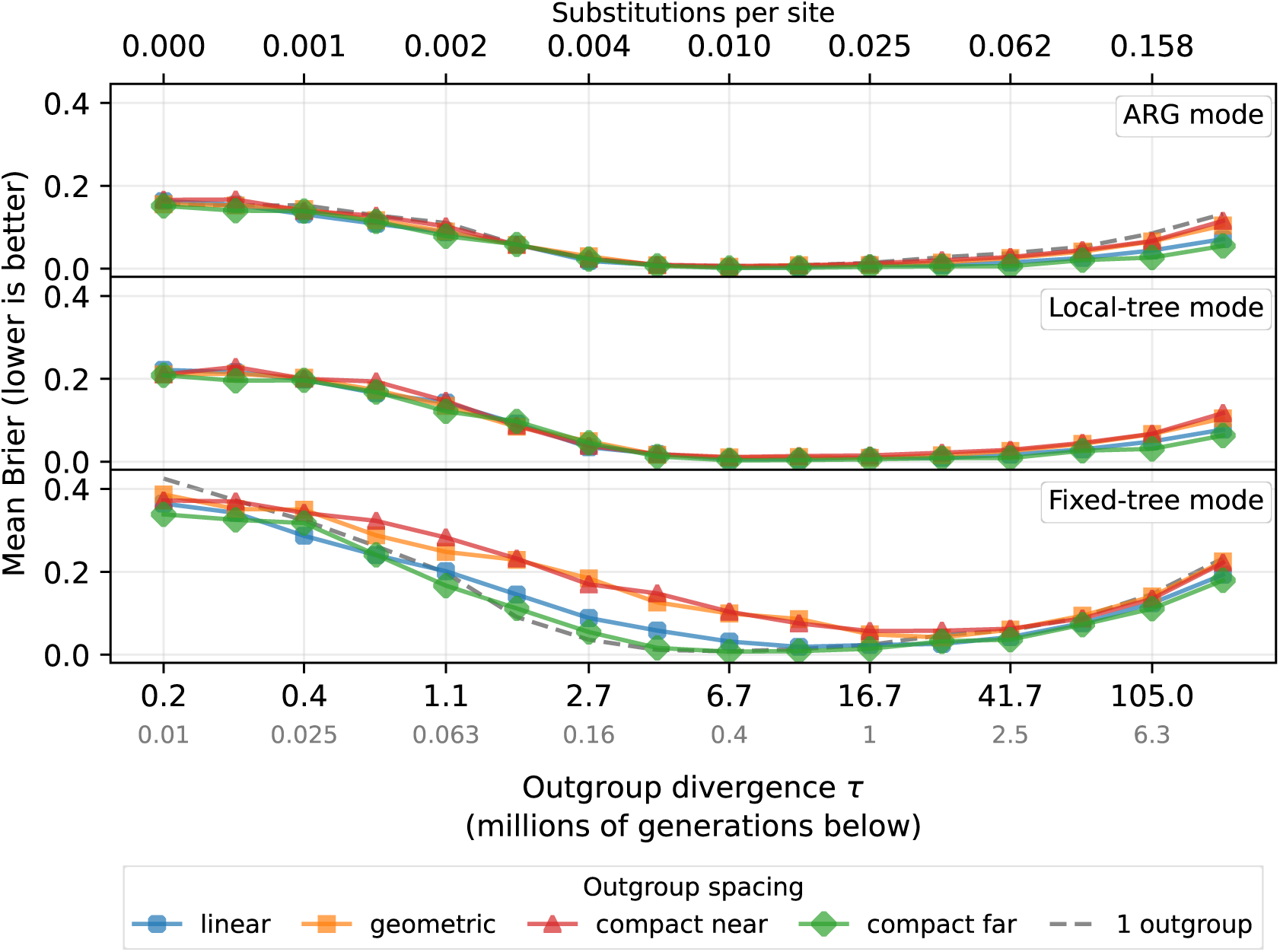
Ancestral-allele recovery against outgroup divergence. The solid lines carry three outgroups, one per outgroup spacing mode; the dashed line is a single outgroup. The mean Brier score is over the ingroup-polymorphic sites of a 1 Mb constant-size simulation with 20 ingroup haplotypes; depth is given in coalescent units *τ* = *T/*(2*N_e_*) with *N_e_* = 3 × 10^4^, so it is independent of *N_e_*, and the top axis converts it to expected divergence 2*Tµ* at *µ* = 1.25 × 10^−8^ per site per generation. Writing the deepest split time, the quantity on the horizontal axis, as *d*, the four outgroup spacings place the other two outgroups at (*d/*3, 2*d/*3) when even (*linear*), at (*d/*9*, d/*3) when geometric (*geometric*), and at (*d/*10*, d/*5) or (2*d/*3, 5*d/*6) when concentrated towards the ingroup (*compact near*) or towards the deepest outgroup (*compact far*). The deepest split is held at *d* throughout, so the four differ only in where the two shallower outgroups fall beneath it.

### C.13 Kernel Runtime

To decompose the per-tool runtimes by the implementation options the kernel exposes, we run all three modes on one shared 100 Mb `msprime` panel of 17 ingroup and 3 outgroup samples (778,783 sites, 250,657 trees), evaluating the same site set so that runtime and peak memory are directly comparable (Table C2). Each configuration runs in its own subprocess, isolating its peak memory. Additional workers benefit ARG mode substantially, though sub-linearly, fixed-tree mode parallelises only its multi-start fit and runs slower when each start is trivial, and local-tree mode scales with its kernel threads rather than the worker count. Peak memory, measured over the whole process group, therefore grows with ARG mode’s workers but stays flat as local-tree threads are added. Only local-tree mode scales quadratically with the panel size (Figure C16).

**Table C2:**
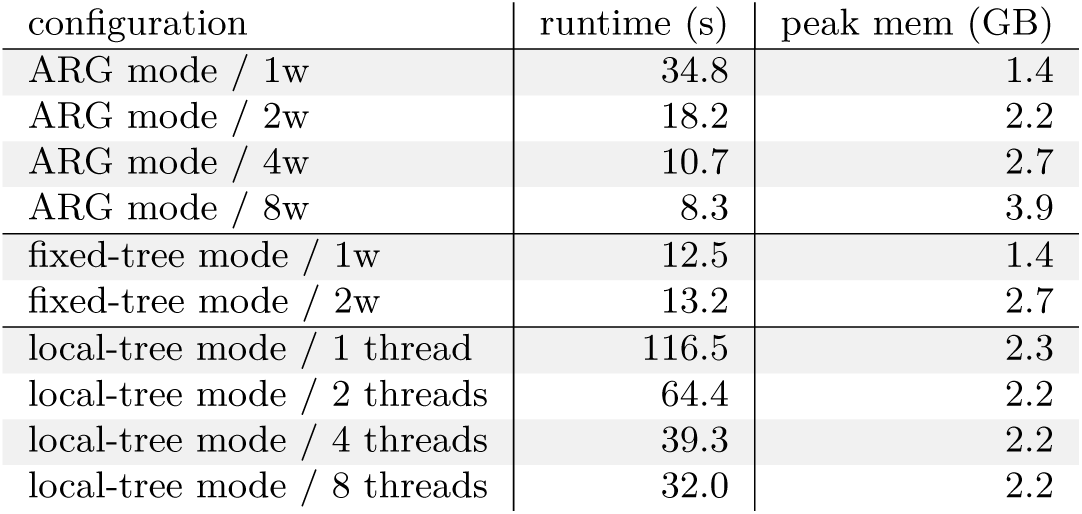
Kernel runtime and peak memory per configuration, on the shared panel described in the text.

| configuration | runtime (s) | peak mem (GB) |
| --- | --- | --- |
| ARG mode / 1w | 34.8 | 1.4 |
| ARG mode / 2w | 18.2 | 2.2 |
| ARG mode / 4w | 10.7 | 2.7 |
| ARG mode / 8w | 8.3 | 3.9 |
| fixed-tree mode / 1w | 12.5 | 1.4 |
| fixed-tree mode / 2w | 13.2 | 2.7 |
| local-tree mode / 1 thread | 116.5 | 2.3 |
| local-tree mode / 2 threads | 64.4 | 2.2 |
| local-tree mode / 4 threads | 39.3 | 2.2 |
| local-tree mode / 8 threads | 32.0 | 2.2 |

**Figure C16:**
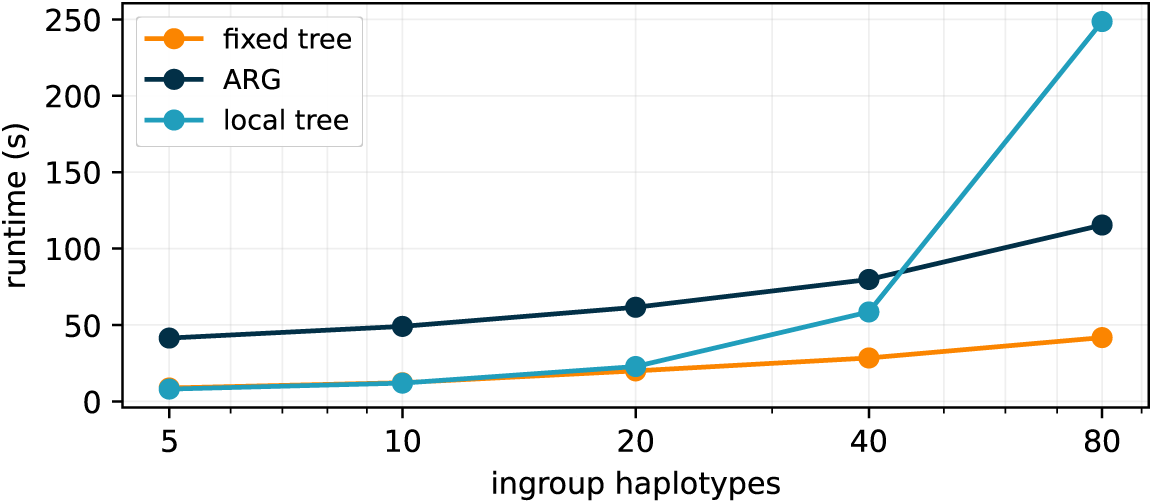
Runtime against ingroup size for the three modes. Time in seconds on a 100 Mb `msprime` panel with three outgroups, subsampled to each ingroup size, with local-tree mode at an ensemble of 32 draws.

